# Vertical climbing in mice: a translationally relevant measure of pain-related impact on natural behaviour

**DOI:** 10.64898/2026.09.30.755655

**Authors:** Morten E. Malmberg, Tobias Bro, Edward Ciputra, Jannie Jutta Jeppesen, Carolyn Marie Goddard, Ida Buur Kanneworff, Heather N. Allen, Helene Dragelund Garcia, Kathrine Louise Jensen, Yingyu Tang, Pernille Colding-Jørgensen, Kholud Walid El Masri, Andrea Soeberg, Astrid Thostrup Davidsen, Naomi K. Grabus, Karina Obelitz-Ryom, Anja Holm, Sarah Louise Christensen, Chelsea Hopkins, Anne-Marie Heegaard, Kenneth Lindegaard Madsen, Andreas Toft Sørensen, Rajesh Khanna, Klas S.P. Abelson, Otto Kalliokoski, Sara Hestehave

## Abstract

Despite urgent need for better analgesics, drug development has been complicated by incomplete bench-to-bedside translation. A likely contributing factor is the poor connection between the experimenter-evoked pain-related measures often used in preclinical models, and the clinical symptoms, like broader functional impairment, reported by patients. Here, we asked whether vertical climbing, a species-specific elective behaviour in mice, could serve as a measure of pain-related impairment. We tested male and female mice in a wide range of pain-related models including models of post-operative pain, inflammatory injury, osteoarthritis, rheumatoid arthritis, cancer-induced bone pain, fibrous dysplasia, neuropathic pain, migraine-related pain and facial neuropathy. Vertical climbing was reduced in all models, across five different laboratories, demonstrating the robustness and consistency of this type of behaviour as a novel pain-related outcome measure. Reduced climbing correlated significantly with established pain-related measures, but its time course diverged from that of evoked mechanical allodynia, while aligned more closely with other non-evoked behaviours, like burrowing, home cage activity, wheel running or weight bearing. Pharmacological validation was bidirectional: gabapentin restored climbing in mice with neuropathic injury, and naltrexone reinstated the climbing deficit in mice that had recovered from intraplantar CFA, consistent with persistent pain held in remission by endogenous opioid signalling. Overall, our results indicate that voluntary vertical climbing is a stable, elective behaviour that provides a robust and translationally relevant measure of pain-related functional impairment in mice.

**Summary:** Vertical climbing, an elective mouse behaviour, is reduced across diverse pain models, providing a robust, pharmacologically validated, translationally relevant measure of functional impairment.

## Introduction

Chronic pain conditions are among the leading causes of years lost to disability [1], yet effective treatments are scarce. Promising preclinical candidates routinely fail to translate into clinically effective therapies [41; 64]. Likely contributing factors include limited predictive validity of preclinical pain models and the poor ability of behavioural measures to quantify signs of pain in rodents [18; 51; 64; 65]. Pain is recognized as a personal and emotional experience [50], which is inherently challenging to assess confidently in a non-verbal rodent, and researchers must rely on behavioural indices as proxies[54]. Traditional assays typically measure reflexive withdrawal responses [54], which capture nociceptive sensitivity, but incompletely represent the broader functional impairment reported by patients living with pain[31]. Furthermore, these measures are usually experimenter-evoked, rendering them vulnerable to observer bias and inter-operator variability, as well as being modulated through experimenter-induced stress to the animals [54], even affected by the sex of the experimenter [59].

To improve translational relevance, two complementary strategies have emerged: i) paradigms that probe the subjective and affective experience of pain, such as conditioned place preference or avoidance [61] or facial grimace scales [35; 60], and ii) measures of spontaneous naturalistic behaviours that animals voluntarily perform under undisturbed conditions but are reduced during pain states. The latter class of endpoints offers several advantages over evoked measures: they more closely parallel the functional disability central to clinical pain, reduce confounders from experimenter interaction, and help distinguish true analgesia from sedation or general behavioural disruption, as motor-impairing drugs will suppress rather than restore behaviour [11; 46].

A range of such behaviours has been explored as translational endpoints, including nest building [21; 22], vocalisation [33], burrowing [4; 63], wheel running [11], and home-cage activity [24]. More recently, vertical exploration and climbing have emerged as ethologically relevant, elective behaviours in mice that reflect internal motivation, motor coordination, and engagement with the environment [55; 68]. Zhang et al. [68] identified cage-lid hanging as the home cage behaviour most reliably suppressed across a diverse panel of inflammatory, neuropathic, and visceral pain models. This suppression was, furthermore, reversible by clinically effective analgesics, although not by kappa-opioid agonists, a drug class with a poor record of clinical translation [68]. Santos et al. [55] subsequently validated voluntary climbing in a mesh-walled vertical cylinder, demonstrating that vertical climbing is stable across repeated testing, reduced by acute visceral pain, and selectively restored by an NSAID, but not by a kappa-opioid agonist [55].

Here, we developed a low-cost vertical climbing apparatus and evaluated its robustness across a variety of pain-related injuries and laboratory settings. In a collaborative, multi-group effort, we assessed vertical climbing behaviour across models of post-operative pain, inflammatory injury, osteoarthritis, rheumatoid arthritis, cancer-induced bone pain, fibrous dysplasia, neuropathic pain, migraine-relevant pain and facial neuropathy. We tested reproducibility across different mouse strains, experimenters and laboratories, and validated the assay against established behavioural endpoints and pharmacological interventions. Overall, our results indicate that voluntary vertical climbing is a stable, elective behaviour which may serve as a robust and translationally relevant measure of pain-related functional impairment in mice.

## 2. Methods

These experiments were performed using ten different models of pain-related conditions across different mouse strains and experimental facilities. Therefore, full methods descriptions of animal-, model- and facility-specifics are included in the **Supplementary file**, while a shortened summary is relayed here.

### 2.1. Animals and housing

Different mouse strains were used for individual model-specific experiments. Animals were always acclimatised for at least one week after arrival from the external vendor, before experiments were initiated, and housed in groups of 2-6, unless otherwise specified (Fibrous Dysplasia experiments involved single housing for automated home cage monitoring). Mice were housed in a temperature-controlled room, maintained on a 12/12-hour light/dark cycle with *ad libitum* food (rodent chow) and water, given housing- and nesting material, gnawing sticks and provided with additional enrichment when feasible. All experiments were performed in accordance with animal experimentation licenses granted by the Danish Animal Experiments Inspectorate, except the FRICT-ION experiments performed at University of Florida under local IACUC approval. For all experiments, model-specific humane endpoints were defined before study-start, and monitored throughout.

### 2.2. Model inductions and study designs

#### 2.2.1. Plantar incision model of post-operative pain

Male and female C57BL/6JRj were anaesthetised in an induction chamber using 5% isoflurane mixed in O_2_ and maintained via facemask at 1.9% during the surgery. The right paw was cleaned and disinfected, and the surgery was performed under aseptic conditions by an experienced surgeon (MEM) with an aseptic assistant (TB). The plantar incision was performed as described by Cowie et al.[14] with addition of aseptic measures. After preparation, the skin on the plantar surface of the paw was incised with a 4 mm longitudinal incision through the skin, and the flexor digitorum brevis muscle was lifted with curved forceps and split in half longitudinally, before returned to the wound cavity, and the skin was sutured with two interrupted sutures (Ethilon 6-0). Un-injured controls were exposed to anaesthesia and surgical disinfection of the paw.

The experiment was divided in sex-separated consecutive cohorts of 12 animals per cohort, with equal numbers from each experimental group. For the initial experiments, climbing behaviour was assessed at pre-injury baseline and approximately at 4 h, 24 h, 48 h, 74 h and 96 h after surgery, to characterise the impact of the model on climbing behaviour. A follow-up experiment was then conducted in only incised male mice, to test whether pre-surgically administered meloxicam (10 mg/kg, s.c.) affected climbing behaviour 4 h after injury. All behavioural tests with this model were conducted by the same male experimenter (TB), who was blinded to group-allocation.

#### 2.2.2. Inflammatory model - Intraplantar Complete Freund’s Adjuvant (CFA)

To model inflammatory pain, we used the intraplantar CFA model [48]. In brief, male and female C57BL/6Nrj mice were deeply anesthetised with isoflurane (2%) and received an intraplantar injection of CFA (7 µL, Sigma Aldrich, F5881) using a Hamilton syringe, and a single s.c. injection of buprenorphine (0.1 mg/kg) for post-procedure care. Control animals were exposed to anaesthesia and the same analgesic treatment, without intra-plantar injection.

The experiment was conducted in two cohorts of 12 animals per cohort, one cohort per sex. All six animals in a cage were allocated to the same treatment group after confirming comparable levels of climbing performance (duration climbing) between cages. Both cohorts were assessed on behaviour (vertical climbing and von Frey) at pre-injury baseline and on Days 2, 4, 7 and 11 after injury. For the female cohort, climbing behaviour was also recorded on Day 22, 1 hour after s.c. injection of naltrexone (NTX, 3 mg/kg) to re-establish pain-related responses. All behavioural tests with this model were conducted by the same female experimenter (CMG), who was blinded to group allocations.

#### 2.2.3. Osteoarthritis model - Monoiodoacetate-Induced Arthritis (MIA)

Induction of the MIA to the left knee joint was performed similarly to previously reported experiments [26–28]. Animals were anaesthetised using isoflurane mixed in O_2_ (induction: 2.5%, maintenance during injection: 2.0%) and shaved around the knee on the left hindleg. Using a 30 G insulin syringe, freshly prepared sodium iodoacetate (sigma, I2512, 1 mg/10μl) was injected intraarticularly into the joint space just below the patella. Control animals were anaesthetised and shaved around the knee but not injected. Model induction was performed by a female experimenter experienced with the model (SH).

The experiment was conducted on equal numbers of male and female C57BL/6JRj mice in one cohort including a total of 40 animals, divided into control or MIA groups (n = 8 or 12, respectively). Vertical climbing behaviour was assessed at pre-injury baseline, and then at Days 3, 7, 10, 14, 20, 28, 36 and 42. Additionally, Dynamic Weight Bearing (DWB) was assessed on Days 8/9, 15/16 and 29/30, but due to the high number of animals, this was measured over two days, one per sex. All behavioural tests with this model were conducted by the same female experimenter (HDG), who was blinded to group allocations.

#### 2.2.4. Rheumatoid arthritis model - Collagen-Antibody-Induced Arthritis (CAIA)

To model rheumatoid arthritis, we used the CAIA-model [5]. In brief, on Day 0, 1.5 mg (delivered in 0.15 mL sterile saline) mouse monoclonal antibodies against type II collagen (Arthrogen-CIA Arthritogenic Monoclonal Antibody 5-clone cocktail, Chondrex Inc.) was injected intravenously (i.v.) in the tail vein. The controls were injected i.v. with 0.15 mL saline. On Day 3, the CAIA groups received LPS (Lipopolysaccharide, Chondrex Inc.) from *E. coli* intraperitoneally (i.p.), with either 5 or 10 µg for the “low-LPS” vs “high-LPS” groups respectively. Controls were injected i.p. with saline.

The experiment was conducted in four consecutive sex-separated cohorts of BALB/cByJrj mice, comprising 12 animals per cohort, with four subjects from each of the three groups in each cohort (control, low-LPS and high-LPS, total n = 8 for each sex). Animals were housed in pairs of the same injury group. All cohorts were assessed on climbing behaviour at pre-induction baseline, and on Days 4, 8 and 14 after model induction. Additionally, burrowing behaviour was assessed on Days 6 and 13, and arthritis severity scores and animal welfare scores were monitored throughout. All behavioural tests were conducted jointly by two female experimenters (PCJ, KWEM), who were both blinded to group allocations.

#### 2.2.5. Cancer Induced Bone Pain - (induction)

Cancer-induced bone pain models were induced via intrafemoral inoculation of cancer cells into right hind legs as described before [16; 17; 30]. Specific details on cell lines, cultures, and strains can be found in the supplementary materials. Briefly, mice were initially anesthetised with a mixture of 43 mg/kg ketamine and 6 mg/kg xylazine i.p., and given 5 mg/kg carprofen (s.c.) for pre-surgical analgesia and 0.5 mL of 0.9 % saline (s.c) to prevent dehydration. Mice were maintained under 2 % isoflurane and kept on a heating table at 37 °C throughout the surgery.

The right hind leg was shaved and disinfected, and a small, horizontal incision was made over the patella tendon. The lateral side of the patella tendon was loosened, and the patella tendon was laterally displaced, allowing access to distal femoral epiphysis. A hole was then made in the femoral epiphysis into the intramedullary femoral cavity with a 30 G needle. Needle placement was verified via X-ray. Using a 0.3 mL insulin syringe, 10 µL of either vehicle (HBSS) or cell suspension was then inoculated into the cavity. The hole was sealed with bone wax (Ethicon, US), the patella tendon was repositioned, and the incision was closed with surgical clips. Post-operative analgesia was maintained with 5 mg/kg carprofen (s.c) on Days 1 and 2. Mice were subjected to daily severity assessment and wound monitoring for four days, and medical clips were removed at the end of day 4.

##### 2.2.5.1. Breast-cancer-induced bone pain

Eighteen female BALB/cOlaHSD mice were obtained at 6-7 weeks of age, inoculated with tumour cells two weeks later, and subjected to behavioural assessment starting from 9-10 weeks of age. Baseline assessment of static weight bearing, vertical climbing and X-ray imaging was performed before tumour inoculation. Mice were randomised based on their body weight and weight bearing baseline ratios into sham (n = 9) and cancer (n = 9) groups. Mice were then inoculated with 10,000 cells/10 µL of 4T1-Luc2 cells (or HBSS for the sham group) at Day 0, followed by post-surgical care and monitoring from Day 1-4. From Day 6-14, mice were subjected to limb use and weight bearing assessment every day, and vertical climbing every second day until reaching a humane or experimental endpoint. To verify disease burden, mice were X-ray imaged again at the terminal end-point to determine the relative bone density of the proximal femur as described before [30].

All behavioural assessments and tumour cell inoculations were performed by one male experimenter (EC), who was blinded throughout to the experimental group allocations.

##### 2.2.5.2. Multiple Myeloma (MM) induced bone pain

Eighteen male C57BL/KaLwRijHsd mice were obtained at 4-5 weeks of age, inoculated with tumour cells two weeks later, and subjected to behavioural assessments starting from 8-9 weeks of age. Baseline assessment of vertical climbing was recorded, and mice were allocated into sham (n = 8) or cancer (n = 10) groups, based on their body weight. Mice were inoculated with 50,000 cells/10 µL of 5TGM1-GFP cells (or HBSS for the sham group) at Day 0, followed by post-surgical care and monitoring from Days 1-4. Starting from Day 8, mice were subjected to behaviour assessment. Limb use was assessed on Days 8, 11, 15, 17, 19, 21, and every day from Day 23 until reaching a humane or experimental endpoint. Vertical climbing was assessed at Day 8, 11, 15, 19, 21, and every other day from Day 23. To verify the disease burden, the spleen was collected from each mouse and weighed.

Limb use assessments and tumour cell inoculations were performed by one male experimenter (EC), while vertical climbing was recorded by a female experimenter (AD). Both experimenters were blinded throughout the experiment to experimental group allocations.

#### 2.2.6. Fibrous Dysplasia (FD)

The transgenic mouse model of FD used in this study was based on a previously reported *Prrx1-Cre*–driven, doxycycline-inducible *GαsR201C* transgenic system [29; 69]. Experimental mice carried the *Prrx1-Cre* and *ROSA26-rtTA* Linker alleles, with or without the *Tet-GαsR201C* transgene. Mice positive for *Prrx1-Cre*, Linker, and *GαsR201C* were classified as fibrous dysplasia mice, whereas littermate mice carrying *Prrx1-Cre* and Linker but lacking *GαsR201C* were used as controls. Genotyping was performed prior to group allocation.

Mice were single-housed in cages (DVCs; Tecniplast) equipped with electromagnetic wheels (GYM500, Tecniplast) where locomotion and wheel running distance was automatically tracked throughout the study [32]. For analysis, only data generated during the dark phase was included.

To induce the model, doxycycline (doxycycline hyclate; Sigma-Aldrich) was administered in drinking water (0.08-0.1 g/L) throughout the experiment for 7 control and 5 FD male mice of approximately 13-14 weeks of age. Vertical climbing was assessed at baseline, and Days 6 and 13 after initiation of treatment with doxycycline. X-ray images of hind limbs were obtained under brief 2.5% isoflurane anaesthesia at baseline, as well as Days 8, 15 and 22 after injury, for scoring of disease burden. X-ray imaging and doxycycline administrations were performed by a female experimenter (YT), while vertical climbing was recorded by another female experimenter (AS). As all behavioural scoring was automated, the experimenters were not blinded.

#### 2.2.7. Neuropathic pain model - Spared Nerve Injury (SNI)

To model neuropathic pain, the SNI model was induced in C57BL/6Nrj mice [33]. In brief, mice were deeply anesthetised with isoflurane and injected s.c. with buprenorphine (0.1 mg/kg, 0.03 mg/mL) for peri-operative care. A small incision was made in the skin of the left hind leg between the hip and the knee. A few drops of a cocktail of lidocaine (2.5 mg/mL) and bupivacaine (1.25 mg/mL) were applied topically to the muscle for 30 seconds before being swabbed away. Using blunt dissection, the muscles were separated to expose the sciatic nerve branches. A single surgical knot was tied around the tibial and peroneal nerve branches, leaving the sural nerve intact, before cutting out a small piece of the tibial and peroneal nerve branches. The surgical wound was closed with two medical clips and for post-operative care, mice were injected with carprofen (5 mg/kg, 0.5 mg/mL) s.c. immediately after surgery. Carprofen injections were repeated the two following days, and surgical clips were removed on Day 3-4 post-surgery.

Two different SNI experiments were conducted, both including C57BL/6Nrj mice (n = 10-12). One study included males only, and with the controls exposed to neither anaesthesia nor sham injury. Another study was conducted with both males and females, and with a sham injury consisting of anaesthesia, incision to the skin, medical clips and the same peri-operative care as described for the SNI procedure. For both studies, pre-injury baseline climbing behaviour was assessed, followed by repeated assessments after SNI (male-only: Day 3, 7 and 12. Male-female study: day 4, 9 and 16) to characterize the model-inflicted impairment on vertical climbing behaviour. For the male-female study, a cross-over study exploring the effects of gabapentin (30 mg/kg) vs vehicle, 1 h after injection, was performed on Days 27 and 30 after injury, including equal numbers of animals from each experimental treatment on each day, and a 3-day washout between. Behavioural tests for the male-only study were performed by the same female experimenter (KLJ), while another female experimenter performed the male-female study (IBK). As all behavioural scoring was automated, the experimenters were not blinded.

#### 2.2.8. Levcromakalim-induced migraine model (LEV)

Levcromakalim-induced migraine was established in C57BL/6JBomTac mice by injection of levcromakalim (1 mg/kg, i.p.) every other day; Days 0, 2, and 4 [9]. Levcromakalim (Tocris Bioscience, Bristol, UK, batch no: 1A/275955) was dissolved in saline with 2 % DMSO to a final concentration of 0.01 mg/ml, which was injected in a volume of 10 ml/kg using a 27 G needle.

Baseline vertical climbing and mouse grimace scores (MGS) were obtained the day before first experimental assessment (day -1). The vertical climbing test was performed 1 and 2 hours post injection of levcromakalim on test Days 0 and 4. On test Day 2, MGS were recorded 1 h after levcromakalim injection. Vaginal lavage samples were taken from female mice after behavioural testing on all test days for oestrous cycle stage determination.

Mice were allocated to vehicle or levcromakalim groups (n = 8-9), stratified by baseline climbing, sex, and home cage. All behavioural experiments were performed by the same female experimenter (JJJ). As all behavioural scoring was automated, the experimenter was not blinded to experimental group.

#### 2.2.9. Orofacial neuropathic pain - Foramen Rotundum Inflammatory Constriction Trigeminal Infraorbital Nerve injury (FRICT-ION)

FRICT-ION surgery was performed on male and female C57BL/6J mice as described [44]. Mice were anesthetized with 5 % isoflurane and maintained on 2.5 % isoflurane throughout the procedure. A small incision was made in the inner buccal mucosa to expose the maxillary branch of the right infraorbital nerve near the foramen rontundum. A 3 mm segment of 4-0 chromic gut suture (Ethicon #724G) was inserted along the nerve into the foramen rotundum. Mice recovered on a heating pad until fully ambulatory and were returned to their home cage. Animals were monitored daily for any changes in body weight, healing, and infection.

The vertical climbing test was performed at baseline before surgery. Animals were also tested for facial mechanical (von Frey) and cold (acetone droplet assay) hypersensitivity prior to surgery. Mice were divided into groups (sham or FRICT-ION, n = 7-8) based on sex and baseline climbing behaviour. Following surgery, mice underwent mechanical and cold facial sensitivity testing and the vertical climbing test every seven days for three weeks. Evoked testing and the vertical climbing test were performed on separate days. The experimenter was blinded to surgical condition (sham or FRICT-ION), and all quantification was performed blinded.

### 2.3 : Behavioural testing and other assays

All the behavioural assessments were performed in the daytime, except for the automated home cage activity measures, which were recorded during the night. Blinding to experimental group was always implemented if observer-scored, while not necessarily if an experiment only included automated assessments. All study designs are shown schematically in **Supplementary Fig S1-10**.

#### 2.3.1. Vertical Climbing

Our vertical climbing setup was based on the previously published device from Santos et al. [55]. The aim was an easy-to-build apparatus, to be made freely available for anyone who would wish to employ it in their experiments. See the full construction-manual and Standard Operating Procedure (SOP) on the open protocol repository, protocols.io: https://dx.doi.org/10.17504/protocols.io.n92ldo7q7g5b/v1.

The vertical climbing assay consisted of an enclosed vertical cylinder with a grid mesh made of galvanized steel (climbable height: 250 mm, diameter: 150 mm, mesh size: 6 mm x 6 mm), standing on a 3D-printed platform made of either black or grey Polyethylene Terephthalate Glycol (PETG). For testing, animals were carefully placed on the inside of the mesh cylinder, and the mesh was carefully placed vertically in the stand. A camera was placed in front of the apparatus from approximately mid-height of the cylinder to capture the full cylinder, including its base, to record the animal’s behaviour in the cylinder (see **Fig 1**).

**Fig 1:**
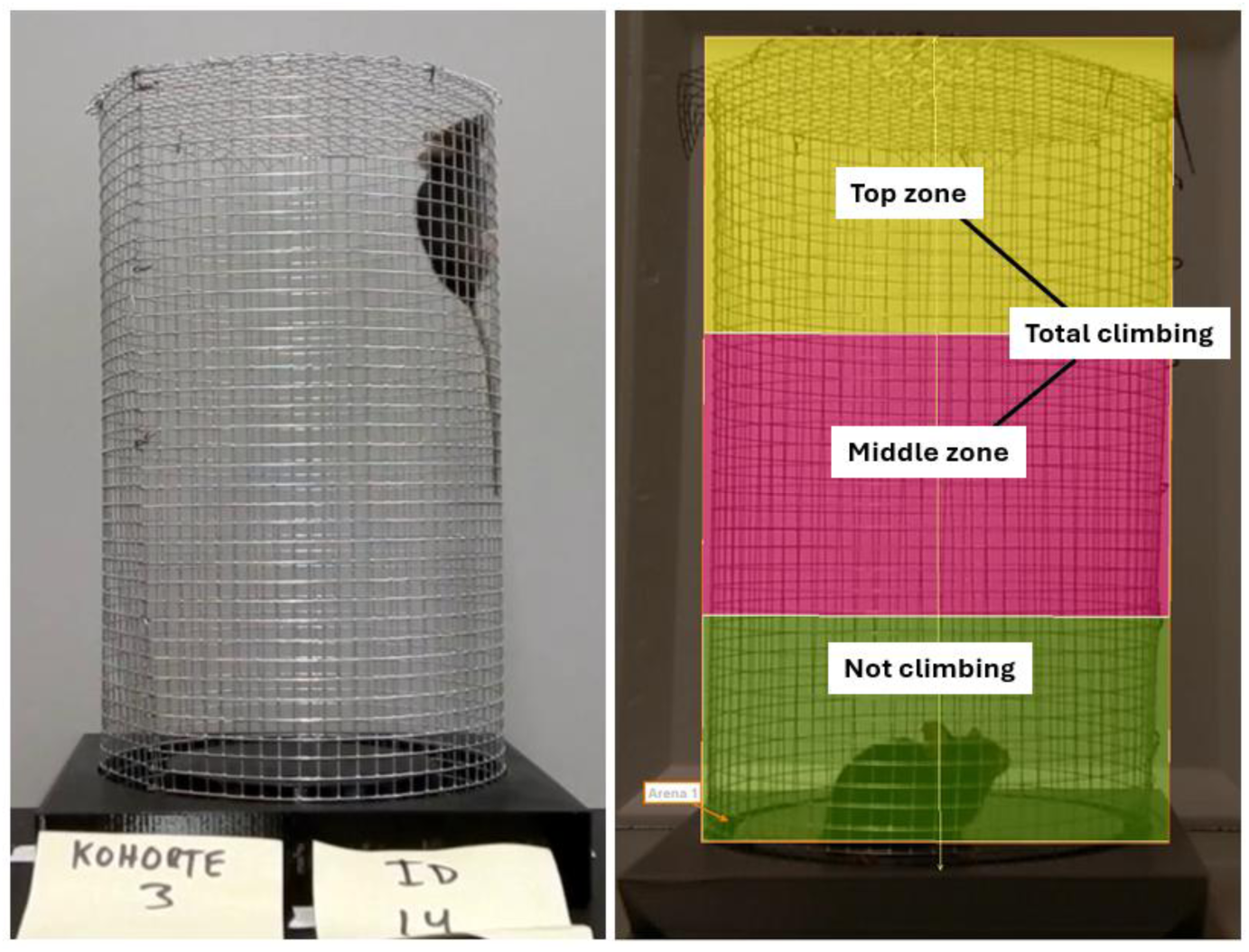
Left: example of a C57BL/6J mouse climbing the vertical climbing cylinder. Right picture displays the Ethovision arena-settings used for tracking. Coloured areas are coded as not climbing (green), middle climbing (red), top climbing (yellow). The total climbing was recorded as middle and top zones combined.

The protocol for the climbing test consisted of a 2-day habituation/training phase, followed by a baseline test before model induction and subsequent testing.

- <u>Day 1 - group habituation,</u> where mice were placed in the cylinder as a group, meaning all mice from a single home cage were placed in the same grid and left to climb together for 15-30 min. This provided minimally stressful conditions for the mice to explore the cylinder with their cagemates.

- Day 2 - single habituation: the mice were placed in the cylinder individually and left to climb for 10-15 min to ascertain that all mice would climb freely in the grid.

- <u>Day 3 - baseline test:</u> each mouse was placed individually in the cylinder, and behaviour was recorded for 11-12 min. For analysis, the first minute in the cylinder was always excluded, to allow the experimenter to move away, and the animal to settle down. After the first minute, the following 10 minutes were used for analysis and tracked either live or retrospectively using Ethovision XT (Noldus technology, NL).

- Test days after model induction: The timing of the following test-days depended on the pain-related model and experimental hypothesis, but the protocol followed the same steps as the baseline test above.

Between tests, the platform was cleaned with 70 % ethanol, to reduce the risk of odour cues affecting the behaviour of the following animal. Often, two vertical climbing cylinders were used to allow two animals to be tested simultaneously. Under such circumstances, great care was taken to place the setups as far away from each other as possible, and in a manner ensuring that a mouse in one grid could not see the mouse in the other grid, using dividers or similar.

To ascertain appropriate levels of natural climbing behaviour, exclusion criteria were implemented. The exclusion criteria were applied to the “Total climbing” (collecting time in top + middle zone) duration at baseline, and a given mouse was excluded from the climbing study, if it spent either 1) more than 570 seconds, or 2) less than 30 seconds climbing, out of the full 600 seconds (10 min). This constitutes 95% of the time tested, where the mouse was either climbing or not-climbing, respectively. The rationale being that the assay was built to quantify voluntary climbing activity, and conditions which modify this activity. These goals would be undermined if, 1) the animal did not consider the base as a safe place to relax / avoid climbing, or 2) if the animal had no natural desire to climb under normal circumstances. Under such circumstances mice were considered non-compliant and excluded. When possible, baseline analysis was made before animals were allocated into groups, for exclusion of extreme- and non-climbers.

Video recordings were analysed using the Ethovision XT 15-17 software for detection of the centre point of the animal when moving around in the cylinder. After discarding the first minute, 10minutes of video recording was analysed, starting from minute 1 to minute 11 using a dedicated arena-template including a lower “non-climbing”-section, “middle climbing” and “top-climbing” sections, as shown in **Fig 1**. As each lab used their own cameras and behaviour rooms, the tracking template will have led to slight differences between each lab, depending on the camera angles and as more or less of the floor or ceiling could be seen in a given video frame. This may also have given some inaccuracies in detecting the distance moved when climbing the front or backside of the metal cylinder. Additionally, for each experimental setup, it was checked that when a given animal was standing on its hindlegs, the centre point did not move into the climbing zone. Importantly though, within each experiment, tracking and arena-settings were kept consistent, making comparisons between groups not affected by these small inaccuracies. Vertical climbing duration (in top + middle zone), frequency in top, and distance moved in the full cylinder, were retrieved and further processed for analysis.

#### 2.3.2. Mechanical allodynia, von Frey (CFA study)

Animals were habituated to red-tinted polyvinyl chloride (PVC) cups on a plastic grid for 10-15 min prior to assessment of mechanical sensitivity through the ascending method [15; 56], where the lowest non-responsive filament was tested on all animals before continuing to the next largest filament. Three out of five applications were recorded as a positive response. Both ipsilateral and contralateral paws were tested as internal control.

#### 2.3.3. Facial allodynia, von Frey and acetone droplet tests (FRICT-ION study)

Animals were habituated to light restraint (Plexiglas tubes, with their tails fixed in place) for three days before behavioural testing was performed. On the test day, mice were placed in light restraint tubes for at least one hour before testing began. Von Frey filaments were used to assess facial mechanical hypersensitivity by probing the face directly behind the whisker pad using the up/down method. Facial cold allodynia was assessed using the acetone droplet test. A 3 mL syringe attached to PE10 tubing with a flared end was used to place a 10 µL droplet of acetone on the animal’s face directly posterior to the whisker pad. The time spent reacting (grooming, face wiping, etc.) was recorded in seconds.

#### 2.3.4. Dynamic Weight Bearing (MIA study)

Dynamic weight bearing (DWB) was assessed in the MIA model, using the Dynamic Weight Bearing 2.0 apparatus (Bioseb, Vitrolles, France), as before [6]. The mice were tested across two days, one sex per day, to minimize scent of the opposite sex. Mice were individually placed in a Plexiglas chamber (11 x 11 x 20 cm) on a sensor pad detecting pressure and monitored by a camera in the lid detecting posture. Mice were allowed to first habituate to the device for 5 min, before capturing pressure data and live recording for 5 min of free movement within the chamber. The equipment was cleaned with 70% ethanol between each subject.

Acquisitions were next manually validated, consisting of verifying the placement of the animal’s limbs by comparing live recordings with a scaled map of the activated pressure sensors of the floor sensor pad. Sequences where the experimenter was uncertain about the correct pairing of limbs and pressure readouts from other body parts, such as the tail, were excluded. A minimum of 1 min of validated testing was used to calculate weight bearing characteristics for each mouse [52; 53]. Of the many values computed by the program following validation, only the DWB values of the hindlimbs were used. DWB values were obtained as the percentage of body weight (%BW) distributed on the ipsilateral leg during immobility.

#### 2.3.5. Arthritis score (CAIA study)

For the CAIA studies, clinical signs of active joint inflammation were assessed daily using the previously published methods by Nandakumar et al. [45] In short, each animal was assessed for visual signs of arthritis - redness and swelling - by clinical paw scoring. Each animal was gently restrained, and inflammation was assessed on all limbs, giving each inflamed toe or knuckle one point, while an inflamed wrist or ankle was awarded five points. Thereby, each limb could receive a score from 0 to 15 (five toes + five knuckles + one wrist/ankle), totalling 0 to 60 points for each mouse (**Supplementary Fig S11**). Thereby, the higher the score, the more widespread signs of disease activity.

#### 2.3.6. Burrowing behaviour (CAIA study)

Burrowing behaviour was assessed as an ethologically relevant, species-typical behaviour [57; 63]. The test was conducted as previously described [57], although with minor modifications. Each burrowing tube was made of PVC (length: 210 mm, diameter: 750 mm, frontal-end: raised and open, back-end: closed, **Supplementary Fig S12**) and was placed in a standard transparent plastic cage (Macrolone Type 3) without bedding. Mice underwent a 3-day habituation protocol as follows: Day 1: 1 h habituation in co-housed pairs to the cage with empty burrowing tube. Day 2: 2 h habituation in co-housed pairs to the cage with a tunnel containing 100 g of bedding material (wood shavings, Brogaarden). Day 3: 2 h single habituation to the cage with tunnel including 100 g of material. After the three days of habituation, two baseline tests occurred, where each animal was allowed to perform burrowing behaviour for 2 h, after which the bedding material remaining in the burrowing tube was measured, and the amount displaced was used as a measure of burrowing activity. The same approach was used for the post-induction test days.

#### 2.3.7. Limb use score (Bone Pain studies)

During each assessment session, mice were first habituated with their cage mates in a standard transparent cage measuring 125 x 266 x 185 mm (Tecniplast, Italy) for ∼10 min. Afterwards, mice were placed, individually, in a similar cage where they were videorecorded for 3 min. A blinded observer then evaluated the mice’s gait and scored their limb use. For the breast cancer bone pain model, mice were scored as follows: 4 – normal gait, 3 – insignificant limping, 2 – significant limping and shift in bodyweight towards contralateral limb, 1-significant limping and partial lack of use of the ipsilateral limb, 0 – total lack of use of the ipsilateral limb[30]. For multiple myeloma mice, mice were scored as follows: 4 – normal gait, 3 – insignificant limping, 2 - significant limping and shift in bodyweight distribution towards contralateral limb, 1 – significant limping and motor impairment, 0 – paraplegia [16].

#### 2.3.8. Static Weight Bearing (Bone Pain studies)

Incapacitance Tester Version 5.2 (Linton Instrumentation, UK) was used to assess the static weight bearing on the hind legs[30]. Before baseline assessment, mice were habituated and trained to rear for a minimum of 3 seconds with each hind paw placed on its own weight-sensing plate. For each assessment session, weight bearing was recorded for 3 seconds in triplicates for each mouse. The distribution of weight on the hind paws was determined with the following equation: % weight bearing = weight on ipsilateral limb/(weight on ipsilateral limb + weight on contralateral limb) (total weight on hind limbs).

#### 2.3.9. Home cage activity and wheel running (FD study)

Mice were single housed in cages (DVCs; Tecniplast) equipped with electromagnetic wheels (GYM500, Tecniplast) where mouse locomotion activity and wheel running distance are automatically tracked[32]. For analysis, data generated during the dark phase were included.

#### 2.3.10. X-ray imaging and bone deformity score (FD study)

In the Fibrous Dysplasia study, X-ray imaging was conducted weekly using the Lumina XR IVIS (In Vivo Imaging System) apparatus (Caliper Life Sciences, Teralfene). Mice were sedated and maintained using 2.5 % isoflurane for 5-10 min. X-ray images were obtained of the entire skeleton and analysed to determine FD burden of each individual animal at different time points (**Supplementary Fig S13**). The primary focus was to follow the progression and to investigate a possible correlation between disease burden and performance in the climbing assay. An ordinal scale was defined in advance with 0 - 3 points depending on the lesion severity in each area of interest, and the scores were added up for each animal (**Supplementary Table S2**). The higher scores suggest higher disease burden.

#### 2.3.11. Mouse grimace scoring (LEV-induced migraine model)

For grimace scoring, mice were placed in a white cardboard espresso cup held in a horizontal position at the edge of a table using a custom-made device (Supplementary **Fig S14**) placed in front of a video camera (LEGRIA HF G70, Canon Inc., Japan). To ensure sufficient light to separate facial features in the black coated mice, recordings were made at 1900-2000 lux illumination.

Mice were habituated to the espresso cups for 30 min a few days before baseline measurement. Furthermore, mice were acclimatized to the set-up for 15 min immediately before recording the 11-minute test for scoring. Videos were scored using PainFace, an open-source machine learning tool developed specifically for mouse grimace scoring of black coated mice (https://painface.net[38]) providing mean total grimace scores between 0 and 8.

#### 2.3.12. Determination of oestrus cycle (LEV-induced migraine model)

Female mice were subjected to vaginal lavage after behavioural testing at baseline and Days 0 and 4 in the LEV experiment following procedures adapted from Cora et al. [12]. A drop of the collected vaginal fluid was smeared in an even thin layer on a glass slide, stained by 0.1% toluidine blue, and the sample was categorised into one of the four stages of the mouse oestrous cycle: prooestrus, oestrus, dioestrus, and metoestrus depending on the presence of cell morphologies and cellularity [12] (Supplementary **Fig S15**). Two individual assessors performed independent scoring of the samples (JJJ & SLC), and a third assessor (AH) clarified discrepancies between the verdicts. The final verdict was the score determined by two of the three assessors.

### 2.4. Statistical analysis

Statistical analysis was performed using GraphPad Prism version 11 (GraphPad Software, Inc., La Jolla, CA, USA) or SPSS version 24 (IBM) for time course datasets where baseline recordings were included as covariates (RM ANCOVA). For data with randomly missing values (due to errors in for instance video-collection of behaviour), mixed-effects models were used. For *post hoc* comparisons between injured and control animals at individual timepoints in time course datasets, Estimated Marginal Means (EMM) were calculated with Bonferroni correction. Full analysis results and approach are displayed in the **Supplementary Table S3**. *P* < 0.05 was considered statistically significant. Error bars in the graphs represent mean ± SEM, unless otherwise specified. Qualitative scores, like limb use and bone deformity scores were presented purely descriptively and not analysed statistically.

For cancer bone pain experiments, the experimental endpoint was considered the humane endpoint, where an animal scored at least 6 on the severity risk assessment or scored a zero on limb use. As is the norm in this field, once that point was reached, the animals’ last measure was carried forward to the following timepoints, as the Last Observation Carried Forward (LOCF) approach, thereby avoiding the misconception that the group was “recovering” as the most severely affected animals were removed from the dataset.

## 3. Results

All statistical results and analysis details are presented in full in the **Supplementary Table S3**, while only summarized here.

### 3.1. Acute postoperative pain impairs vertical climbing

We first asked whether acute postoperative pain reduces vertical climbing, using the plantar incision model [14]. Male and female C57BL/6JRj mice received plantar incision or served as controls, and were followed for 5 days (**Fig 2**). Data were analysed with a mixed-effects model (sex * injury * time, with baseline as covariate; full results in **Supplementary Table S3**). Across the full 5-day time course, incision produced only a trend for effects of injury (P = 0.059) on the distance moved in the climbing cylinder (**Fig 2-A1**), but when the analysis was restricted to the acute phase at 4 h, the effect was significant across sex (**Fig 2-A2**). The post-operative injury produced more prominent changes in the frequency of climbing to the top of the cylinder (**Fig 2-B**) and the total duration spent climbing (**Fig 2-C**), where significant effects of injury were detected for both the full-time course data sets, as well as when focusing on the 4 h timepoint. None of the three outcome measures showed any statistically significant effects of sex or sex*injury interaction, suggesting that the general vertical climbing behaviour was comparable for males and females, and that their vertical climbing behaviour was affected equally.

**Fig 2:**
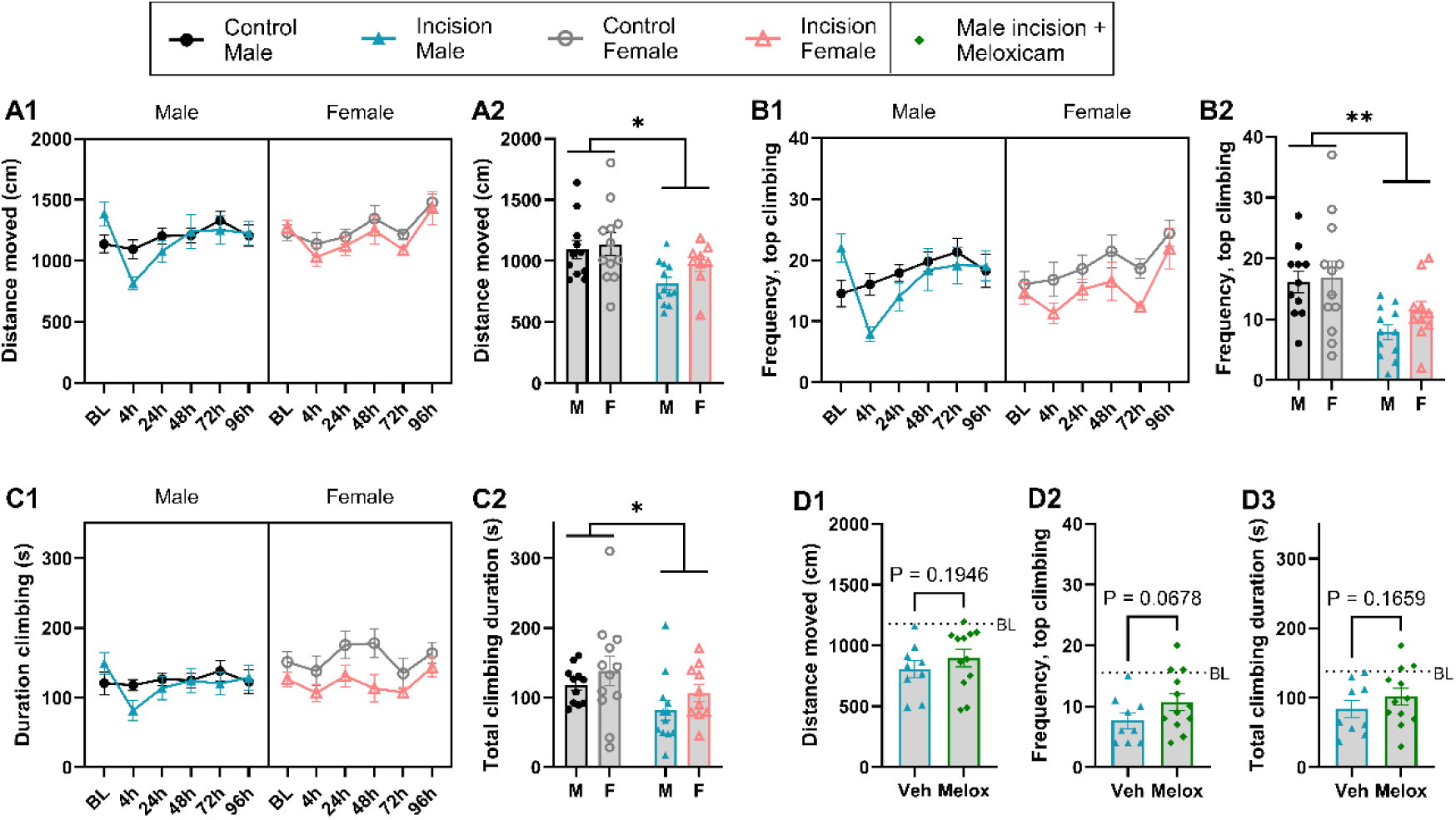
Acute postoperative pain impairs vertical climbing. Plantar incision was induced under isoflurane anaesthesia in male and female C57BL/6JRj mice. Vertical climbing behaviour was recorded for 10 min at baseline (BL) and at 4 h, 24 h, 48 h, 72 h, and 96 h after injury, and behaviour was quantified through automated video-tracking. (**A1**) Time course development of distance moved (cm) during the study and (**A2**) at the 4 h timepoint. (**B1**) Time course development of frequency in reaching the top of the cylinder during the study, and (**B2**) in focus at 4 h. (**C1**) Time course development of total duration climbing (s) during the study, and (**C2**) at 4h. In another cohort of male mice, all animals were exposed to plantar incision injury, and meloxicam (10 mg/kg) or vehicle was provided at the time of surgery. At 4 h after surgery vertical climbing was assessed through **(D1)** distance moved, **(D2)** frequency in the top and **(D3)** total duration climbing. Dotted line suggests pre-injury average baseline. A2, B2, C2: *: P<0.05, **: P<0.01, as suggested by injury-effect in a 2-way ANOVA (factors: sex & injury). D1-3: p-values suggested by unpaired t-tests. n = 9-12. Error bars denote mean ± S.E.M.

As individual differences in pre-injury climbing activity could be a pre-disposing factor for the levels of vertical climbing also after the induction of postoperative pain, baseline climbing activity was included as a covariate in the statistical analyses. For almost all outcome measures (except for top frequency: P = 0.055), it was found to significantly influence climbing throughout the experiments. This suggests that some of the individual propensity for high or low climbing activity is maintained after injury and should be factored into the analysis of the behaviour.

As the climbing deficits were most prominent at 4 h, we also assessed the effects of peri-operative treatment with the NSAID meloxicam (10 mg/kg s.c.), in the immediate post-operative period after incision injury. The dose was selected to be in the high end of the recommended dose-range (5-10 mg/kg) for mice [20]. At 4 h after injury, we only found a weak trend (P = 0.068) for analgesic improvement of climbing activity when focusing on the top frequency parameter, but no trend in distance moved or total climbing duration (**Fig 2-D1-3**). This suggests the analgesic compound was not efficient enough to noticeably restore climbing behaviour in this model.

### 3.2. Vertical climbing is decreased by inflammatory injury

We next examined vertical climbing in the CFA model of inflammatory pain [48] (**Fig 3**). Male and female C57BL/6NRj mice received a subcutaneous injection of CFA into the plantar surface of one hind paw and were followed for 11 days, measuring both vertical climbing and mechanical allodynia. The inflammatory injury produced consistent and robust evoked mechanical allodynia that persisted at least until day 7 after injection (**Fig 3A**). By comparison, the effect on vertical climbing was more subtle and primarily seen in the early phase of the injury. There was no significant decrease in the distance moved (**Fig 3B**, RM ANCOVA, sex*injury, covariate: baseline), but in contrast, both the frequency of climbing in the top (**Fig 3C**) and the total duration climbing (**Fig 3D**) were significantly reduced by the inflammatory injury across sex. The impairment was most pronounced in the early phase, particularly at Day 2 after injection (P=0.006-0.007, *post hoc* EMM with Bonferroni correction), was fully recovered on Day 11, in parallel with the resolution of mechanical allodynia.

**Fig 3.**
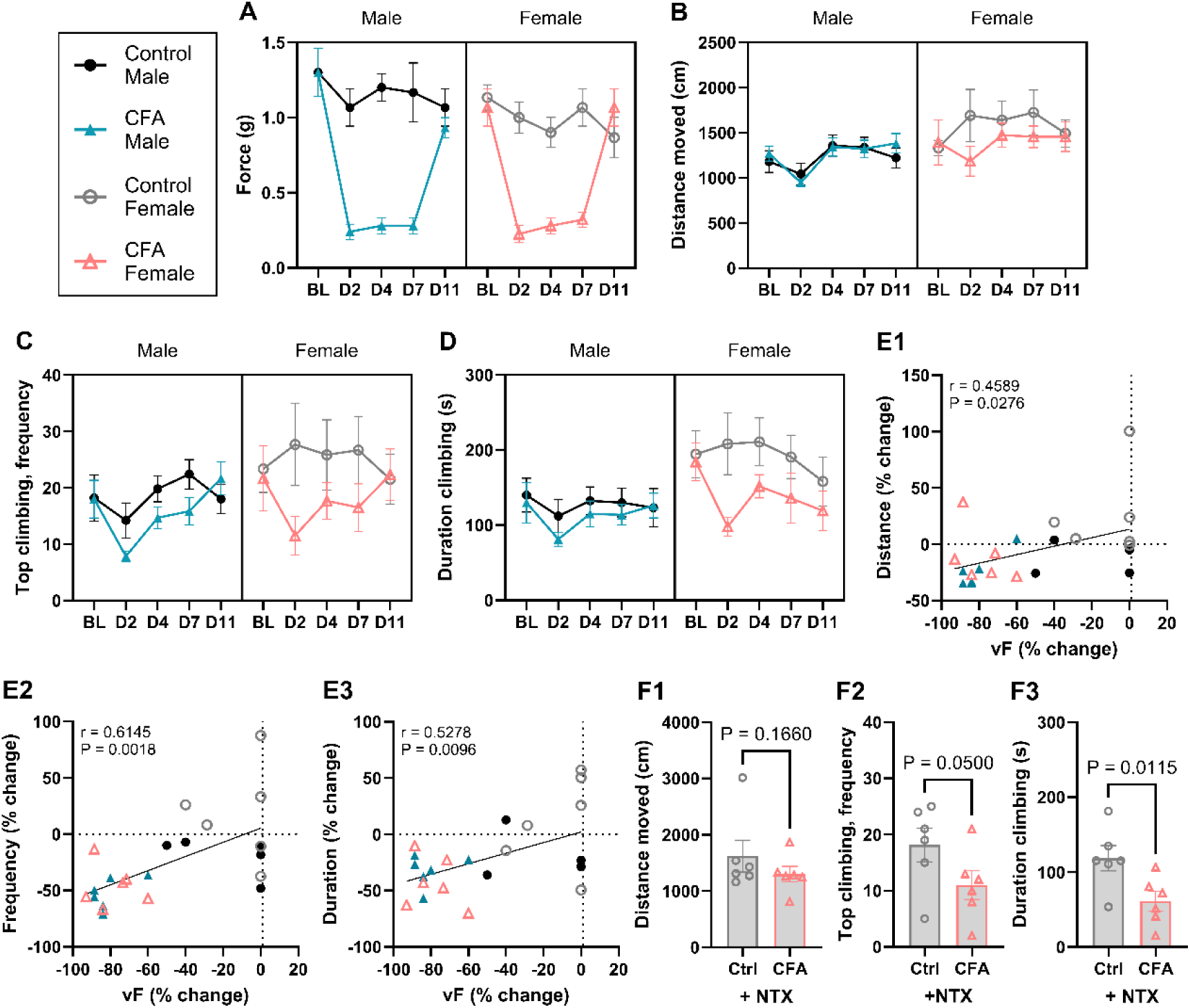
Inflammatory injury induces mechanical allodynia and decreases vertical climbing activity in the acute phase of the injury. Intraplantar injection of Complete Freund’s Adjuvant (CFA) was performed under isoflurane anaesthesia in male and female C57BL/6NRj mice. Mechanical allodynia and vertical climbing behaviour was assessed at baseline (BL) and 2, 4, 7, and 11 days after injury, and vertical climbing behaviour was quantified through automated video-tracking. (**A**) Time course development of CFA-induced mechanical allodynia, (**B)** Time course development of CFA-induced impairment in distance travelled (cm) during the 10 min test, (**C)** Time course development of CFA-induced impairment in top climbing frequency during the 10 min test, (**D)** Time course development of CFA-induced impairment in total duration climbing (seconds) during the 10 min test. (**E**) Correlations between vF-change and climbing-impairment on Day 2, for **(E1)** distance moved, **(E2)** top climbing frequency, and (E3) total duration climbing, as measured via Pearson’s correlation analysis. (**F)** Naltrexone (3 mg/kg) was administered to both control and CFA-injured female mice at Day 22, and 1 h later, they were assessed in the climbing cylinder for 10 min. This partially re-instated the climbing deficit for (**F1**) distance moved, (**F2**) top climbing frequency and (**F3**) total duration climbing (s). Error bars denote mean ± S.E.M. n = 5-6 per gender and treatment group.

We next investigated the potential correlation between the degree of mechanical allodynia and the decrease in climbing-related outcomes for individual animals. At the early peak of the inflammatory model (Day 2), we found significant correlations between the degree of mechanical allodynia and the decrease in distance travelled, number of times travelling to the top (top frequency) and the total duration climbing (**Fig 3-E1-3**). But from Day 4, the correlations were no longer significant (data not shown), suggesting a disconnection between the level of hypersensitivity and the climbing behaviour in the later stages of the disease.

To test whether the recovery of climbing depended on endogenous opioid signalling, we administered the opioid antagonist, naltrexone (NTX) on Day 22 to both control and CFA-injured female mice, a timepoint at which the injury no longer produced allodynia or climbing deficits NTX reliably re-instates the mechanical hypersensitivity in mice that have recovered form inflammatory injury [13], and we asked whether it would likewise reinstate the climbing deficit. NTX did so, and selectively for the measures that had been sensitive to the injury. Distance moved did not differ between CFA-injected animals and controls after NTX (**Fig 3-F1**), but NTX reinstated the deficit in both the frequency of reaching the top of the cylinder (**Fig 3-F2**), and the total duration climbing (**Fig 3-F3**). The recovery of climbing, like that of mechanical hypersensitivity, was therefore maintained by endogenous opioid signalling.

### 3.3. Vertical climbing is decreased by arthritis conditions

Having established that climbing is sensitive to injuries of the plantar hind paw, the very surface the animal uses to grip the wire mesh, we next wanted to investigate the effects of pain associated with the musculoskeletal system. We therefore first explored the changes in climbing behaviour in two commonly used models of joint pain, the local development of knee osteoarthritis in the MIA model [27; 49], and the systemic rheumatoid arthritis CAIA model [5].

MIA was induced in the left knee joint of male and female C57Bl/6JRj mice, and the behaviour was followed for 6 weeks thereafter. The MIA model is usually characterized by a biphasic impairment in weight bearing, with a strong initial deficit, peaking around Day 7 after injury, reflecting the acute reaction to the injected chemical, followed by a relatively steady state from two weeks onwards[27; 28]. Dynamic Weight Bearing (DWB) confirmed this profile with a marked drop at one week after injury (**Fig 4-A1**), followed by a less prominent and more steady state at two (**Fig 4-A2**) and four weeks (**Fig 4-A3**). At two weeks after injury, there was also a significant effect of sex on the data-set, as seen by the slightly higher deficit for the females, as often seen for this model[28]. Similarly, across sex, the climbing activity was significantly impaired, as seen by a reduced distance moved (**Fig 4B**), frequency of times making the way to the top (**Fig 4C**) and the total duration climbing (**Fig 4D**). Curiously, as this study was carried out over six weeks following injury, it was clear that the general climbing activity was decreasing with time, also for the control animals, which had received anaesthesia only. This may reflect ageing of the animals, as other types of climbing activity fall steeply between the age of 2 to 3 months of age [68], resembling the age trajectory seen here. As a result, the temporal dynamics of the climbing activity impairment did therefore not mirror the weight bearing profile completely, as both MIA and control animals were slowly decreasing in activity. Consistent with this, DWB and vertical climbing parameters only correlated at one week (**Fig 4-E1-3**), while there were no significant correlations at two and four weeks after injury (**Supplementary Fig S15**). Across the fulltime course, the analysis only showed significant effects of injury and time, but no effects of sex or sex*injury interactions. It should be noted, though, that the effects appeared somewhat blunted in females, particularly at later time points, where the steep decline in control activity may mask or dilute the effect of injury.

**Fig 4.**
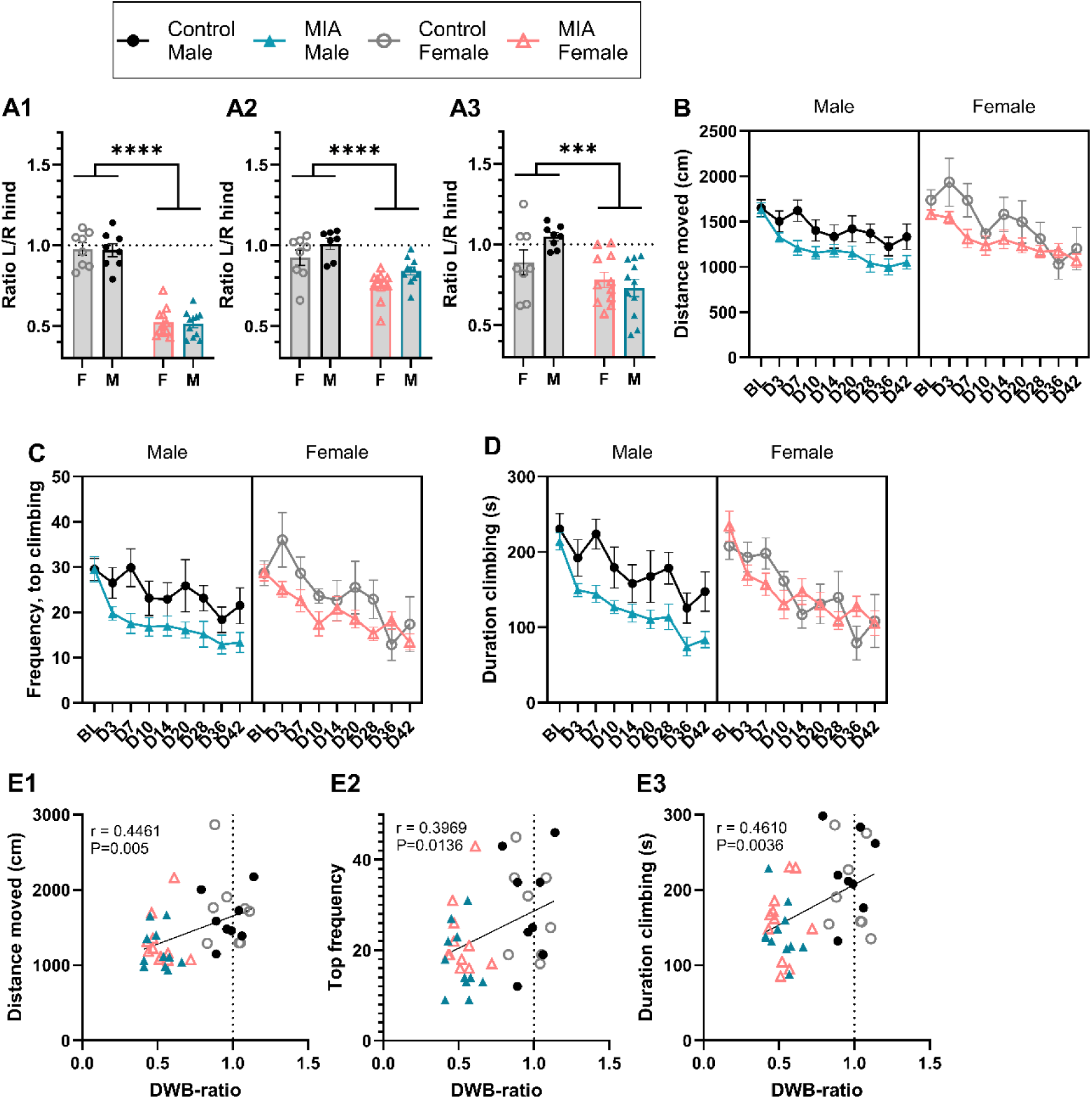
The osteoarthritis model, MIA, decreases dynamic weight bearing and vertical climbing activity. MIA was induced in the left hind leg of male and female C57BL/6JRj mice, and disease progression was confirmed using **A**) Dynamic Weight Bearing at (**A1**) Day 8/9, (**A2**) Day 15-16, and (**A3**) Day 29/30 after injury. Vertical climbing behaviour was recorded for 10 min at baseline (BL) and at D3, D7, D10, D14, D20, D28, D36 and D42 after model induction, and displayed as (**B**) distance moved (cm), (**C**) the frequency in the top of the climbing cylinder, and (**D**) total duration climbing (s). (**E**) At Day 7-9, there were significant correlation between the Dynamic Weight Bearing deficit and the decreased vertical climbing outcome measures, (**E1**) distance moved (cm) vs DWB-ratio, (**E2**) top frequency vs DWB-ratio, and (**E3**) duration climbing (s) vs. DWB-ratio, as measured via Pearson’s correlation analysis. Error bars denote mean ± S.E.M. n: controls: 8, MIA: 11.

The rheumatoid arthritis model, CAIA [5], was induced in BALB/cByJRj mice by an i.v. injection of collagen antibodies (AB) on Day 0, followed by an i.p. injection of LPS on Day 3, to trigger the inflammatory response. Two different doses of LPS were used (5 µg and 10 µg) to assess whether the LPS dose affected model development. Arthritis scores followed the expected disease development, with visible signs of arthritis appearing between Day 4 to 5 after AB administration, meaning in the first days after LPS injection (**Fig 5A**).

**Fig 5.**
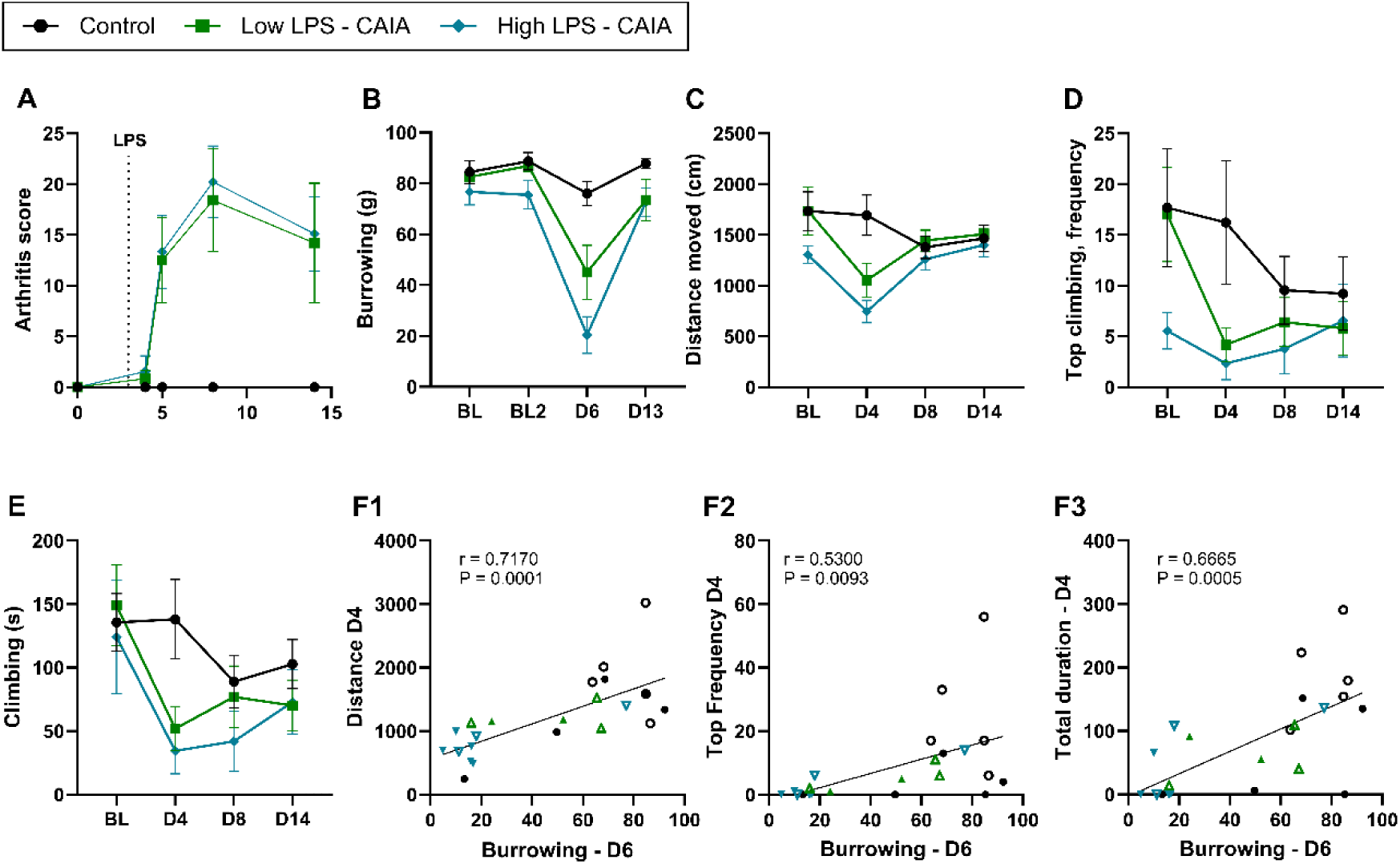
The rheumatoid arthritis model, CAIA, decreased both burrowing and vertical climbing activity. CAIA was induced in male and female BALB/cByJRj mice through an i.v. injection of antibodies on Day 0, and on Day 3, a dose of LPS was given, either at a low (5 µg) or high (10 µg) dose. (**A**) Arthritis scores quantified the number and extent of inflammatory active joint sites. (**B**) Burrowing behaviour was assessed through the removal of bedding material from a burrowing tube during a 2 h session at baseline and on Days 6 and 13. The maximum amount of bedding that could be removed from the tube was 100 g. **C-E**: Vertical climbing was recorded for 10 min at baseline and on Days 4, 8 and 14, and displayed as **(C**) Distance moved (cm), (**D**) frequency of climbing to the top of the climbing cylinder, and (**E**) total duration climbing (s). (**F)** Correlations were tested between burrowing (**D6**) and the different climbing outcomes that were measured on Day 4, as seen for **F1**) distance moved (cm), **F2**) frequency in the top, and **F3**) duration climbing (s), as measured via Pearson’s correlation analysis. As 50 % of the animals were excluded from the climbing study due to pre-defined exclusion criteria, timeline data is presented combined for males and females, although sex is included as a factor in the analysis, and individual datapoints are marked according to sex: Males: closed symbols. Females: open symbols. Arthritis- and burrowing scores are only presented from the animals that were included in the climbing analysis. Error bars denote mean ± S.E.M. Control: n = 9 (f:5/m:4), low: n = 5 (f:3/m:2), high: n = 9 (f:4/m:5).

Burrowing behaviour, assessed on Days 6 and 13 as a reference pain-associated behaviour, was significantly impaired by injury, but a significant injury*time interaction suggests that the effect of injury was not consistent throughout the study (**Fig 5B**). *Post hoc* comparison of Estimated Marginal Means (EMM) confirmed that on Day 6 burrowing was significantly impaired in the high-LPS group (P < 0.001), while only trending for the low-LPS (P = 0.052). The two LPS did not differ from each other, and no differences remained on Day 13.

Vertical climbing followed a similar pattern. Across the study period, distance moved showed no main effects of injury, but a significant interaction between time and injury, suggesting that the injury effects varied over time (**Fig 5C**). *Post hoc* comparison of EMM detected a significant decrease in distance moved for both LPS groups on Day 4, but not at the later timepoints (**Supplementary Table S3**). Frequency of reaching the top was highly variable between individuals and showed no significant effects or interactions (**Fig 5D**). Similarly, total climbing duration showed only trends for injury (P = 0.079) and for time*injury interaction (P=0.057) however, *post hoc* comparison again revealed a significant reduction in the high-LPS group on Day 4, but not for the low-LPS (P=0.098) (**Fig 5E**).

Burrowing and climbing deficits were also related at the individual level. The degree of impairment on burrowing (Day 6) and vertical climbing behaviour correlated significantly on Day 4, measured by distance moved (**Fig 5-F1**), top frequency (**Fig 5-F2**) and total duration climbing (**Fig 5-F3**), and with total climbing at D13/14 (**Supplementary Fig S16A**), indicating that these two parameters were equally impaired by the disease. Similarly, climbing also correlated significantly with arthritis scores (**Supplementary Fig S16B**), linking the level of climbing to the disease burden.

These results must be interpreted with caution. The poor baseline climbing performance of the BALB/cByJRj strain and the resulting exclusions produced unequal group sizes and unequal baseline levels, particularly for distance moved and top climbing frequency, and with 50 % of the planned animals excluded the study is underpowered. Equal numbers of males and females were excluded, and as statistical analysis showed no significant effects of sex for any of the outcome measures, we have pooled males and females for the graphical presentation (**Fig 5**), although sex was still included as an explanatory variable in the statistical analysis (**Supplementary Table 5**). With these caveats, these results suggest that the vertical climbing behaviour is most prominently impaired in the acute phase of the CAIA model development, which aligns well with the results from the burrowing test.

### 3.4. Vertical climbing is decreased by bone pain conditions

After examining the impact of conditions affecting the joints, we next turned to skeletal and bone pain. Therefore, we first explored two different cancer-induced bone pain (CIBP) models, where cancer cells (breast cancer or multiple myeloma) were injected into the femoral bone, to model cancer metastasis to the bone.

Breast CIBP [30] was induced in female BALB/cOlaHSD mice, which produced the expected decrease in limb use (**Fig 6A**) and weight bearing (**Fig 6B**), with a gradual onset from Day 8 onwards. Climbing was affected in parallel with tumour growth: distance moved (**Fig 6C**), frequency of reaching the top of the cylinder (**Fig 6D**) and duration of climbing activity (**Fig 6E**) all showed a significant injury*time interaction across the full time course dataset, whereas the main effects effect of injury was diluted by the disease-developmental phase before Day 8. *Post hoc* EMM comparisons resolved the interaction and identified significant effects of injury on Days 12-14 for all three measures. Additionally, comparing with the reference behaviour, static weight bearing, there were significant correlations between the level of weight bearing and climbing impairment at the last timepoint measured for each animal (**Fig 6F1-3**).

**Fig 6.**
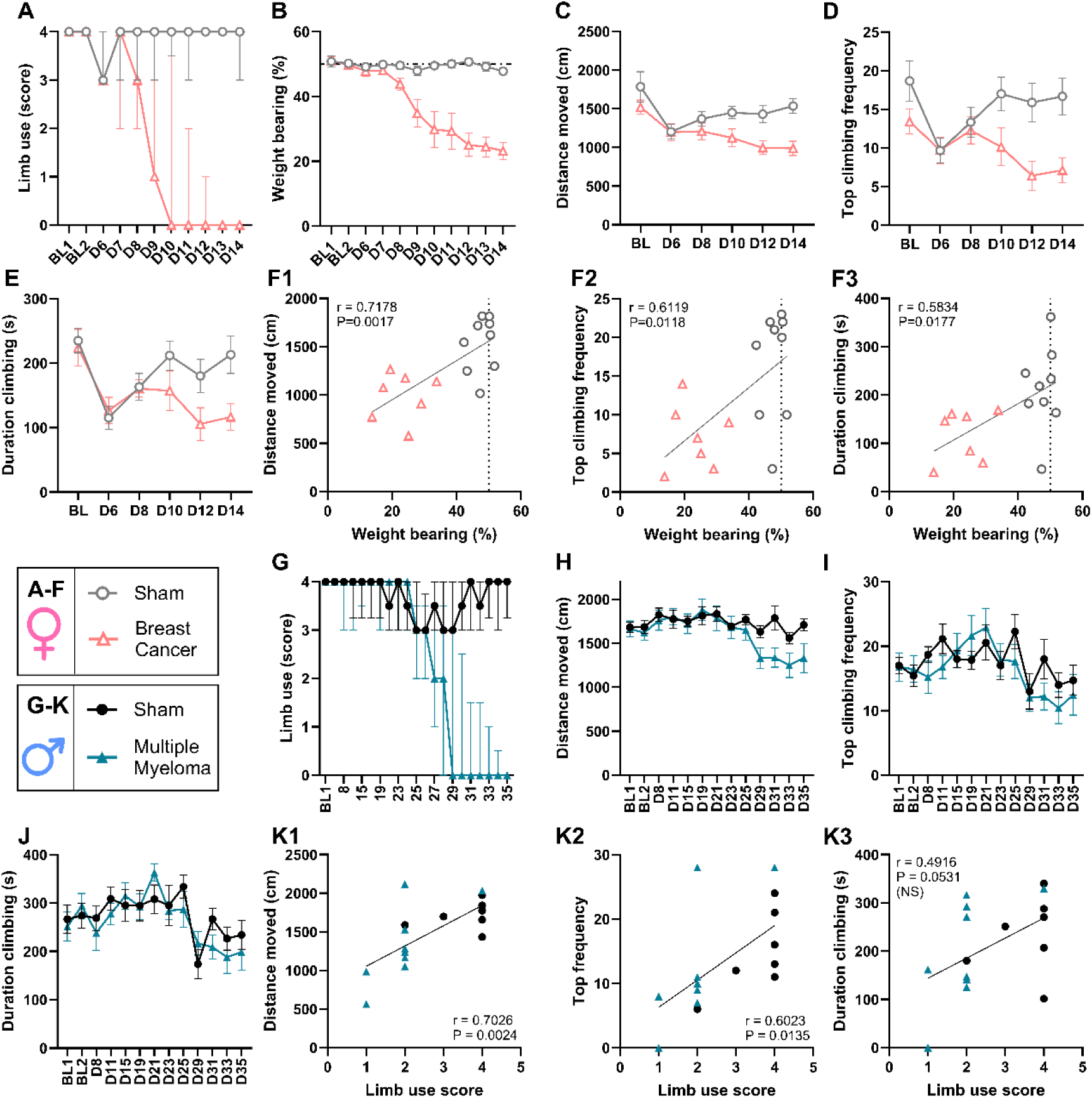
Cancer-induced bone pain models decrease vertical climbing activity. CIBP was induced via intra-femoral injection of cancerous cells during anaesthesia, and vertical climbing, weight bearing and limb use were recorded. **A-F**: Breast Cancer Induced Bone Pain was induced in female BALB/cOlaHSD mice (sham: n = 9, cancer: n = 7) and they were monitored for 14 days, or until reaching the humane endpoint. (**A**) Limb use was qualitatively scored daily. (**B**) Time course development of impairment in static weightbearing (%). Vertical climbing behaviour was recorded for 10 min at baseline (BL) and at Days 6, 8, 10, 12 and 14 after model induction, and displayed as **(C**) distance moved (cm), (**D**) frequency of climbing to the top of the climbing cylinder, and (**E**) total duration climbing (s). (**F**) At day the last measure for each animal, there were significant correlation between the weight bearing deficit (%) and the decreased vertical climbing outcome measures, (**F1**) distance moved (cm), (**F2**) top frequency, and (**F3**) duration climbing (s), as measured via Pearson’s correlation analysis. **G-K**: Multiple Myeloma Cancer Induced Bone Pain was induced in male C57BL/KaLwRijHsd mice (sham: n = 7, cancer: n = 9) and they were followed for 35 days, or until reaching the humane endpoint. (**G**) Limb use was qualitatively scored daily. Vertical climbing behaviour was recorded for 10 min at baseline (BL) and at selected days between Day 8 to 35 after model induction and is displayed as (**H**) distance moved (cm), (**I**) frequency in the top of the climbing cylinder, and (**J**) total duration climbing (s). (**K**) At day the last measure for each animal, there were significant correlation between the weight bearing deficit (%) and the decreased vertical climbing outcome measures, (**K1**) distance moved (cm), (**K2**) top frequency, and (**K3**) duration climbing (s), as measured via Pearson’s correlation analysis. All data is “LOCF” = last observation carried forward. Limb use scores: error bars denote median ± interquartile range. Other datasets: error bars denote mean ± S.E.M.

Secondly, multiple myeloma CIBP [17] was induced in male C57BL/KaLwRijHsd mice. As seen by the limb use scores (**Fig 6G**), this is a model of slower disease progression, with a decrease in limb use only from approximately 27 days after induction of the model. The late and variable development of the disease was also seen in the vertical climbing behaviour, where signs of impairment only started to emerge in the late stages. The model induced a significant time*injury interaction for the distance moved in the climbing cylinder, suggesting that the injury effects were different over time. We therefore compared EMM and found that the injury only produced a significant decrease in distance moved on Day 31, while trending on D29, D33 and D35 (P=0.07-0.10) (**Fig 6H**). For the other two main parameters, top climbing frequency (**Fig 6I**) and climbing duration (**Fig 6J**), the effects were more blunted and did not reach significance. At the end of the study, limb use correlated significantly distance moved (**Fig 6-K1**) and top frequency (**Fig 6-K2**), but showed only a trend for correlation with the total duration climbing (P=0.0531, **Fig 6-K3**).

As both breast cancer and multiple myeloma CIBP models were unilateral cancer-induced models, we wanted to assess the impact of a non-cancerous disseminated bone disease and therefore assessed a doxycycline-induced transgenic mouse model of fibrous dysplasia (FD). Mice were single housed for the experiment for assessment of general activity in the home cage using automated home cage monitoring system (Digital Ventilated Cage). Approximately a week after doxycycline administration was initiated, FD mice began to show reduced home cage activity (**Fig 7A**), with significant effects of injury and injury*time interaction. EMM comparisons confirmed significant *post hoc* differences between control and FD mice from Day 8 to 15 after initiation of doxycycline, after which the locomotion approached normal levels. Voluntary wheel running activity in the home cage appeared more severely affected, with significant effects of injury, with an injury*time interaction, and with significant *post hoc* comparisons already from Day 7 and almost to study end (**Fig 7B**). Bone damage was assessed at selected timepoints and showed a progressive decline in scores from Day 8 onwards (**Fig 7C**).

**Fig 7.**
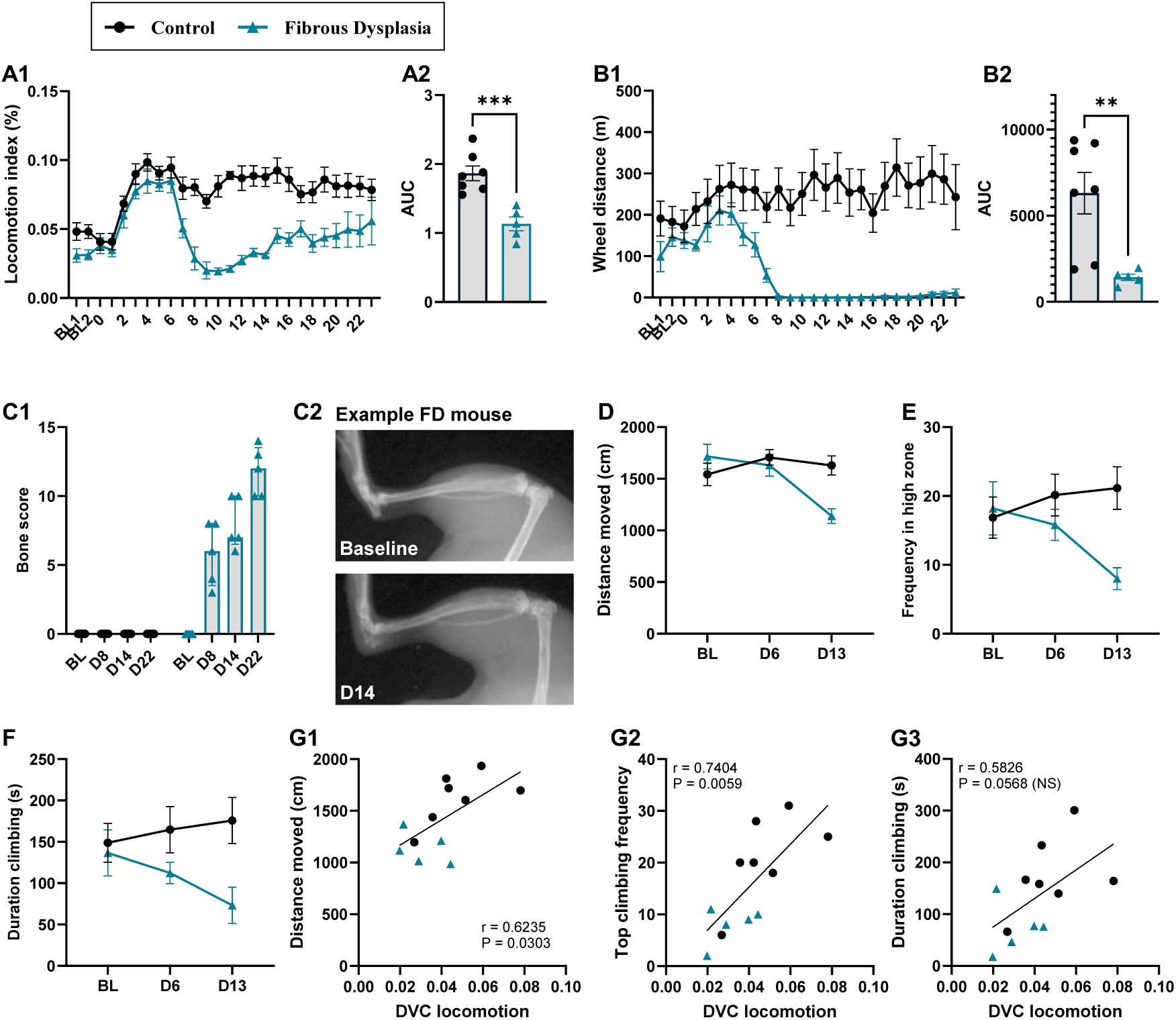
Fibrous Dysplasia decreases general home cage activity and vertical climbing behaviour. Fibrous dysplasia, a non-cancerous bone pain model, was induced in a transgenic mouse line, induced via doxycycline given to both control and FD mice in the drinking water throughout the experiment. (**A1**) Locomotor activity was measured continuously and non-invasively in the home cage via sensors below the cage (DVC system). Activity is only displayed from the animal’s active phase (nighttime). **A2**: AUC from the entire study-period. (**B1**) Wheel running distances (m) were measured continuously and non-invasively in the home cage, and are displayed for the animals’ active phase (nighttime). **B2**: AUC from the entire study period. (**C1**) X-ray scoring of bone damage at different timepoints during the experiment. **C2**: X-ray example from mouse 245 at Day 14. **D-F**: Vertical climbing was recorded for 10 min at baseline and on Days 6 and 13 and displayed as **(D**) distance moved (cm), (**E**) frequency of climbing to the top of the climbing cylinder, and (**F**) total climbing duration (s). **G1-3:** Correlations were tested between home cage locomotion on D13 and the different climbing outcome measures on the same day, as seen for (**G1**) distance moved, (**G2**) Frequency in the top, and (**G3**) duration climbing, as measured via Pearson’s correlation analysis. Males only, all animals were single housed. FD: n = 5, control: n = 7. BL = baseline. Bone scores: error bars denote median ± interquartile range. Other datasets: error bars denote mean ± S.E.M.

Vertical climbing behaviour was only evaluated at Days 6 and 13 and showed significant impairment by injury for all three outcomes measures, distance moved (**Fig 7D**), frequency in the top (**Fig 7E**) and total climbing duration (**Fig 7F**). For all parameters, there were also significant injury*time interactions, and EMM comparisons at the different timepoints clarified that the climbing activity was only significantly altered at Day 13. Home cage locomotor activity correlated significantly with climbing behaviour on Day 13 for all three measures, distance moved (**Fig 7-G1**), top climbing frequency (**Fig 7-G2**) and total duration climbing (**Fig 7-G3**). Similarly, significant correlations were also observed between the climbing parameters and the wheel running activity (**Supplementary Fig S17A**) and the X-ray scores on day 14 (**Supplementary Fig S17B**).

Taken together, these results show that the vertical climbing behaviour is markedly impaired by the fibrous dysplasia model in a time-dependent manner, and with significant correlation between degree of impairment and both home cage activity and disease severity.

### 3.5. Vertical climbing is decreased by neuropathic pain and partially restored by analgesics treatment

We next turned to neuropathic pain and the spared nerve injury (SNI) model [33], ore of the most widely used preclinical models of chronic pain. Here, we induced the SNI injury in male and female C57BL/6NRj mice and characterised their vertical climbing activity for 16 days after injury. Vertical climbing was significantly impaired by injury, as seen by a reduction in the distance moved (**Fig 8A**), the number of times reaching the top of the cylinder (**Fig 8B**), and the total duration climbing on the wire mesh (**Fig 8C**). Of note, for this injury model, sex was found to be a significant factor for both the distance moved and the top frequency, with females generally climbing less in the later stages, but statistically, there was no sex*injury interaction, likely due to the SNI-injury producing reduced climbing activity for both sexes. After confirming that the model produced impaired climbing activity, we wanted to assess whether the behaviour could be reinstated after administration of an analgesic. This was particularly important as the SNI injury also produces damage to motor neurons, and an analgesic treatment would help clarify if the decrease in vertical climbing was related to the sensory or motor effects of the model. The most used reference compounds for neuropathic pain are the gabapentinoids: pregabalin and gabapentin, although these are known to produce potential sedative effects [23; 58; 62]. Therefore, we carefully selected a relatively low dose of gabapentin (GBP, 30 mg/kg) and also administered it to the sham controls, to detect potential side effects affecting the general locomotor activity. On Days 27 and 30 after injury, we performed a cross-over study comparing vehicle vs. GBP in both SNI and sham controls and found no effects of GBP in sham animals (**Supplementary Fig S18**), confirming that the dose was neither sedative nor inducing hyperactivity. Contrarily, when GBP was given to SNI animals, there was a significant increase in vertical climbing activity, as measured by distance moved (**Fig 8-D1**), and climbing duration (**Fig 8-D2**), while only a trend for GBP-induced increased frequency in the top was observed (P = 0.059, **Fig 8-D3**). This suggests that the analgesic effects of GBP partially reinstated the vertical climbing behaviour and thereby validated the usefulness of the test for pain-related outcomes.

**Fig 8.**
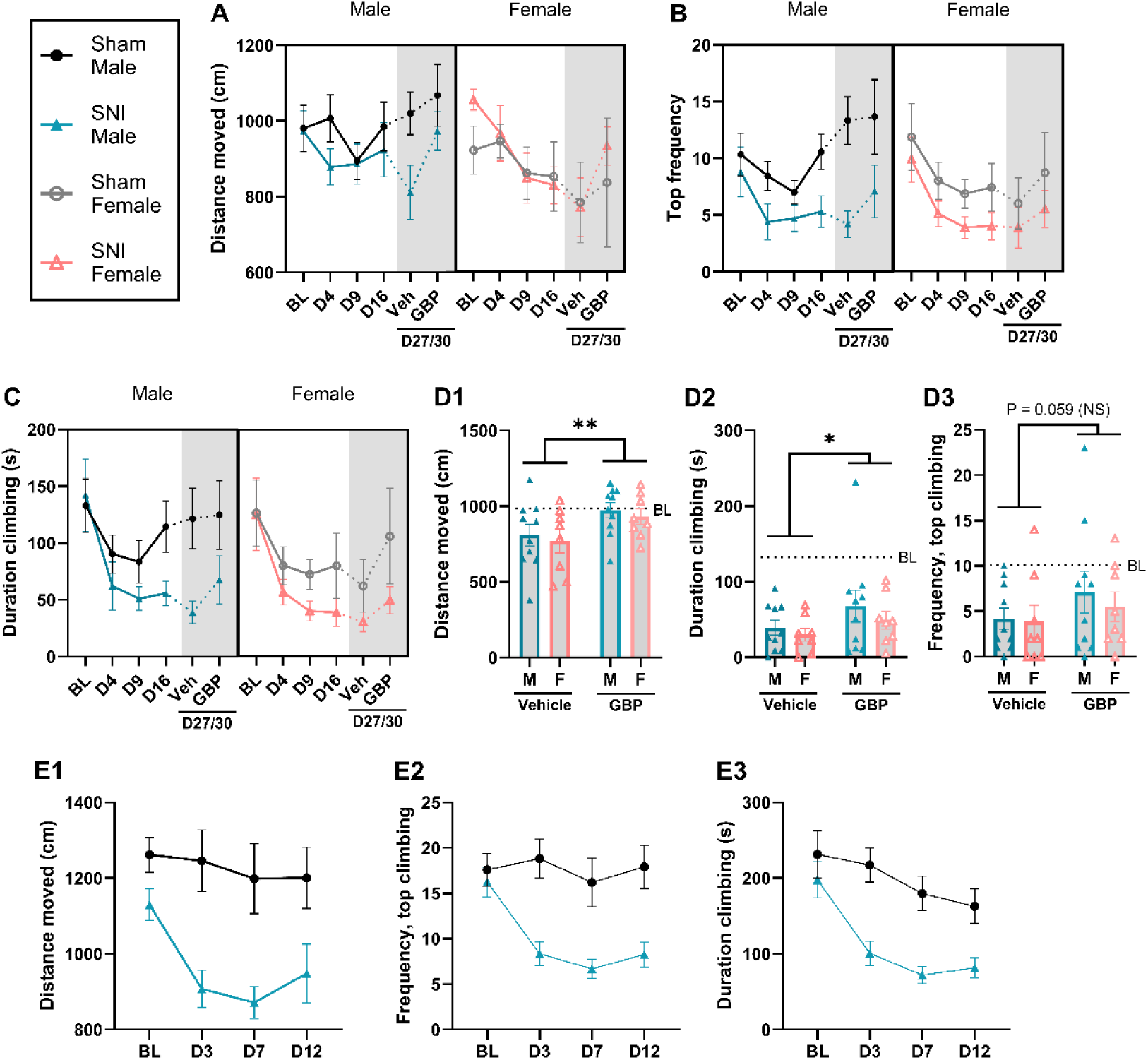
Neuropathic pain decreases vertical climbing and is partially restored by gabapentin. Spared Nerve Injury (SNI) was induced under isoflurane anaesthesia in male and female C57BL/6NRj mice in comparison with sham-injured controls (n = 7-10). **A-D**: Vertical climbing was recorded for 10 min at baseline and on Day 4, 9, 16 and during a gabapentin-study on Day 27/30 displayed as **(A**) distance moved (cm), (**B**) frequency of climbing to the top of the climbing cylinder, and (**C**) total duration climbing (s). (**D**) On Day 27/30 a cross-over study testing i.p. gabapentin (GBP, 30 mg/kg) vs. vehicle was conducted, showing that GBP partially restored climbing behaviour, as measures through (**D1**) distance moved, (**D2**) duration climbing, and (**D3**) frequency in the top. Dotted line suggests pre-injury baseline average. *: P<0.05, **: P<0.01, NS: not significant, as suggested by injury-effect in a 2-way ANOVA (factors: sex & injury). F = female, M = male. (**E**) During another experiment, SNI-injured males were compared with naïve controls in vertical climbing, as measured via (**E1**) distance moved (cm), (**E2**) frequency in the top, and (**E3**) duration climbing (s) (n = 10-12). Error bars denote mean ± S.E.M.

Of note, across the time course of the SNI study, there was also a clear decrease in activity for the sham injured control animals, particularly during the first test days, up until Day 9 (**Fig 8A-C**). This suggests that the surgery and associated wound healing, in itself, produced some decrease in activity. Therefore, another cohort of animals was tested, where the control animals had not undergone surgery. Interestingly, under those conditions, the decline in activity for the controls was less prominent, producing a larger effect size between SNI and control (**Fig 8-E1-3**). Additionally, the prominent decrease in vertical climbing activity from the SNI-injury was confirmed.

### 3.6. Vertical climbing is also decreased by models not causing pain directly to the locomotor-system

Next, we wanted to assess the potential effects pain in models not directly affecting the limbs, like models causing pain-related symptoms in the head and face. We therefore first looked to the model of levcromakalim (LEV)-induced migraine[9], which was induced in male and female C57BL/6JBomTac. The Mouse Grimace Scale, a measure of non-evoked pain-like behaviour in migraine-models [39], showed a significant increase in grimacing from baseline to Day 2, confirming the presence of pain-like behaviours and successful model induction (**Fig 9A**). Vertical climbing behaviour was assessed at both 1 and 2 h after injection on Days 0 and 4. For the frequency climbing to the top, there were no significant effects of LEV, sex, or any interactions, although the effects clearly depended on whether the assessment was conducted at 1 h or 2 h after the injection (**Fig 9B**). EMM comparisons across sex only found significant differences between LEV and vehicle groups on the first day at 1 h post-induction (P=0.028) and a trend at D4-1h (P=0.072), while no differences at the 2h timepoints. Total duration climbing appeared to be a more sensitive outcome for this model, as significant effects were detected for LEV, but also with significant interactions between LEV and time, suggesting that the effects changed over time (**Fig 9C**). *Post hoc* EMM comparisons showed significant differences between control and LEV at the 1-hour, but not the 2–hour, measures on both days. The increase in grimacing on correlated significantly with both frequency (**Fig 9-D1**) and total duration (**Fig 9-D2**) at Day 0-1h.

**Fig 9.**
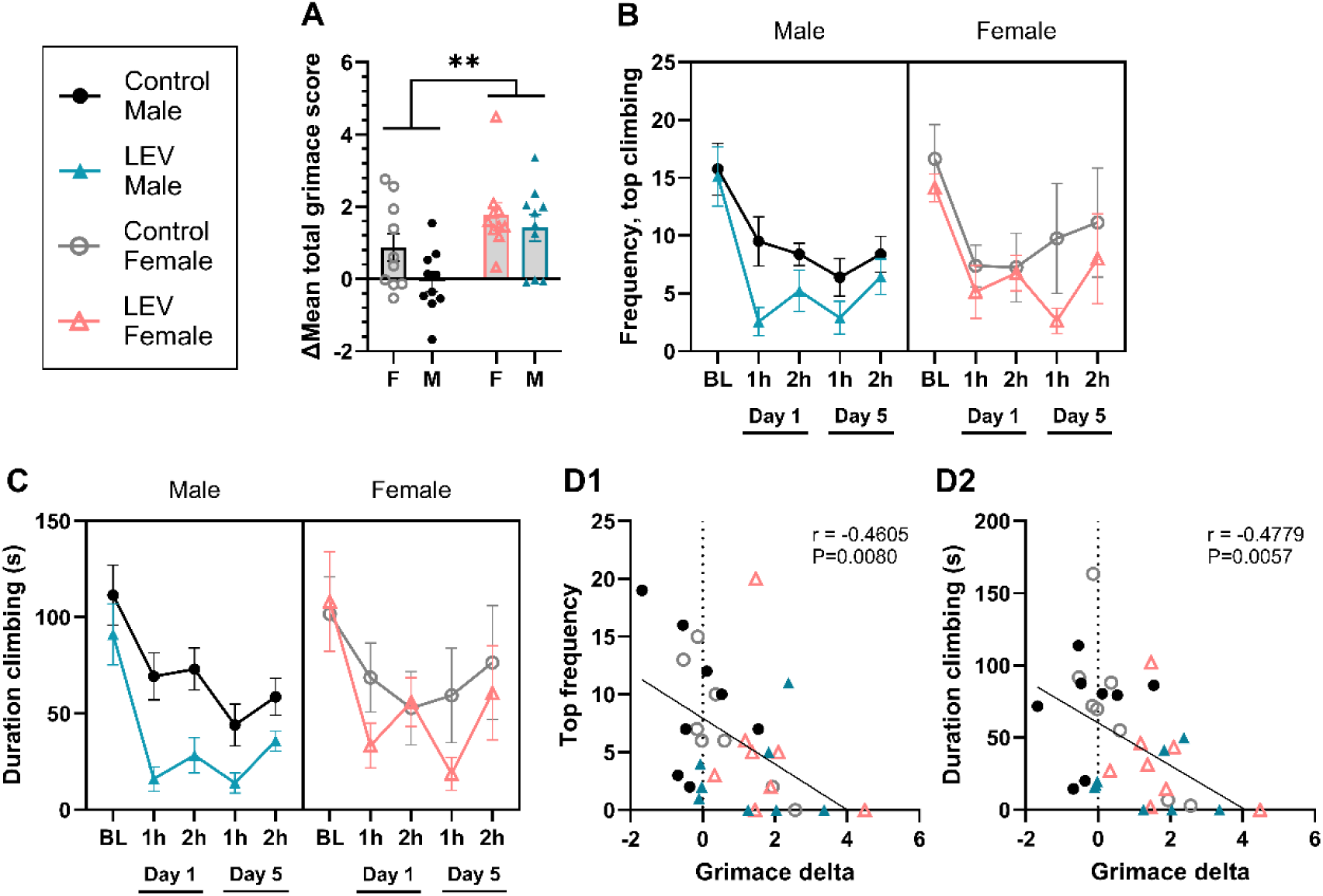
Vertical climbing was impaired in the model of levcromakalim-induced migraine with direct correlation to grimacing behaviour. Migraine-relevant pain was induced in mice by i.p. injection of levcromakalim (LEV) on Days 0, 2 and 4. (**A**) Mouse Grimace Scale was assessed at baseline and on Day 2, 1 hour after injection, with displayed values showing an increase from baseline to D2 in mean total grimacing. **: P = 0.0014, suggest significant effect of LEV across sex, as determined using 2-way ANOVA (sex*LEV). F = female, M = male. **B-C**: Vertical climbing was recorded for 10 min at baseline and 1 and 2 h after injection on Days 0 & 4, and displayed as (**B**) frequency of climbing to the top of the climbing cylinder, and (**C**) total duration climbing (s). **D1-2:** Correlations were assessed between MGS (**D2**) and the different climbing outcomes that were measured on Day 0, as seen for Fig. **D1**) Frequency in the top, and **D2**) Duration climbing (s), as measured via Pearson’s correlation analysis. Males, veh: n = 8, LEV: n = 9, females: veh: n = 8, LEV: n = 8. Error bars denotes mean ± S.E.M.

This suggests that the vertical climbing is more severely affected in the LEV model at the early timepoint (1 h), and that it significantly correlates with the degree of facial grimacing from the model.

Distance moved is not presented for this dataset as the contrast between the climbing stand and the mouse’s coat colour proved difficult to distinguish for the infrared camera, used for this experiment. The animal was not accurately detected when on the bottom, in the non-climbing zone. This made the distance parameter imprecise, as the animal could not be accurately tracked when moving in the bottom. This did, however, not compromise the parameters where the animal was climbing. Upon adaptation with a white stand, this problem was solved.

We then examined the FRICT-ION model of periorbital neuropathic pain [44], which neither affects a limb used for climbing nor is it expected to be aggravated by physical activity, and in which pain should be confined to the facial region. The model produced robust and significant mechanical (**Fig. 10A**) and cold allodynia (**Fig. 10B**) in the orofacial region. Climbing was less markedly affected: injury had no significant effect on distance moved in the cylinder (**Fig. 10C**) but significantly reduced both the frequency of climbing to the top (**Fig. 10D**) and total climbing duration (**Fig. 10E**). There were no significant effects of time or sex, whereas the baseline covariate was significant for all three measures. Mechanical allodynia did not correlate with the distance moved or top frequency (**Fig. 10-F1** and **F2**), but it did correlate significantly with climbing duration (**Fig. 10-F3**). Cold allodynia, in turn, did not correlate with any of the climbing parameters (**Supplementary Fig. S19**). Thus, although FRICT-ION produces only orofacial signs of pain, it still reduces the willingness to climb, indicating that the climbing deficit does not depend on pain associated with limb use or exercise. At the same time, the level of climbing was only weakly related to mechanical allodynia.

**Fig 10.**
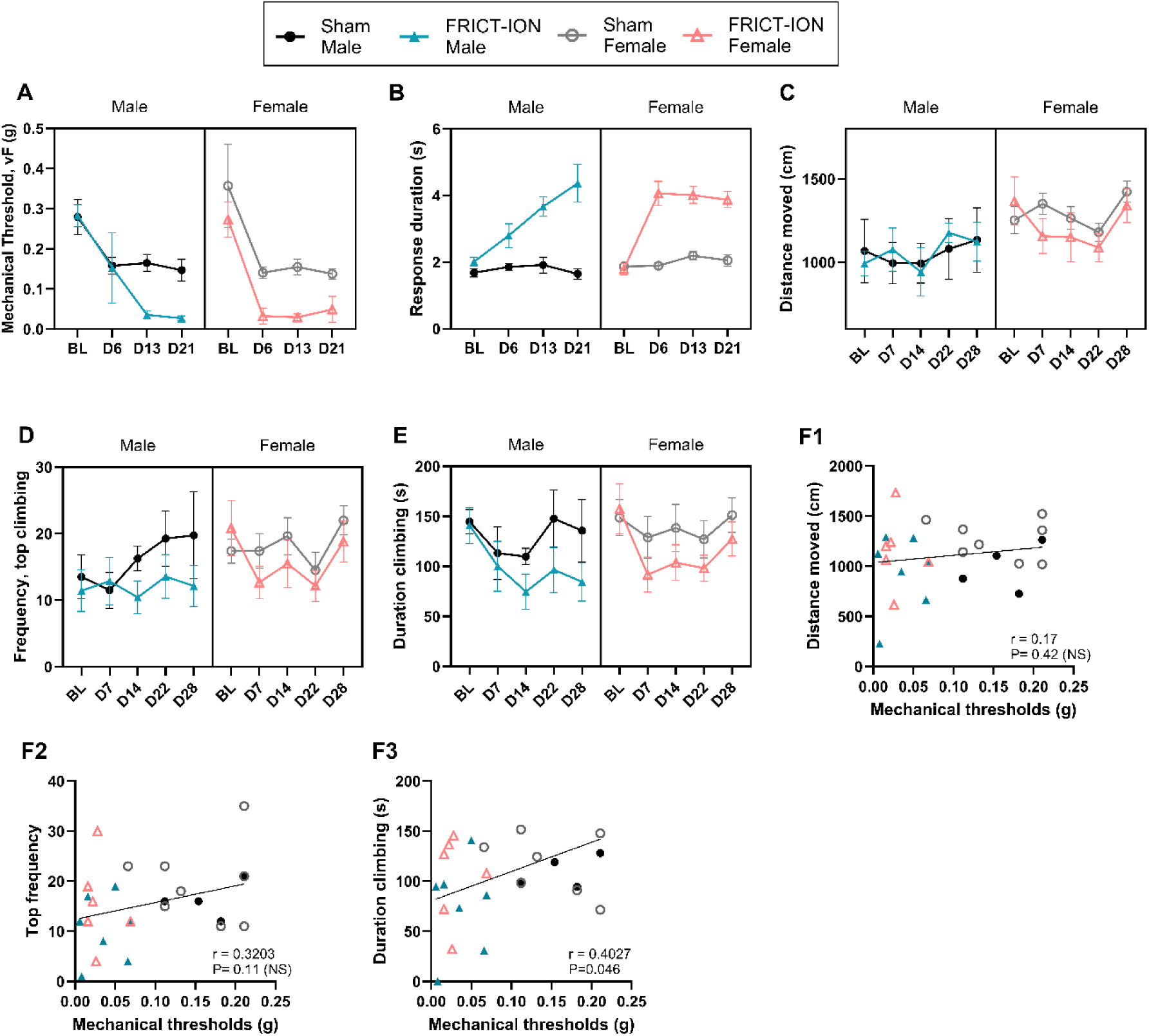
The periorbital neuropathic pain model, FRICT-ION, induces mechanical and cold allodynia, and decreases vertical climbing activity. The FRICT-ION model was induced under isoflurane anaesthesia in male and female C57BL/6J mice. Mechanical and cold allodynia, and vertical climbing behaviour was assessed at baseline (BL) and at 1, 2, 3, and 4 weeks after injury. **A:** Mechanical allodynia was suggested by a decline in von Frey threshold (g). **B:** Cold allodynia was suggested by an increase in responding to a drop of acetone, applied to the animal’s face. **C-E**: Vertical climbing was recorded for 10 min and the results are displayed as **(C**) distance moved (cm), (**D**) frequency of climbing to the top of the climbing cylinder, and (**E**) total duration climbing (s). (**F**) Correlations between the mechanical response threshold and climbing-impairment at Day 14, for (**F1**) distance moved, (**F2**) top climbing frequency, and (**F3**) total duration climbing, as measured via Pearson’s correlation analysis. Error bars denote mean ± S.E.M. n = 4-8 per sex-specific group.

### 3.7. Baseline vertical climbing levels differ between experiments and mouse strains and are differentially affected by the type of pain-related model

Finally, we wanted to compare the climbing performance across studies. As a pre-requisite for the assay, it was important to ascertain that animals were motivated to climb and comfortable manoeuvring around the vertical assay before starting the experiment to assess potential model-induced impairments. Exclusion criteria were therefore applied, and although the majority of the experiments saw relatively low levels of exclusions (0-13 %), two sub-strains showed a markedly higher level of exclusion in all cohorts in their studies, namely the C57BL6/J-BOMTAC used for the LEV-experiments (28 % excluded), and the BALB/cByJRj used for the CAIA studies (50 % excluded) (**Fig 11A**), suggesting these two sub-strains are poor climbers. Of note, there were no significant differences in exclusions for the two sexes.

**Fig 11:**
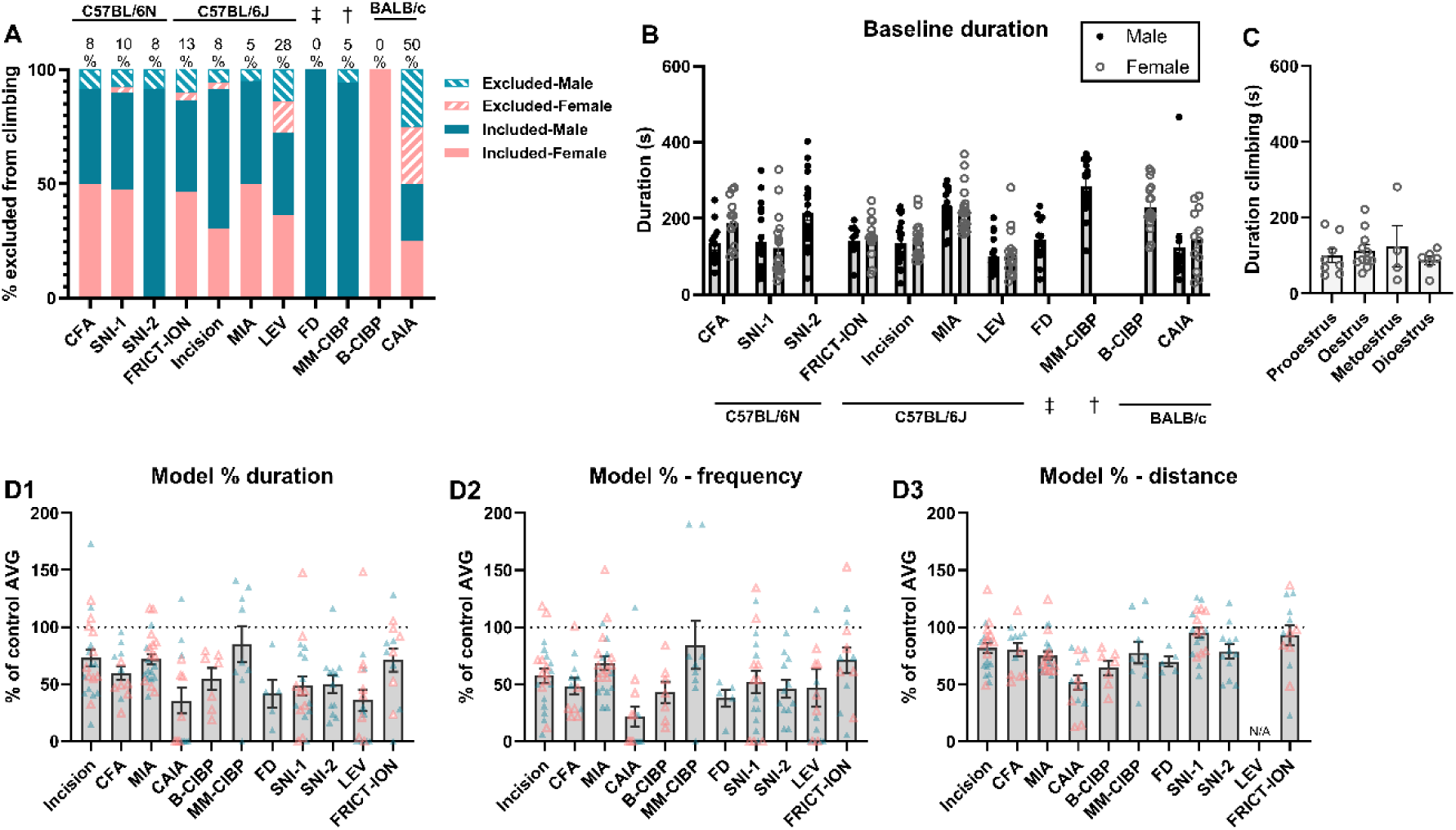
Comparisons across experiments at baseline and at most affected timepoint for each model. A: Exclusions. **B**. Comparison of raw baselines from all experiments. **C**: Oestrus cycle at baseline. **D**. % decrease in vertical climbing at the most severely affected timepoint for each model, as normalised to control average at the same timepoint (Incision: 4h. CFA: D2. MIA: D7. CAIA (low + high groups): D4. SNI: D12/16. Bone pain-models: the last measure. LEV: D0-1h, FRICT-ION: D14), as measured by **C1**) total duration climbing, **C2**) top frequency, and **C3**) distance moved. Models: Incision: plantar incisions model, CFA: intraplantar Complete Freund’s Adjuvant, MIA: Monoiodoacetate Induced Arthritis, CAIA: Collagen Antibody Induced Arthritis, Breast CIBP: Breast Cancer Induced Bone Pain, MM CIBP: Multiple Myeloma Cancer Induced Bone Pain, FD: Fibrous Dysplasia, SNI: Spared Nerve Injury (SNI-1: male-female experiment, SNI-2: male-control experiment), LEV: Levcromacalim-induced migraine, FRICT-ION: Foramen Rotundum Inflammatory Constriction Trigeminal Infraorbital Nerve injury. Strains: † = C57BL/KaLwRijHsd, ‡ = FD mice carrying (Prrx1-Cre⁺; Linker⁺; Gαs^R201C/+^).

Combining baselines of the included animals from all studies showed no significant effects of sex on the climbing performance (**Fig 11B**), clearly suggesting that males and females have comparable innate vertical climbing activity. On the other hand, there were significant differences in level of activity for the different experiments, before the model induction. And although some differences could be strain-specific, *post hoc* comparison showed significant differences even when the experiment was conducted using the same vendor-supplied mouse sub-strain, age, protocol and in the same facility and room. For example, there were significant differences in baseline climbing duration between the two SNI experiments (Tukey’s multiple comparisons test: P = 0.0003). As these experiments were done by different experimenters eight months apart, it is uncertain how much of the baseline differences were caused by experimenter- or animal-related differences. Similarly, using the same strain, albeit different experimenters and behavioural rooms, there were also significant baseline differences between baseline climbing activity for the MIA and incision experiments, performed in the same - vendor-supplied sub-strain (P<0.0001). Overall, as baseline levels fluctuate between experiments, appropriate controls are key for this outcomes measure.

On a subset of female mice (35 female C57BL/6JBomTac mice), the impact of oestrus cycle on baseline climbing behaviour was assessed. Naturally, as the oestrus cycle is a fluctuating biological process, there was not an even spread of animals in each cycle stage, but the cycle-stage did not affect the animals’ baseline climbing levels (**Fig 11C**). Five of these females were excluded due to the pre-defined exclusion criteria, but the excluded animals were also evenly distributed across the different cycle stages, meaning that poor climbing compliance could not be explained by the oestrus cycle. Likewise, oestrus cycle did not interact or confound the reduced climbing behaviour observed in the LEV study (**Supplementary Fig S19**).

Next, for each experiment, the timepoint with the biggest drop in vertical climbing behaviour was selected for each model, and the data was normalised to the average of the sex-specific control group on the same day, to compare the degree of impairment for the different models. The model with the lowest degree of climbing-impairment, when assessed through duration climbing, was the Multiple Myeloma model, with only 84.8 ± 15.7 % (mean ± S.E.M.), although clearly reflecting the inconsistent development of the model in this experiment. The remaining models were all below 73 %, and most prominently affected models were the systemic models, CAIA (35.5 ± 11.41 %), LEV (35.8 ± 9.6 %), and FD (41.7 ± 12.5 %) (**Fig 11-C1**). Comparing the three experiments with neuropathic pain models suggested that nerve-injuries affecting the hind limb produced more profound impairment than when the nerve injury was located to the orbital nerve, with an average of 48.5-50% for SNI vs 71.2 for FRICT-ION. The top frequency parameter showed a profile comparable to the duration climbing (**Fig 11-C2**), while for the distance moved, the degree of impairment was less prominent, as it ranged from 51.8-95.4 % (CAIA - SNI-1), and of note, for the models primarily producing unilateral plantar or facial sensitivity (CFA, incision, FRICT-ION and SNI) it was only decreased 78-95%. (**Fig 11-C3**).

For all analyses of time course datasets, baseline was always included as a covariate to factor in the variation in the innate desire to climb. Across all climbing-related outcome parameters, the baselines were found to be significant covariates for almost all datasets (except breast CIBP), suggesting that the animal’s pre-model induction natural activity level was still reflected in their behaviour after the induction (**Supplementary Table S4** for overview).

## Discussion

Our study evaluated the utility of vertical climbing activity as a measure of pain-related impairment in naturalistic behaviour. We built a low-cost vertical climbing apparatus (protocol and build guide freely available: https://dx.doi.org/10.17504/protocols.io.n92ldo7q7g5b/v1) and tested ten different pain models with different injury aetiology and characteristics across multiple laboratories. For each model, we compared climbing outcomes with established pain-related endpoints and, where feasible, performed pharmacological validation. Overall, vertical climbing activity came out as a relevant behavioural measure for all models.

We focused on three main parameters, as they bring each their quality in characterising the pain-impaired behaviour. As previous studies exploring climbing-activity, like climbing in the cage-lid [68], on a sloping chainmail [47] or in a vertical cylinder [55], we identified the duration of climbing as a key parameter. In addition, inclusion of the distance moved and the frequency in the top were intended to capture if the painful injury resulted in more cautious movements, which could be seen by a decreased distance, or animals aborting the ascend to the top, indicating an experience of pain associated with the task or that the mouse felt more vulnerable to falling down. Across models, total climbing duration appeared as the most robust marker, as almost all models produced a significant decrease in this parameter, followed by top frequency. The distance moved was less affected by the local injury models, causing pain directly to the paw or face, like the paw incision, SNI, FRICT-ION, and intraplantar CFA models, while it appeared to be a more consistent marker for models affecting the musculoskeletal system, like bone pain and arthritis models (**Table 1**).

**Table 1:**
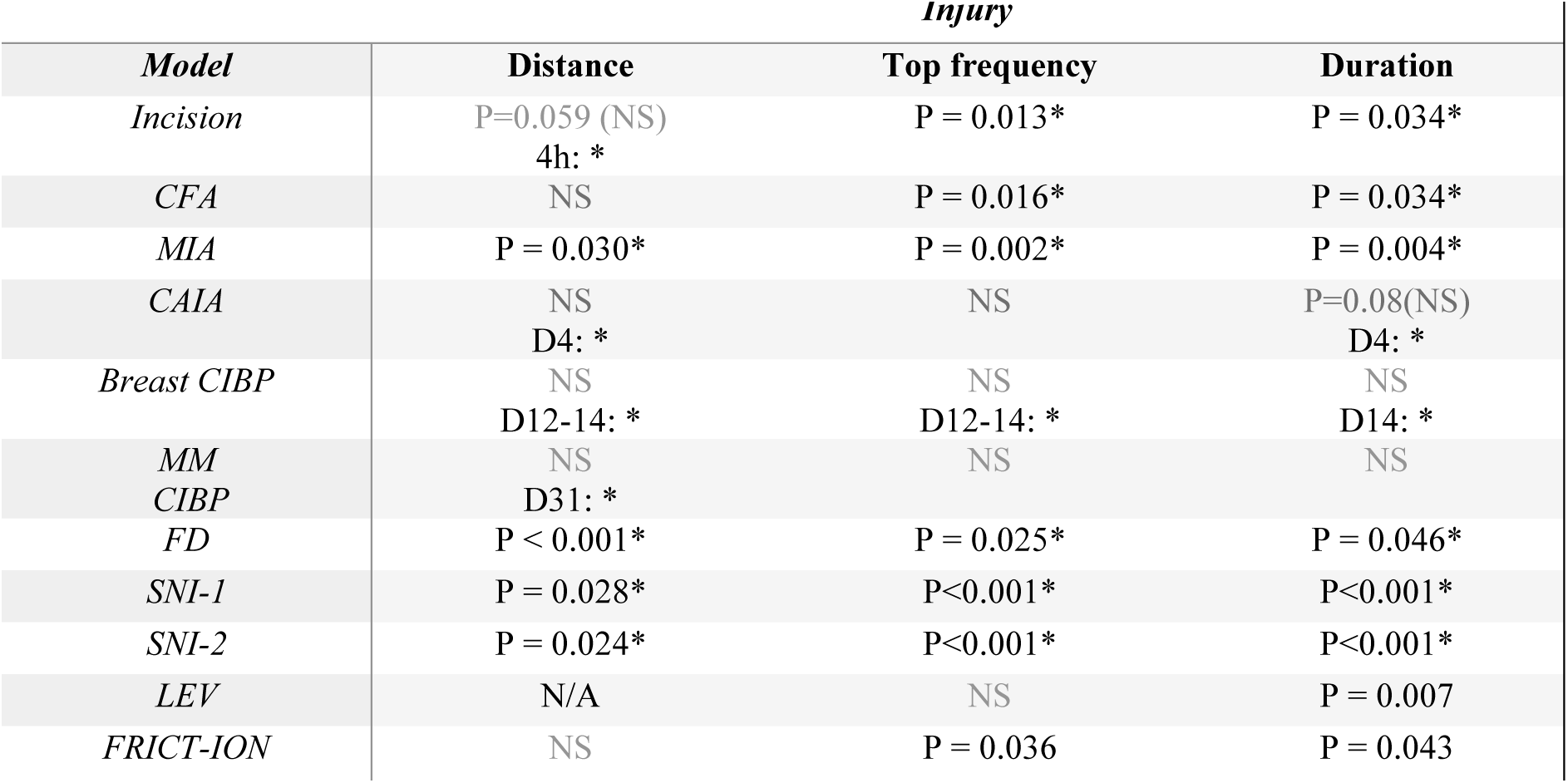
Overview of model-specific results. Summarized data from statistical analysis, presented in full in **supplementary Table S3**. All data was analysed using RM ANCOVA (or mixed effects analysis), factors: sex*injury*time, covariate = baseline. For datasets without significant injury-effects, but significant time*injury interactions, the particular days of significant differences between control and injury group are mentioned, as estimated by Estimated Marginal Means with Bonferroni correction. For results below P=0.1, the P-value was specified, while for P>0.1, they were simply stated: NS = non-significant. *: P<0.05. Models: Incision: plantar incisions model, CFA: intraplantar Complete Freund’s Adjuvant, MIA: Monoiodoacetate Induced Arthritis, CAIA: Collagen Antibody Induced Arthritis, Breast CIBP: Breast Cancer-Induced Bone Pain, MM CIBP: Multiple Myeloma Cancer-Induced Bone Pain, FD: Fibrous Dysplasia, SNI: Spared Nerve Injury (SNI-1: male-female experiment, SNI-2: male-control experiment), LEV: Levcromacalim-induced migraine, FRICT-ION: Foramen Rotundum Inflammatory Constriction Trigeminal Infraorbital Nerve injury.

Because climbing is an elective behaviour, pre-existing motivation and competence are essential. We therefore applied habituation/training sessions and exclusion criteria, and although most of the experiments had relatively low levels of exclusions, two sub-strains (C57BL6/J-BomTac and BALB/cByJRj) showed consistent high exclusion rates and appear poorly suited for this assay. Baseline climbing levels varied substantially between experiments, even when sub-strain, vendor, age, and room conditions were consistent, suggesting experimenter, temporal, or subtle environmental influences. These observations underscore the need for: 1) careful habituation/training prior to study start, with potential additional rounds of habituation for the poor/fearful climbers, 2) analysis of baselines before allocation into groups for exclusion of non-compliant animals and stratified block randomisation according to climbing level, and 3) inclusions of baseline levels as covariates in the statistical analysis.

Curiously, for some models there was a clear disconnect between the temporal dynamics for impaired vertical climbing compared with the traditional symptomatic profile for evoked measures, like mechanical sensitivity. In the levcromakalim-induced migraine model, mechanical hypersensitivity progressively develops from 1 to 2 hours after injection, and with further worsening for each injection-day [9], but for vertical climbing, different temporal characteristics were seen, with the highest severity already at 1 h after the first injection, and no additive effect of later injections. This may suggest that the decreased vertical exploration is not caused by the peripheral hypersensitivity upon grasping the metal mesh, but rather by an aggravation of headache by physical activity, a key diagnostic criteria for migraine [2]. Similarly, in our hands, mechanical allodynia lasts for 3-4 days for the plantar incision model [37], and until day 7 for the CFA model, but for both models vertical climbing was mainly decreased in the acute phase of the injuries, at 4 h or Day 2 for the two models respectively. Additionally, the effect sizes are drastically different when comparing model-induced changes in mechanical sensitivity and vertical climbing, with much smaller effect sizes for the latter. The disconnect in effect size and temporal dynamics likely reflect differences in what each assay probes: evoked tests measure localized nociceptive sensitivity at the injury site, whereas climbing integrates motivation, coordination and the capacity to perform a complex task. Compensation by uninjured limbs, higher inter-individual variability in motivation, and non-pain factors (like anxiety, fatigue, sedation) can all reduce the magnitude and alter the time course of climbing deficits relative to evoked endpoints. Meanwhile, the temporal dynamics and effect size aligned much closer with the other non-evoked measures, like burrowing, weight bearing, limb use or home cage activity measures.

Overall, vertical climbing activity was affected in all models, including the neuropathic orofacial pain model, FRICT-ION, suggesting that the assay is also sensitive to pain-conditions neither affecting the limbs nor known to be aggravated by exercise. Importantly though, comparing with the neuropathic model affecting the motor-system (SNI), the FRICT-ION model produced a lower magnitude of impairment. Generally, the models with the most severe effects on climbing were the systemic models, CAIA, FD and LEV, that cause effects beyond the local injury in one limb. This may naturally reflect the general severity of the models. It also serves as a reminder that mice are highly motivated to climb, meaning that when an injury only affected one limbs, they may simply rely more on the other three legs to maintain the climbing behaviour. For the CAIA model, impaired climbing was seen on the first day after administration of LPS for activation of the immune system, a procedure known to cause severe discomfort [66]. Therefore, it is difficult to ascertain the impact of the rheumatoid-like pathology vs. the discomfort from the LPS. Additionally, this study was also undermined by the high exclusion rate for the BALB/c sub-strain used in this experiment (50 %), limiting the power and preventing clear conclusions in the later stages of this study. Interestingly, similar exclusion rates were reported from a collaborator using the same sub-strain in the climbing cylinder (data not shown), while the BALB/cOlaHSD in our Breast CIBP studies showed consistent climbing compliance. BALB/c lines have previously been characterised to adopt a more defensive behavioural strategy in novel environments, causing lower levels of locomotion and explorative activity in BALB/c than C57BL/6J strains [36], indicating that these characteristics are sub-strain specific.

Pharmacological manipulations provided mixed support for predictive validity. For the plantar incision model, meloxicam only produced a mild trend for restoring the top frequency parameter, suggesting that meloxicam is either insufficient in providing analgesic relief in this model, as suggested by some (preliminary results), or that the low effect size and high variability of the climbing assay have diluted the potential for detecting an effect. For the CFA model, naltrexone reinstated the climbing impairment after CFA recovery, similar to the traditional reinstatement of pain hypersensitivity suppressed by the endogenous opioid system [13]. Lastly, for the SNI model, gabapentin was used to clarify if the impairment in climbing behaviour was due to motor or sensory effects from the injury. Crucially, as most effective analgesics for chronic pain models, like opioids and gabapentinoids, can produce sedative effects [23; 25; 62], the compounds may further decrease, rather than restore, pain-impaired climbing behaviour [55]. Through pilot experiments, we established a non-sedative dose of gabapentin (30 mg/kg), which reportedly also improves mechanical sensitivity in the model [23], and, crucially, we found that it partially restored climbing in SNI-injured mice, while showing no effects in sham-injured. These findings therefore indicate that the assay is sensitive to both pharmacological reinstatement and reversal of pain-induced climbing impairment, supporting the translational relevance of the assay.

With many assays available for measuring climbing activity, the challenge imposed on the animal must be considered. Cage-lid hanging, which entails carrying the entire body weight by the limbs [68], may be too demanding and high-risk for severely affected models to engage in any climbing activity, while something like the sloping chainmail [47] may be less demanding and advantageous under such circumstances. The vertical climbing assay is considered intermediately difficult, and relatively mild models, like plantar incision, may show more prominent and extended declines in climbing activity if tested using the over-head cage-lid hanging paradigm instead. For comparison, SNI-injured mice were down to 46 % of normal cage-lid hanging duration [68], while it was 48-50 % in our vertical climbing assay, and approximately 70 % in the chain-mail assay [47], suggesting a less pronounced decline in the latter setup. A more demanding task may therefore suggest more prominent changes from the injury but may also pose more challenging to reinstate with interventions. We therefore recommend carefully considering the expected model severity and potential implications from the interventions, before selecting which type of climbing assay to employ.

Previous work suggested drastic declines in cage-lid hanging [68] or general home cage activity [28] as mice age, particularly for females. This factor may have caused some of the decline in climbing seen in experiments conducted over an extended timespan, like the MIA study, where the control animals also decreased their climbing prominently over time. This may be partially explained by ageing of the animals, or that reduced novelty from repeated testing results in less explorative activity. Additionally, in both the SNI and CIBP experiments, sham controls also showed prominent drops in the first week after injury, albeit with some recovery following, while this phenomenon was not seen when conducting the SNI experiment with naïve controls. This highlights the need for carefully selecting the type of control-procedure, as sham surgeries in themselves may cause prominent drops in activity.

As recommended [19], sex was always included as a factor in the analysis. Across all studies, the two sexes were not dissimilar in climbing-related exclusions or baseline climbing levels. There were also no effects of oestrus cycle on neither baseline climbing nor climbing after migraine model induction (LEV study). Conflicting results suggest that oestrus cycle may have no effect on general activity [67], induce higher activity during oestrus and metoestrus [10; 34], or that oestrus effects are strain-dependent[40]. Although some pain-related mechanisms and behaviour are sexually dimorphic [42; 43], only one study here showed significant sex effects (the SNI study). Importantly, there was no interaction between sex and injury, suggesting that the injury did not affect the climbing behaviour in two sexes differently. Unfortunately, the studies may not have been sufficiently powered to detect sex*injury interactions, and lack of significant effects does not necessarily exclude the potential of those interactions existing. Of note, it did appear that for some studies, like MIA and SNI, there were more blunted differences between injured and controls in females, also influenced by more prominent declines in activity for control/sham females.

As vertical climbing may be affected by a variety of other factors affecting general motivation and wellbeing, it cannot be considered pain-specific and should be used in combination with other behavioural read-outs probing other aspects of the pain-phenotype. As any other non-evoked naturalistic pain measure, it has high variability between individuals, experiments, and strains, calling for careful inclusion of appropriate controls and careful habituation/training protocols. Lastly, it may be particularly sensitive to other manipulations causing changes in general activity levels, like drug-induced side effects. Here, we focused on the crude location of the animal, but we also noticed that some animals simply adapted the way that they climbed, rather than decreasing their activity. Therefore, more nuance of the behaviour may be obtained through machine-learning-guided mapping of climbing characteristics [7], similar to the recent machine-learning-guided mapping of neuroethological characteristics of stimulus-evoked responses [3; 8].

In conclusion, vertical climbing was impaired in a wide range of pain-related models, across five different laboratories, demonstrating the robustness and consistency of this type of behaviour as a novel pain-related outcome measure. The behaviour correlated significantly with most other pain-related measures, although the temporal dynamics differed from those of evoked mechanical allodynia, while being more closely aligned with other non-evoked behaviours, like burrowing, home cage activity, running wheel activity, or weight bearing. Notably, we validated the assay through pharmacological interventions, either providing pain relief (gabapentin) or pain reinstatement upon recovery (naltrexone). As many other non-evoked naturalistic behaviours, it may suffer from high individual variability and lack pain-specificity, although it may possess higher translational value for the clinical symptoms seen in patients. Altogether, this indicates that vertical climbing behaviour has broad suitability as an outcome measure in preclinical pain research.

## Conflict of interest statement

please see supplementary **Table S5** for full disclosure of conflicts of interest and funding. Several authors are affiliated with and/or co-founders of companies developing non-opioid drugs for treatment of pain (Regulonix LLC and Zyneyro), but as the present experiments are independent and unrelated to their research and development activities, this is not considered a conflict of interest.

## Acknowledgements

We wish to thank Trine Marie Ahlmann Glahder for invaluable technical assistance. We also wish to thank the Rodent Metabolic Phenotyping Platform at the Novo Nordisk Foundation Center for Basic Metabolic Research (CBMR) for letting us use their behaviour rooms and running wheels for the MIA and CAIA experiments. We wish to thank Camilla Skånstrøm Dall, Mathilde Caldara, Sascha Agergaard Martinussen, and Natascha Synnøve Olsen for their technical support.

## Author contributions

MEM, TB and SH developed the vertical climbing assay for these experiments. SH coordinated and designed experiments with all involved parties. MEM, TB, EC, HNA, SLC, KOR, CH, AMH, KLM, ATS, KSPA and SH conceptualized studies. MEM, TB, EC, OCJ, CMG, IBK, HNA, PCJ, KLJ, HDG, YT, KOR, CH, SLC and SH designed experiments for the different models. MEM, TB, EC, PCJ, JJJ, CMG, IBK, HNA, NKG, KLJ, HDG, YT, KWEM, AS, ATD, SLC and SH conducted experiments. SLC, KOR, AMH, KLM, ATS, RK, KSPA, OK and SH supervised experiments. TB, EC, JJJ, HA, YT, KWEM, AS, ATD, KOR, AH and SH analysed results. MEM, EC, JJJ, HNA, YT and SH, drafted model-specific methods sections. OK provided statistical guidance. SH merged all methods, stratified and performed statistical analysis and wrote the first collected draft of the manuscript. All authors then had the opportunity to discuss the results and comment on the manuscript. Find specified author-contributions and financial support in supplementary **Table S5-6**.

## Data availability statement

All data is presented in the manuscript or supplementary files and can be made available upon request to the corresponding author (SH).

## Notes

https://www.protocols.io/view/vertical-climbing-assay-for-mice-n92ldo7q7g5b/v1

### Funding

- Lundbeck Foundation: LBF R467-2024-544, R389-2021-1596
- European Union: BonePainIII Grant No. 101120524
- Carlsberg Foundation: CF23-1620
- Independent Research Fund, Denmark: 3162-00044B
- Innovation Fund Denmark: Industrial PhD 3194-00050B
- Center for Applied Laboratory Animal Research, Denmark

