## Supplementary file for "Vertical climbing in mice: a translationally relevant measure of pain-related impact on natural behaviour"

1. **Full methods**

**1.1. Animals**

All the experiments were performed in accordance with animal experimentation licenses, granted by either the local IACUC at University of Florida (FRICT-ION: IACUC protocol 202400000002), or the Danish Animal Experiments Inspectorate:

- Bone pain models: 2025-15-0201-01862
- Fibrous Dysplasia: 2021-15-0201-00872 and 2026-68838
- CAIA: 2025-79831
- MIA: 2025-79427
- Plantar Incision: 2024-15-0201-01627
- SNI+CFA: 2021-15-0201-01036
- LEV: 2022-15-0201-01347

Animals were always acclimatized for at least one week after arrival from the external vendor, before experiments were initiated, and housed in groups, unless otherwise specified. They were provided ad libitum food and water, given housing- and nesting material, gnawing sticks and provided with additional enrichment when feasible. Specifics for the various stable conditions below:

**Supplementary Table S1**. Technical overview of animal-related experimental details.

| **Exp.** | **Age (weeks)** | **Sex** | **Strain** | **Vendor** | **n** (planned) | **Cage density** (co-housing of control and injured) | **Induction analgesics** | **Exclusions due to baseline climbing** |
|---|---|---|---|---|---|---|---|---|
| Plantar incision<br><br>+ Melox. | 8 | Male<br>Female<br><br>Male | C57Bl/6JRj | Janvier | 9-12 (12) | 4 (mixed) | None | 2/24<br>1/24<br><br>3/24 |
| MIA | 8 | Male + female | C57Bl/6JRj | Janvier | 8-11 (8-12) | 4-5 (separated) | None | 0/20<br>1/20 |
| CAIA | 8 | Male + female | BALB/cByJRj | Janvier | 2-5 (8) | 2 (separated) | None | 12/24<br>12/24 |
| SNI | 8 | Male<br>Female | C57Bl/6NRj | Janvier | 7-10 (10) | 5 (mixed or separated) | Bup. 0.1 mg/kg s.c.,<br>+ Lido/Bupi<br>+ 3 days Carprofen (5 mg/kg) | 3/20<br>1/20 |
| SNI – control exp | 8 | Male | C57Bl/6NRj | Janvier | 10-12 (12) | 5-6 (separated) | Bup. 0.1 mg/kg s.c.,<br>+ Lido/Bupivacaine<br>+ 3Days Carprofen (5 mg/kg) | 2/24 |
| CFA-intra-plantar | 10 | Male<br>Female | C57Bl/6NRj | Janvier | 5-6 (6) | 6 (separated) | Bup., 0.1 mg/kg s.c. | 0/12<br>1/12 |

| | | | | | | | | |
|---|---|---|---|---|---|---|---|---|
| Migraine model | 7-13 | Male + Female | C57BL/6J BomTac | Taconic A/S | 8-9 | 2-6 (mixed) | None | 6/22 7/25 |
| Breast cancer bone pain | 6-7 | Female | BALB/ cOlaHSD | Envigo | 7-9 (9) | 4-5 (mixed) | 5 mg/kg carprofen, pre- + D1 + D2 | 0/18 |
| Multiple myeloma bone pain | 4-5 | Male | C57BL/ KaLwRijHsd | Envigo | 7-9 (9) | 3-4 (mixed) | 5 mg/kg carprofen, pre- + D1 + D2 | 1/18 |
| Fibrous Dysplasia | 12-13 | Male | Transgenic $G\alpha_s^{R201C}$ mice x B6.Cg-*Gt(ROSA)26Sor*$^{tm1(rtTA,EGFP)Nagy}$/J | In house breeding* | 5-7 (5-7) | Single | None | 0/12 |
| Trigeminal neuropathic pain | 10 | Male + female | C57Bl/6J | The Jackson Lab | 4-8 (7-8) | 5 (mixed) | none | 1/15 4/15 |

*Originally obtained from The Jackson Lab but maintained in our own facility.

##### 1.1.1. Plantar incision studies

Male and female specific pathogen free C57BL/6Jrj mice (Janvier; Le Genest-Saint-Isle, France), aged 8 weeks on arrival, were housed in a facility accredited by the Association for Assessment and Accreditation of Laboratory Animal Care (AAALAC) at the Faculty of Health and Medical Sciences, University of Copenhagen. Upon arrival, mice were ear-marked and housed in individually ventilated cage (Blue Line Type III 1290, Tecniplast S.p.A, Buguggiate, Italy) at a density of four animals per cage. They were then left undisturbed for seven days. After acclimatisation, we transferred the mice to the experimental room for behavioural testing and housed them inside a ventilated cabinet ("Scantainer", SCANBUR, Karlslunde, Denmark), without the plastic lid. Four mice (two incision and two control animals) were housed per cage. The mice were housed on soft cellulose bedding (ALPHA-dri, Shepherd Specialty Papers, Watertown, TN, USA), with clear tunnels (50 x 3 x 100 mm Clear Handling Tunnel, Datesand, Bredbury, UK), 2-4 aspen gnawing bricks (Chew Block Small, Tapvei, Harjumaa, Estonia), red transparent plastic shelters (JAKO, Molytex, Glostrup, Denmark) and nesting material (10 g Carfil Nesting Cup, SCANBUR). Mice had *ad libitum* access to water and food pellets (Altromin 1314, Altromin Spezialfutter GmbH & Co. KG, Lage, Germany) throughout the study. The facility maintained a temperature of 22 °C (± 2 °C) with a humidity of 55 % (± 10 %), and the cages were ventilated at 75 air changes per hour. Animal rooms were on a 12/12-hour light/dark cycle with a 30-minute twilight-phase, with lights at 50 % intensity from 5:30, increasing to 100 % at 6:00, and decreasing to 50 % at 17:30 before turning off at 18:00. The experimental room was maintained on a 12/12-hour light/dark cycle without the twilight phase. When possible, the mice were tunnel handled, as a standard procedure for the facility.

The Danish Animal Experiments Council approved all procedures (license 2024-15-0201-01627) and the study was overseen by a local animal welfare body. We defined humane endpoints before initiating the study. They included pronounced dehydration, lethargy, infection of the incision wound, weight loss above 20 %, and blood in urine, mouth or faeces. In addition, we used a scoring system twice daily to monitor acute and accumulated lameness with acute severe lameness and moderate lameness lasting more than three days being humane endpoints. No animals reached the predefined humane endpoints.

##### 1.1.2. Arthritis studies (CAIA and MIA):

The mice were housed in an AAALAC (Association for Assessment and Accreditation of Laboratory Animal Care) accredited, Department of Experimental Medicine, Faculty of Health and Medical Sciences at the University of Copenhagen, Denmark. Environmental conditions were maintained at a controlled temperature of 20-22°C and a relative humidity of 55±10%. A standardized artificial circadian rhythm was applied, consisting of a 12-hour light/12-hour dark cycle, with lights on from 6:00 to 18:00 and lights off from 18:00 to 6:00.

Mice were housed in individually ventilated cages (IVC; Techniplast, Emerald Type II, EM500) and provided with ad libitum access to food (SAFE D30, Scientific Diet) and water. During Day 3-9 following LPS injection (Day 3) in the CAIA studies, food pellets were softened in water and placed on the cage floor to ensure easy accessibility and were replaced every morning and evening during this period. For the CAIA-experiments, mice were housed in pairs, while for the MIA-experiments they were housed in groups of 4. In both cases, animals were always housed with animals from the same intervention-group.

Environmental enrichment was provided in all cages and included nesting material (Cardboard Mouse House, Carfil nesting cup, 10 g; Carfil Quality, Belgium), gnawing bricks (S-brick, 50 mm x 10 mm x 25 mm; Brogaarden, TAPVEI), and bedding material (Wood shavings, Brogaarden). For the CAIA-experiments, running wheels were included in the cages. A clear handling tunnel (Datasand, Scanbur) was used during handling when possible, facilitating animal observation while reducing handling-related stress and anxiety as well as the risk of bites for both animals and handlers.

Predefined humane endpoints were assessed throughout the studies, following a previously used protocol[9; 13]. Humane endpoints focused on weight loss (max. 15%), fur quality, posture and changes in general behaviour.

##### 1.1.3. Bone pain studies:

Animals were kept in a specific pathogen-free animal facility at the Department of Drug Design and Pharmacology, University of Copenhagen, Denmark. Environment of the housing room was regulated at 22 ± 2˚C room temperature, 55±10% humidity, and 60% white light intensity with 12h/12h light dark cycle (light cycle: 7 AM – 7 PM). Female mice were housed in groups of 4-5 per cage, while male mice were housed 3-4 per cage. Mice were housed in individually ventilated transparent GM500 cages (Techniplast, Italy) enriched with sawdust bedding (Tapvei 2HV, Brogaarden, Denmark), wooden gnawing block (Aspe småklodser, Brogaarden, Denmark), small red translucent tube (Polycarbonate Mouse Tunnel; Datesand, UK), small red housing unit (Datesand, UK), paper ropes (Diamond twist, Envigo, Italy), and paper nesting (paper shavings; Brogaarden, Denmark). Mice received standard chow (Altromin 1314F, Brogaarden, Denmark) and tap water ad libitum. Cages were changed biweekly, while food and tap water were replenished weekly. Mice were acclimatized to the facility and experimenter at least 7 days prior to the behavioural habituation and baseline assessment. When possible, the mice were tunnel handled, as a standard procedure for the facility.

For health monitoring, a predefined severity scoring system was designed to monitor mice body weight, coat condition, body function, and behaviours. Monitoring was conducted throughout the experiment starting on baseline measurements until reaching humane or experimental endpoint. The humane endpoint was reached if a mouse scored at least 6 on the severity risk assessment or reached limb use 0 following model induction. The experimental endpoint was reached when all the cancer mice had been euthanized. All the mice were euthanized by cervical dislocation at both endpoints.

##### 1.1.4. SNI and CFA studies:

Wild type male and female C57BL/6Nrj mice (aged 8 weeks at the beginning of experiment) (Janvier, France) were housed (4-6 per cage) at an AAALAC (Association for Assessment and Accreditation of Laboratory Animal Care) accredited Animal Core Facility Unit at Faculty of Health and Medical Sciences, University of Copenhagen, Denmark. In the housing room temperature was kept at 22±2°C and humidity was kept at 55%±10% with a 12/12 hour light/dark cycle (6AM-6PM) with free access to standard rodent chow (Altromin 1342 (Brogaarden, Denmark) and water. Cages were enriched with sawdust bedding, nesting material, a wooden stick, a transparent tunnel for tunnel handling and a transparent red cover. Mice were initially handled by tunnel handling and gradually trained to be handled by scooping. Animals were allowed at least seven days of habituation to the animal facility before initiation of experiment. Mice were monitored at least once weekly (SNI) or once daily (CFA) after model induction by a severity scoring system looking at body weight, fur quality, posture and changes in general behaviour. The humane endpoint was defined as reaching a score of 4. No animals met this criterion. Animals were sacrificed by cervical dislocation when reaching the experimental or humane endpoint.

##### 1.1.5. Mouse model of levcromakalim-induced migraine studies:

Experiments were performed at the Translational Research Centre, Copenhagen University Hospital – Rigshospitalet, Glostrup, Denmark. Mice were male and female C57BL/6J BomTac (Taconic, Denmark) delivered at seven weeks of age and housed in groups of two to six individuals. Upon arrival at the facility, mice were acclimatized at least one week prior to habituation for the experiments. Mice were housed under a 12-hour light/dark cycle, lights on at 7 AM in European standard open cages (measuring 45x35x20 cm) with free access to food (Premium Scientific Diet, SAFE Lab) and tab water. Temperature was controlled between 21 and 23 degrees Celsius. Cages were enriched with two types of nesting materials and multiple plastic and cardboard shelters and tunnels on the floor and attached to the lid. Mice were tunnel handled. At the start of experiments, animal age ranged from 7-13 weeks, weight 17.7-24.4 g (females) and 25.0-35.1 g (males).

None of the test subjects in this study were euthanized for any disease related nor experimental causes. The experiments were approved by the Danish Animal Experiments Inspectorate and performed in accordance with the relevant guidelines and regulations of approval number 2022-15-0201-01347.

#### 1.1.6. FRICT-ION studies:

Male and female C57Bl6/J mice from Jackson Labs (Bar Harbour, ME, USA) were group housed 5 animals per cage in ventilated cages with ad libitum access to food and water in the McKnight Brain Institute Animal Facility at the University of Florida, Gainesville, FL, USA. Animal were maintained in housing rooms with a 22°C room temperature and a 12-hour light:dark cycle, with lights on at 6:00 and lights off at 18:00. Cages consisted of corncob bedding, nesting material, and a transparent tunnel. Animals were habituated to the housing room for at least 7 days prior to experimental habituation. Animals were 10 weeks old at the start of testing. Following surgery, mice were monitored daily for body score and weight loss. No animals were excluded from the study for surgical-related complications. Sham and FRICT-ION mice were housed together.

### Model-inductions and study design:

#### 1.1.7. Post-operative pain model (Plantar incision):

Mice were anaesthetized with isoflurane. Mice were individually placed in a sliding top chamber (Kent Scientific) with 5 % isoflurane delivered in 100 % oxygen (flow rate of 500 ml/min) until loss of the righting reflex, after which we maintained anaesthesia with ~1.9 % isoflurane delivered in 100 % oxygen via a face mask for spontaneous breathing (flow rate 100 ml/min) while placed on a heat mat. The right hind paw was cleaned and disinfected *lege artis* with 4 % chlorhexidine soap and 70 % ethanol. No perioperative analgesia was administered in the characterization study, as the study aimed to assess pain-related outcomes, and analgesic intervention would be a confounding factor in the experimental model. In the validation study, all mice received an injection subcutaneously with either 10 mg/kg meloxicam diluted to 0.01 mg/mL in PBS, or PBS. Injections were administered under anaesthesia prior to the incision.

The plantar incision was performed as described by Cowie et al.[5] with minor changes. After preparation, the skin on the plantar surface was incised with a 4 mm longitudinal incision through the skin with a #11 surgical blade. The flexor digitorum brevis muscle was lifted with curved forceps and split in half longitudinally. The area was moistened with saline, the muscle was returned to the wound cavity, and the skin was sutured with two interrupted sutures (Ethilon 6-0 nylon monofilament, ref. 697H, Ethicon Inc., Cornelia, GA, USA). Anaesthesia-only mice, termed "control", had their paw cleaned and disinfected under anaesthesia and were kept unconscious for 13 minutes; the average time to complete a surgery. The procedures were conducted at 9:00-12:00. The procedure was performed under fully aseptic conditions by an experienced surgeon (MEM) with a non-sterile assistant (TB).

Following the procedure, mice were allowed to recover independently in a cage halfway underlined with a heat mat (1180 x 280 mm - 62W, LP Racks, Juelsminde, Denmark) until walking, before being returned to the home cage with conspecifics.

The experiment was divided in sex-separated consecutive cohorts of 12 animals per cohort, including equal numbers from each experimental group. For the initial experiments, climbing behaviour was assessed at pre-injury baseline and approximately at 4h, 24h, 48h, 74h and 96h after surgery in control and incision male and female mice, to characterise the impact of the model on climbing behaviour (**Fig S1A**). A follow-up experiment was then conduced in male incision animals alone, to test the effects of pre-surgically administered meloxicam (10mg/kg) on climbing behaviour at 4h after injury (**Fig S1B**). All behavioural tests with this model were conducted by the same male experimenter (TB), who was blinded to group-allocation. The experiment was terminated by providing mice with isoflurane anaesthesia until unconscious followed by cervical dislocation.

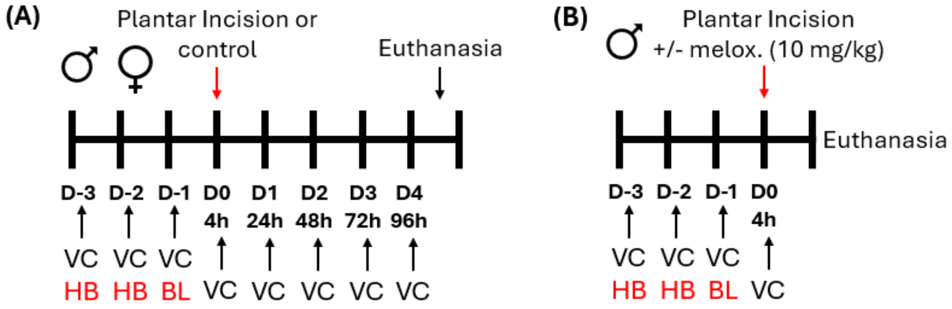

**Fig S1:** Study design for studies involving plantar incision. A: Characterisation study including both males and females. B: meloxicam study, including only males. Day 0 = induction of injury. VC = vertical climbing. HB = habituation. BL = baseline. Melox. = meloxicam.

Exclusions: Based on pre-defined exclusion-criteria on climbing baseline levels, two females (incision group) and one male (control group) were excluded from the initial characterisation experiment. From the following meloxicam study, 3 males were excluded (vehicle group). As the climbing analysis was only performed after the study-completion, baseline behaviour was not included in group allocation, and therefore unfortunately affected the groups unequally. Additionally, due to unfortunate events, some video-files were compromised or deleted by human mistakes or technical errors, leading to missing recordings for the characterisation study: 48h: 2 females, 72h: 13 females.

#### 1.1.8. Inflammatory pain model – intraplantar CFA

To model inflammatory pain, we used the intraplantar CFA model [22]. In brief, mice were deeply anesthetized with isoflurane (2%) and received an intraplantar injection of CFA (7 µL, Sigma Aldrich, F5881) using a Hamilton syringe with a 30 G needle and were given a single injection of Temgesic (buprenorphine, 0.1 mg/kg) for post-procedure care. Control animals were exposed to anaesthesia and analgesic treatment with Temgesic, without intra-plantar injection. The experiment was conducted in two different cohorts, one pr each sex, and all animals in a cage were allocated to the same treatment-group after confirming equal levels of climbing performance (duration climbing) between cages.

C57Bl/6N mice, aged 10 weeks, housed in groups of 6 pr cage, were habituated to the climbing apparatus and von Frey testing chambers prior to Complete Freund's Adjuvant (CFA) induction. Climbing habituation consisted of three consecutive days: on Day −7 (D−7), all mice from each cage were placed together in the climbing apparatus for 45 minutes; on Day −6 (D−6), individual mice were placed in the climbing chamber for 15-minute sessions; and on Day −3 (D−3), a 15-minute baseline climbing recording was obtained, provided that all mice demonstrated appropriate climbing behaviour. Climbing recordings were performed 2 mice at a time in a single room.

For von Frey habituation, mice were first placed in individual cups on a wire mesh platform for 1 hour on Day −7 (D−7). On Day −6 (D−6), mice were exposed to the von Frey filaments while on the grid, and on Day -5 (D-5) baseline mechanical sensitivity thresholds were measured using the ascending method.

Following baseline measurements, the model was induced (explained above), and mechanical sensitivity and climbing behaviour were assessed on Day 2, 4, 7 and 11 post-CFA injection. For the female-cohort, climbing behaviour was also recorded on Day 22 1 hour after s.c. injection of naltrexone (3 mg/kg, Sigma, N3136-250mg, lot BCBX4989) to re-establish pain-related responses. All behavioural tests with this model were conducted by the same female experimenter (CMG), who was blinded to group-allocation.

At the end of the experiment, mice were euthanised using cervical dislocation.

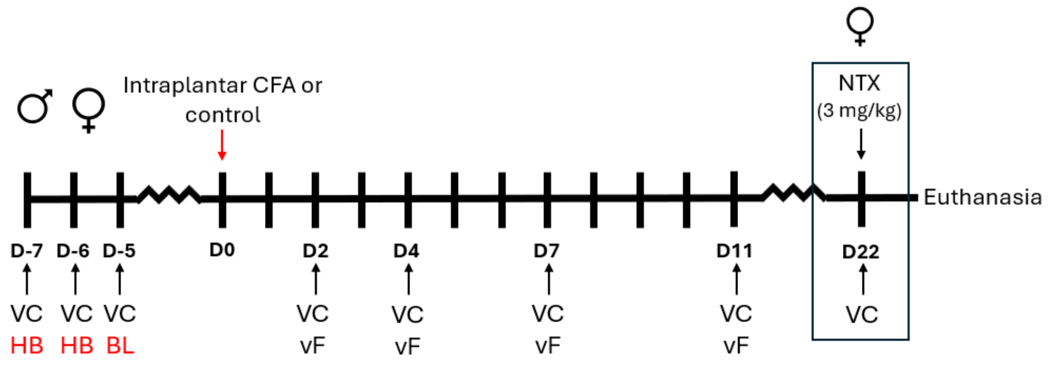

**Fig S2.** Study design for studies involving the intraplantar CFA-model. Day 0 = induction of injury. VC = vertical climbing. HB = habituation. BL = baseline. vF: von Frey measure (mechanical allodynia). NTX = Naltrexone, which was only conducted in the female cohort.

#### 1.1.9. Osteoarthritis model - Monoiodoacetate Arthritis Model (MIA).

Induction of the Monoiodoacetate Arthritis Model to the knee joint (MIA) was performed similar to previously reported [10-12]. On the day of injection, the monoiodoacetate solution was freshly prepared in sterile saline for injection of Sodium Iodoacetate (Sigma, I2512) in saline. Animals were anaesthetized in an induction chamber using Isoflurane 2.5% mixed in $O_2$ at a flowrate of 1 L/min and maintained via facemask at 2.0% during the injection. Anaesthetic depth was confirmed by lack of withdrawal reflex to a pinch to the tail. The animal was then placed on its back in dorsal recumbency, and the fur was shaved in the area around the knee on the left hindleg. The knee was stabilized and fixed in a slightly bend position and the patellar tendon was visualized as a white line below the skin. Using a 30G insulin-syringe (BD Micro-Fine Plus Demi, 0.3ml (30G)) the injection of Sodium Iodoacetate (1mg/10µl) was made intraarticularly in the joint-space by applying it perpendicularly through the skin and tendon just below the patella. Control animals were anaesthetized and shaved around the knee but not given any injections. Model-induction was performed by a female experimenter experienced with the particular model (SH).

The experiment was conducted on equal numbers of male and female C57BL/6Nrj in one cohort including a total of 40 animals, housed in groups of 4. The mice were allocated to group pr cage as follows: Controls: n=8, MIA: n=12, to account for potential loss of animals due to MIA-induced toxicity. Vertical climbing behaviour was assessed at pre-injury baseline, and then at day 3, 7, 10, 14, 20, 28, 36 and 42. Additionally, Dynamic Weight Bearing (DWB) was assessed on Day 8/9, 15/16 and 30/31, but due to the high number of animals, this was done over two days, divided pr sex. All behavioural experiments with this model were conducted by the same female experimenter (HDG), who was blinded to group-allocation.

Exclusions: One male animal designated for the MIA-group, was excluded at baseline due to reaching the preset exclusion-criteria for the climbing behaviour, as he climbed to the top of the cylinder, and remained there for the entire duration of the test. One female mouse showed signs of MIA-induced toxicity immediately after the induction of the model, and was therefore euthanized immediately, as stated by our predefined model-related humane endpoints.

At the end of the experiment, mice were anaesthetized with Isoflurane until unconscious and euthanized using cervical dislocation. Following dissection of the knee-joint, damage was visually confirmed for all MIA-animals.

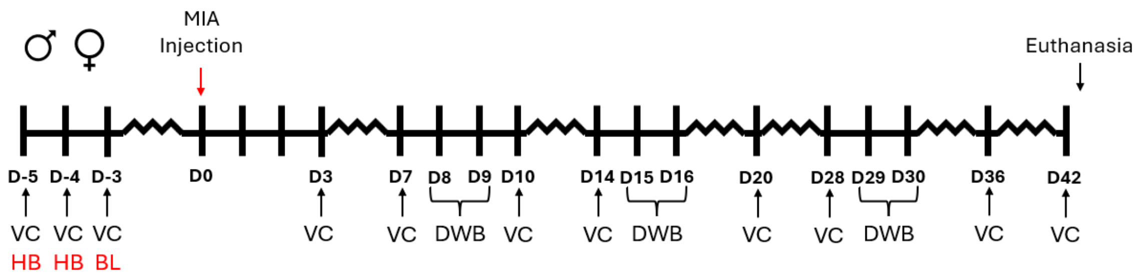

**Fig S3:** Study design for studies involving the MIA-model. Day 0 = induction of injury. VC: vertical climbing. HB: habituation. BL: baseline. DWB: Dynamic Weight Bearing was performed over two days, - males one day, females the other.

#### 1.1.10. Rheumatoid arthritis model - Collagen Antibody Induced Arthritis (CAIA)

To model Rheumatoid Arthritis, we used the CAIA-model [1]. In brief, on Day 0, 1.5 mg (in 0.15 mL saline) mouse monoclonal antibodies against type II collagen (Arthrogen-CIA Arthritogenic Monoclonal Antibody 5-clone cocktail, Chondrex Inc., Woodinville, WA, USA) was injected intravenously (i.v.) in the tail vein by an animal caretaker. The controls were injected i.v. with 0.15 mL saline. On Day 3, one experimenter injected mice of the CAIA groups with LPS (Chondrex Inc., catalog nr.: 9028) from *E. coli* via intraperitoneal (i.p.) route of administration, either 5 or 10µg (in 0.1 mL) for the "low-LPS" vs "high-LPS" groups respectively. Controls were injected with 0.1 mL saline.

The experiment was conducted in four consecutive sex-separated cohorts of BALB/cByJrj mice, including 12 animals pr cohort, with four subjects from each of the three groups (control, low-LPS and high-LPS). Animals were housed in pairs of the same injury-group. All cohorts were assessed on climbing behaviour at pre-induction baseline, and Day 4, 8 and 14 after model-induction. Additionally, burrowing behaviour was assessed on Day 6 and 13, and arthritis severity scores and animal welfare scores were monitored throughout. All behavioural experiments with this model were conducted by the same female experimenters (PCJ, KWEM), who were both blinded to group-allocation.

Exclusions: One female was euthanized on Day 6 due to predefined model-related humane endpoint. A total of 48 mice were included in the study (24 of each sex), and half (24 animals), were excluded due to predefined exclusion-criteria on climbing baseline behaviour. The exclusions were equal for males and females. For burrowing and arthritis scores, only data from animals included on climbing behaviour, are presented for this model. At the end of the experiment, mice were euthanised using cervical dislocation.

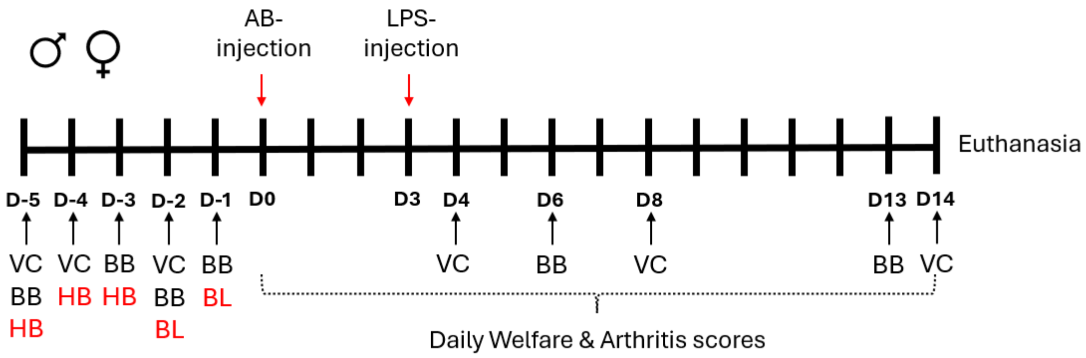

**Fig S4:** Study design for studies involving the CAIA-model. Day 0 = induction of injury. VC: vertical climbing. BB: Burrowing behaviour. AB: antibody. HB: habituation. BL: baseline.

The climbing assessment in the CAIA-model was obtained as a part of a larger study, also including home cage measures, gut microbiota analysis, biochemical- and histological measures and a very detailed welfare and severity scoring. Therefore, only representative data from the animals that passed the exclusion-criteria for the climbing assay, is presented here for supporting the understanding of the disease model in the context of climbing behaviour, while the remaining data will be presented separately in a future publication.

#### 1.1.11. Cancer-induced bone pain models

Cells culture: All the cell lines were cultured in a sterile environment at 37°C with 5% CO2. Both cell lines were subcultured at a ratio of 1:4 to 1:5 upon reaching 80% confluency.

4T1-Luc2 Cells: 4T1-Luc2 cells (ATCC, Manassas, VA, US) were cultured in phenol red-free RPMI 1640 (#32404014, Gibco) supplemented with 10% FBS (04-007-1A, Biological Industries) and 1x Pen-Strep-Glutamine (10-378-016, Gibco). The cells were cultured for approximately two weeks prior to the surgery. Two days prior to model induction, the cells were subcultured with at ratio of 1:4 to ensure log phase growth. On the day of tumour cell inoculation, cells were quickly detached with 0.5% trypsin-EDTA solution at 37°C and resuspended in Hank's Balanced Salt Solution (HBSS, 15175-053, Gibco) at a density of $10^6$ cells/mL. Cells were kept on ice throughout the surgery.

5TGM1-GFP Cells: 5TGM1-GFP cells were cultured in suspension in RPMI 1640 media containing GlutaMAX-l (61870, Gibco) supplemented with 1% penicillin/streptomycin (10000 U/mL, 15140122, Fischer Scientific), 1% sodium pyruvate (1 mM, 11360, Gibco), 1% MEM nonessential amino acids (100 mM, 11140, Gibco), and 10% FBS (04-007-1A, Biological Industries). Two days prior to model induction, the cells growth was boosted by adding a fresh warm medium 1/3 of the total culture volume. On the day of tumour cell implantation, cells were harvested and resuspended in HBSS to a density of 5 x $10^6$ cells /mL. Cells were kept at room temperature throughout the surgery.

CIBP model induction: Cancer-induced bone pain models were induced via intrafemoral inoculation of cancer cells into right hind legs as described before[7; 8; 15]. Mice were initially anesthetized with a mixture of 43 mg/kg of ketamine (Ketaminol, MSD Animal Health, The Netherlands) and 6 mg/kg of xylazine (Rompun Vet, Bayer, Germany), i.p. Mice then received eye ointment (Ocry-Gel, Domes Pharma, UK), 5 mg/kg carprofen (s.c, Carprosan Vet, Dechra, The Netherlands) as pre-surgical analgesia, and 0.5 mL of 0.9 % saline (s.c) to prevent dehydration. Mice were maintained under 2% isoflurane and kept on the heating table at 37°C throughout the surgery.

Following the loss of pedal reflex, the right hind leg was shaved with veterinary clippers and disinfected with 70% ethanol. A small horizontal incision was made over the patella tendon. The lateral side of the patella tendon was loosened, and patella tendon was laterally displaced, allowing access to distal femoral epiphysis. A hole was then made in the femoral epiphysis into intramedullary femoral cavity with 30G needle, in which needle placement was verified via X-ray. Using a 0.3 mL insulin needle, 10 µL of either vehicle (HBSS) or cell suspension were then inoculated into the cavity. The hole was sealed with bone wax (Ethicon, US) and irrigated with a sterile 0.9% saline. The patella tendon was repositioned to original position, and the incision was closed with surgical clips.

Post-operative analgesia was maintained with 5 mg/kg carprofen (s.c) on days 1 and 2. Mice were subjected to daily severity assessment and wound monitoring for four days. Medical clips were removed from all mice at the end of day 4.

#### 1.1.12. Breast CIBP – study design

Eighteen female BALB/cOlaHSD were obtained at 6-7 weeks old, inoculated with tumour cells at 8-9 weeks old, and subjected to behavioural assessment starting from 9-10 weeks old. Prior to baseline behavioural assessments, mice were habituated and trained to the weight bearing procedure from D-6 to D-3 and vertical climbing at D-3. Two baseline behavioural assessments were taken for limb use at D-6 and D-5, weight bearing at D-2 and D-1, and vertical climbing at D-3 and D-2. Following the last behavioural baseline sessions, mice right hind legs were imaged with X-ray for baseline with IVIS Lumina XR Optical Imager (Caliper Life Sciences).

Prior to tumour cell inoculation, mice were randomized based on the body weight and weight bearing baseline ratios into sham (n=9) and cancer (n=9) group. Mice were then inoculated with 10.000 cells/10 µL of 4T1-Luc2 cells (or HBSS for sham group) at D0, followed by post-surgical care from D1-D4. Starting from D6, mice were subjected to limb use and weight bearing every day and vertical climbing every second day until reaching humane or experimental endpoint. To verify disease burden, mice were imaged again with X-ray at terminal end point to determine the relative bone density on the proximal femur as described before[15]. Behavioural assessment and tumour cell inoculation were performed by a single male experimenter (EC), who was masked throughout on the experimental group randomization. All the behavioural assessments were performed during light cycle (7 AM – 7 PM).

Exclusions: Two mice from the cancer group were excluded due to tendon displacement following tumour cell inoculation, resulting in n=7 in the cancer group. Additionally, one animal from the sham group was excluded from this dataset due to the pre-defined exclusion-criteria on baseline climbing activity.

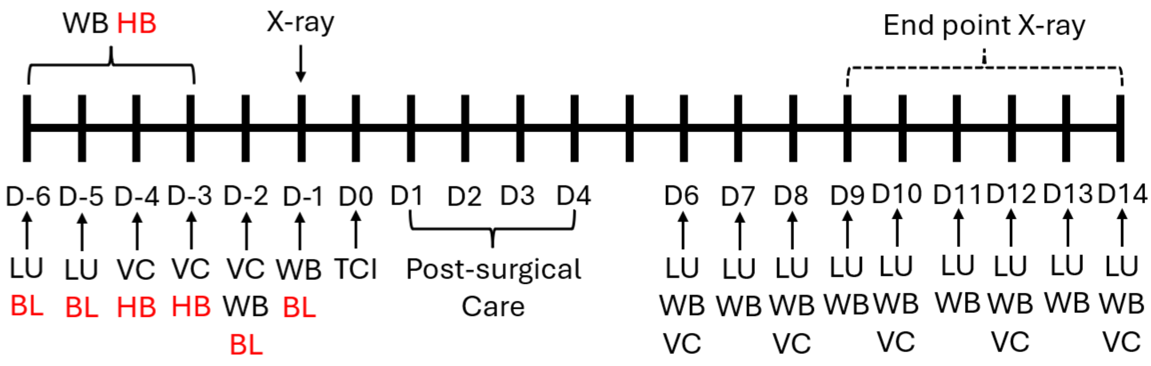

**Fig S5.** Experimental design and timeline for breast cancer bone pain model. *LU: Limb Use, WB: Weight Bearing, VC: Vertical Climbing, BL: Behavioural Baseline, HB: Behavioural Habituation, TCI: Tumour Cell Inoculation.*

#### 1.1.13. Multiple Myeloma CIBP - study design

Eighteen male C57BL/KaLwRijHsd mice were obtained at 4-5 weeks old, inoculated with tumour cells at 6-7 weeks old, and subjected to behavioural assessments starting from 8-9 weeks old. After facility and experimenter acclimatization, mice were habituated with vertical climbing at D-6. Two baselines recording were taken for limb use at D-2 and D-1 and vertical climbing at D-5 & D-4.

For tumour cell inoculation, mice were randomized based on the body weight and into sham (n=8) and cancer (n=10) group. Mice were inoculated with 50.000 cells/10 µL of 5TGM1-GFP cells (or HBSS for sham group) at D0, followed by post-surgical care from D1-D4. Starting from D8, mice were subjected to behaviour assessment as shown in **Figure S6**. Limb use was assessed at D8, D11, D15, D17, D19, D21, and every day from D23 until reaching humane or experimental endpoint. Vertical climbing was assessed at D8, D11, D15, D19, D21, and every other day from D23 to humane or experimental endpoint. To verify the disease burden, spleen tissue was collected from each mouse and weighed (data not shown). Limb use assessment and tumour cell inoculation were performed by experimenter (EC), while vertical climbing was performed by another experimenter (AD). Both experimenters were masked throughout the experiment from experimental group allocation. All the behavioural assessments were performed during light cycle (7 AM – 7 PM).

Exclusions: One mouse was excluded due to tendon displacement following tumour cell inoculation, resulting in n=9 in cancer group. No animals were excluded based on pre-defined exclusion-criteria related to climbing baseline performance.

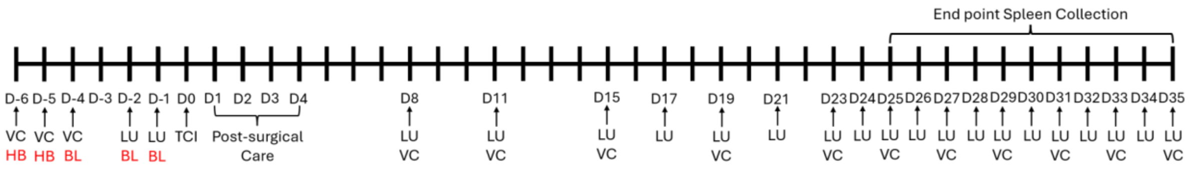

**Figure S6**. Experimental design and timeline for multiple myeloma bone pain model. *LU: Limb Use, WB: Weight Bearing, VC: Vertical Climbing, BL: Baseline, HB: Habituation, TCI: Tumour Cell Inoculation.*

Study limitations for CIBP studies:

In alignment with the 3Rs guideline to reduce animal use, an automated behavioural assay, Blackbox, was also tested in the same experiments which vertical climbing data reported in this study, although not reported here. Undisturbed Blackbox assessment was performed for 5 minutes recording immediately after limb use assessment and prior to vertical climbing. As this assay is observer free, non-biased, and spontaneous behaviour assay, the inclusion of this assay would not confound the vertical climbing results.

For breast cancer bone pain model, Blackbox assessments were performed on D7 and upon mice reaching limb use score 1 for morphine validation. For multiple myeloma bone pain model, Blackbox assessments were performed on D8, D11, D15, D17, D19, D21, and every day from D23 until reaching humane or experimental endpoint. Morphine validation was performed on D26 in this model, which may have influenced the vertical climbing on D27, and this timepoint was therefore removed. Nevertheless, the longitudinal trend of the disease remains consistent.

##### 1.1.14. Fibrous Dysplasia (FD) model

The transgenic mouse model of fibrous dysplasia (FD) used in this study was based on a previously reported *Prrx1-Cre*–driven, doxycycline-inducible *GαsR201C* transgenic system[14; 29]. Experimental mice carried the *Prrx1-Cre* and *ROSA26-rtTA* Linker alleles, with or without the *Tet-GαsR201C* transgene. Mice positive for *Prrx1-Cre*, Linker, and *GαsR201C* were classified as fibrous dysplasia mice, whereas littermate mice carrying *Prrx1-Cre* and Linker but lacking *GαsR201C* were used as controls. Genotyping was performed prior to group allocation.

Mice were housed in Digital Ventilated Cages (DVCs; Tecniplast) equipped with electromagnetics wheels (GYM500, Tecniplast) where mice locomotion activity and wheel running distance are automatically tracked [16]. For analysis, data generated during the dark phase were included.

In this study, doxycycline (doxycycline hyclate; Sigma-Aldrich) was administered in drinking water (protected from light) of 7 control and 5 FD male mice at approximately 13–14-week-old. All the mice were dosed with 0.1 g/L loading concentration for the first two days and changed to 0.08 g/L maintenance concentration until the end of the experiment. The study designs are shown in **Figure S7**. Vertical climbing was assessed at baseline, and D6 and 13 after initiation of treatment with doxycycline. X-rays were obtained under brief 2.5% isoflurane anaesthesia at baseline, Day 8, 15 and 22 after injury, for scoring disease burden. At the end of the experiment, mice were euthanised using cervical dislocation under isoflurane anaesthesia after the X ray. X ray, DVC cage monitoring and Doxycycline given were performed by a female experimenter (YT), while vertical climbing was performed by another female experimenter (AS). As all behavioural scoring was automated, the experimenters were not blinded.

Exclusions: No animals were excluded based on pre-defined exclusion criteria related to climbing baseline performance.

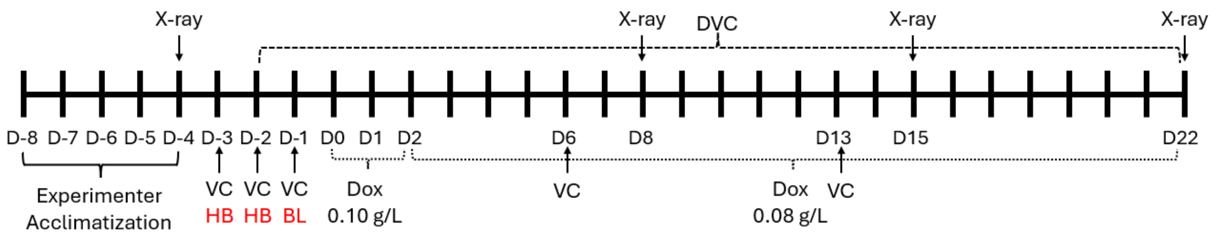

**Figure S7. Experimental design and timeline for FD bone pain model.** VC: Vertical Climbing, BL: Behavioural Baseline, HB: Behavioural Habituation, Dox: Doxycycline administration: loading dose: 0.10 g/L, maintenance: 0.08 g/L, DVC: Digital Ventilated Cages.

#### 1.1.15. Neuropathic pain model - Spared Nerve Injury (SNI)

To model neuropathic pain, the spared nerve injury (SNI) model was used [17]. In brief, mice were deeply anesthetized with isoflurane and injected s.c. with Temgesic (buprenorphine, 0.1 mg/kg, 0.03 mg/mL). A small incision was made in the skin of the left hind leg between hip and knee. A few drops of a cocktail of lidocaine (2.5 mg/mL) and bupivacaine (1.25 mg/mL) were applied topically to the muscle for 30 seconds before swabbed away. Using stump dissection, the muscles were separated to expose the sciatic nerve branches. A single surgical knot was tied around the tibial and peroneal nerve branches, leaving the sural nerve intact, before cutting out a small piece of the tibial and peroneal nerve branches. The surgical wound was closed with two medical clips and mice were injected with Rimadyl (carprofen, 5 mg/kg, 0.5 mg/mL) s.c. immediately after surgery. Rimadyl injections were repeated the two following days, and surgical clips were removed on day 3-4 post surgery.

Two different SNI-experiments were conducted, both including C57BL/6Nrj mice. One study included males only, and with the controls exposed to neither anaesthesia nor sham-injury. Another study was conducted with both males and females, and with a sham-injury consisting of anaesthesia, incision to the skin, medical clips and the same peri-operative care as described for the SNI-procedure. In the male-only, SNI and control-animals were not mixed in the same cage, while for the male-and-female study, two cages of females were mixed to obtain equal baseline vertical climbing behaviour between groups.

Animals were habituated to the experimental room for a minimum of 30 min before initiation of the experiment. The climbing test consisted of a training phase and a test phase, only the test phase was recorded. The training phase consisted of 2 days. On day 1 of training, the mice were placed in the grid as a group, meaning all mice from a single home cage were placed in the same grid and left to climb together for 15-20 min. On day 2 of training, the mice were placed in the grids individually and left to climb for 10 min to make sure all mice were able to climb freely in the grid. The test phase consisted of one baseline test before surgery and 3-5 tests after surgery (male-only study: Day 3+7+12 after SNI surgery, male-female-study: Day 4 + 9 + 16 and a cross-over drug-test Day 27+30). At each test day, mice were placed in the grids individually and left to climb for 12 minutes. A video camera placed in front of the grids recorded the animals' behaviour on test days and Ethovision XT 15 (Noldus) was used to track the distance travelled by the animal. Two grids were used at the same time placed in a manner that any mouse in one grid could not see the mouse in the other grid, only hear them.

GBP study: For the male-female study, a cross-over study exploring the effects of gabapentin (30 mg/kg, i.p.) vs vehicle (saline) was included on Day 27 and 30 after injury, including equal numbers of animals from each experimental treatment on each day, and 3-day washout between. The effects on climbing were assessed 1h after injection.

Behavioural tests for the male-only study were performed by the same female experimenter (KLJ), while another female experimenter performed the male-female study (IBK). As all behavioural scoring was automated, the experimenter was not blinded.

Exclusions: for the SNI-characterisation and later GBP-study, 4 animals were excluded based on the pre-defined exclusion-criteria on baseline climbing activity, 3 females and 1 male. The day of surgery, a sham-injured female was found dead in the cage, for unknown reasons.

At the end of the experiment, mice were euthanised using cervical dislocation.

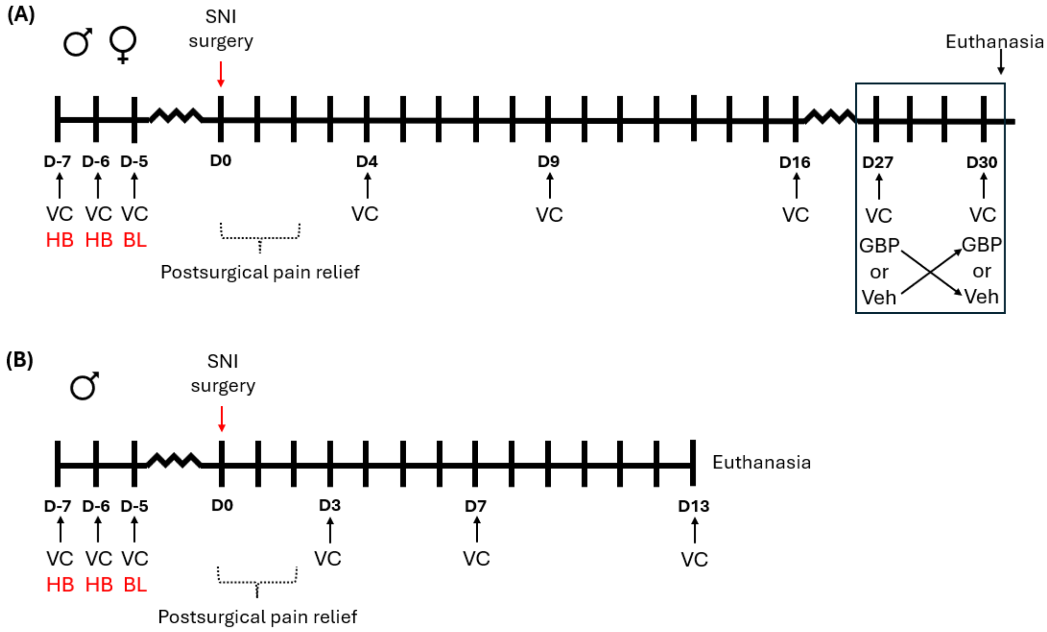

**Figure S8. Experimental design and timeline for SNI experiments. A)** Study including both males and females, exploring the effects of SNI vs sham-injured animals. **B)** Male-only study, exploring the effects of SNI vs naïve controls. VC: Vertical Climbing, BL: Behavioural Baseline, HB: Behavioural Habituation, SNI: Spared Nerve Injury. GBP: gabapentin, 30 mg/kg, Veh: Vehicle.

#### 1.1.16. Mouse model of levcromakalim-induced migraine

The mouse model of levcromakalim-induced migraine was established through injection of levcromakalim (1 mg/kg, i.p) every other day, days 0, 2, and 4 [3]. Levcromakalim (Tocris Bioscience, Bristol, UK, batch no: 1A/275955) was dissolved in saline with 2% DMSO to a final concentration of 0.01 mg/ml, which was injected i.p. in the lower right abdominal quadrant in a volume of 10 ml/kg using a 27-gauge needle. Levcromakalim and the corresponding vehicle solution were stored at -20 degrees Celsius and not subject to any freeze thaw cycles prior to use.

The vertical climbing test was performed 1 and 2 hours post injection of levcromakalim on test days 0 and 4. On test day 2, mouse grimace scoring (MGS) was performed 1 h after levcromakalim injection. Vaginal lavage samples were taken from female mice after behavioural testing on all test days for oestrous cycle stage determination.

Mice were allocated to vehicle or levcromakalim groups with stratification to baseline climbing, sex, and home cage.

All behavioural experiments were performed by the same female experimenter (JJJ). As all behavioural scoring was automated, the experimenter was not blinded.

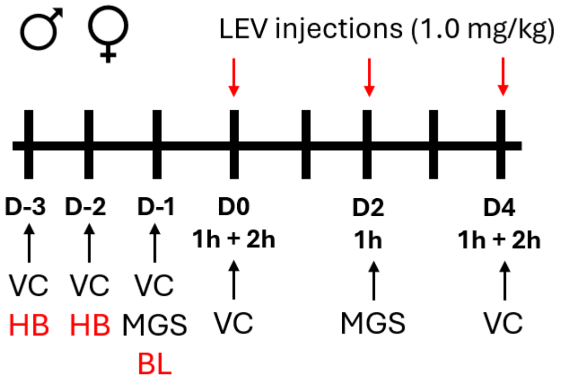

**Figure S9. Experimental design and timeline for levcromakalim experiments.** The study including both males and females, exploring the effects of LEV vs VEH. LEV: levcromakalim, VEH: vehicle, VC: Vertical Climbing, BL: Behavioural Baseline, HB: Behavioural Habituation, MGS: Mouse Grimace Scale.

At the end of the experiment, mice were reused for other experiments.

#### 1.1.17. Periorbital neuropathic pain model (FRICT-ION)

FRICT-ION surgery was performed on male and female C57BL/6J mice as described[20]. Mice were anesthetized with 5% isoflurane and maintained on 2.5% isoflurane throughout the procedure. A small incision was made in the inner buccal mucosa to expose the maxillary branch of the right infraorbital nerve near the foramen rontundum. A 3 mm segment of 4-0 chromic gut suture (Ethicon #724G) was inserted along the nerve into the foramen rotundum. Mice recovered on a heating pad until fully ambulatory and were returned to their home cage. Animals were monitored daily for any changes in body weight, healing, and infection. No animals were excluded due to complications from surgery.

The vertical climbing test was performed at baseline before surgery. Animals were also tested for facial mechanical (von Frey) and cold (acetone droplet assay) hypersensitivity prior to surgery. Mice were divided into groups (sham or FRICT-ION) based on sex and baseline climbing behaviour. Following surgery, mice underwent mechanical and cold facial sensitivity testing and the vertical climbing test every seven days for four weeks. Evoked testing and the vertical climbing test were performed on separate days. The experimenter was blinded to surgical condition (sham or FRICT-ION), and all quantification was performed blinded.

Exclusions: A total of 5 mice (1 female, 4 male) were excluded based on the pre-defined exclusion-criteria related to climbing baseline performance.

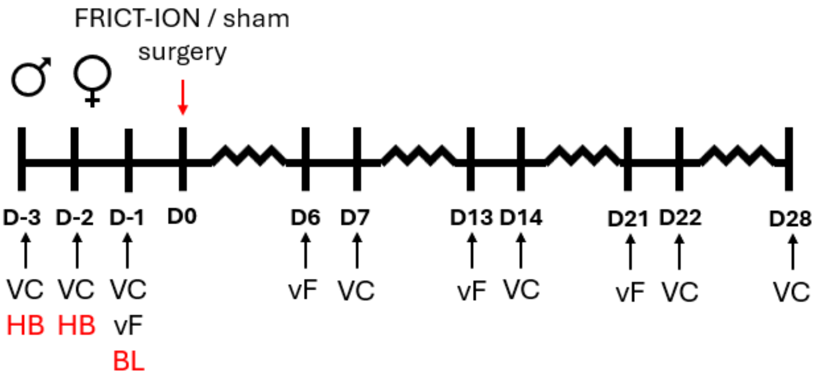

**Fig S10: Experimental design and timeline for FRICT-ION experiments.** The study including both males and females, exploring the effects of FRICT-ION vs sham-injured animals. VC: Vertical Climbing, BL: Behavioral Baseline, HB: Behavioral Habituation, vF: von Frey (mechanical allodynia).

### 1.2. Measures / assays

#### 1.2.1. Vertical climbing

Our vertical climbing setup was based on the previously published device from Santos et al. [25], although with modifications. Initial pilot experiments explored different dimensions and wire mesh material. We considered fully encasing the experimental setup to provide surroundings with less disturbance from sounds, smells, and other environmental factors, but ultimately decided against it. The total time each mouse spent climbing in the different prototype assays was assessed, and the results showed that an assay with a diameter of 15 cm and height of 25 cm, without a lid, was where mice exhibited the highest climbing activity. This assay configuration was therefore selected for use in the final experimental setup, and a custom 3D-printed platform was developed to secure stability. This setup was then used for all experiments in this study. The aim was an easy-to-build assay, to be made freely available for anyone who would wish to employ it in their experiments. See the full construction-manual and Standard Operating Procedure (SOP) on the open protocol repository, protocols.io: https://dx.doi.org/10.17504/protocols.io.n92ldo7q7g5b/v1 .

The vertical climbing assay consisted of an enclosed vertical cylinder made of galvanized steel mesh (height: 250 mm, diameter: 150 mm, mesh size: 6 mm x 6 mm), standing on a 3D-printed platform made of either black or grey Polyethylene Terephthalate Glycol (PETG). For testing, animals were carefully placed on the inside of the mesh cylinder, and the mesh was carefully placed vertically in the stand. A camera was placed in front of the assay from approximately mid-height of the cylinder to capture the full cylinder, including base, to record the animal’s behaviour in the assay (**see Fig 1**).

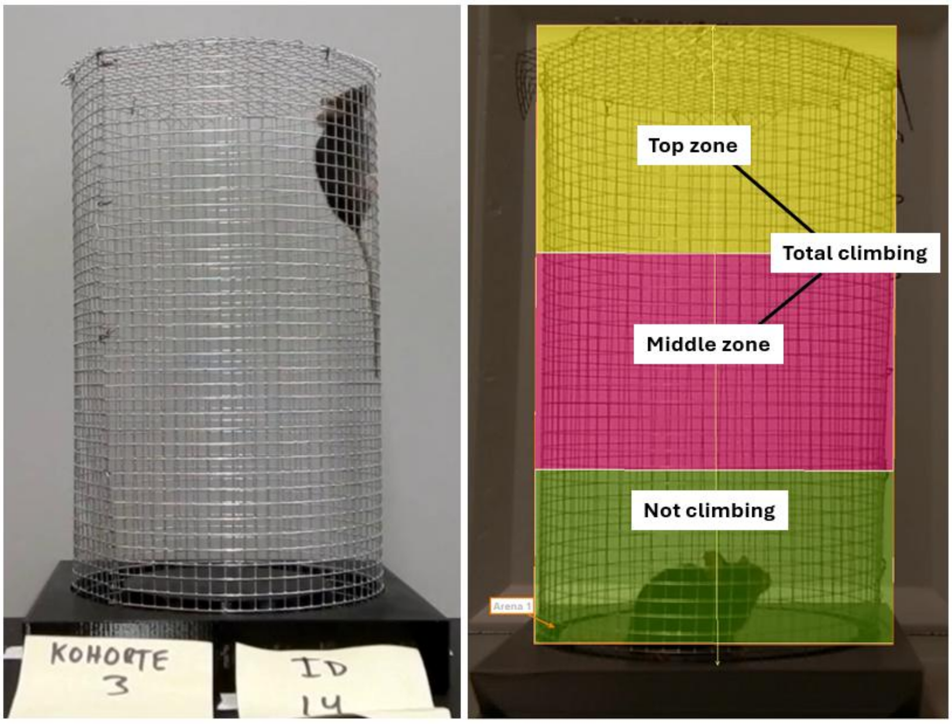

**Fig 1:** Example of a C57BL/6 mouse climbing the vertical climbing cylinder. Right picture displays the Ethovision arena-settings used for tracking.

The protocol for the climbing test consisted of a 2-day habituation / training phase, followed by a baseline test before model-induction.

- Day 1 - group habituation: mice were placed in the cylinder as a group, meaning all mice from a single home-cage were placed in the same grid and left to climb together for 15-30 min. This provided minimally stressful conditions for the mice to explore the cylinder with their cage-mates.
- Day 2 - single habituation: the mice were placed in the cylinders individually and left to climb for 10-15 min to make sure all mice were able to climb freely on the grid.
- Day 3 - baseline test: each mouse was placed individually in the cylinder, and behaviour was recorded for 11-12 min.. For analysis, the first minute was always excluded, to allow the experimenter to move away, and the animal to settle down. After the first minute, the following 10 min. were used for analysis and tracked either live or retrospectively using Ethovision XT (Noldus technology, NL)

- Test-days after injury: the timing of the following test-days depended on the injury-model and experimental hypothesis, but the protocol followed the same steps as the baseline test above.

Between each animal, the platform was cleaned with 70% ethanol, to reduce the risk of odour cues affecting the behaviour. Often, two vertical climbing cylinders were used to allow two animals to be tested simultaneously, and under such circumstances, great care was made to place the setups as far away from each other as possible, and in a manner that any mice in one cylinder could not see the mouse in the other cylinder, using dividers or similar.

Video recordings were analysed using Ethovision XT17, 16, or 15 software for detection of the centre point of the animal when moving around in the cylinder. After discarding the first minute, ten-minutes of videorecording was analysed, starting from minute 1 to minute 11 using a dedicated arena-template including a lower "non-climbing"-section, "middle climbing"- and "top-climbing" sections, as shown in **Fig 1**. As each lab used their own cameras and behaviour rooms, the tracking-templated will have led to slight differences between each lab, depending on the angles and as more or less of the floor or ceiling could be seen in a given video frame. This may also have given some inaccuracies in detecting the 'distance moved' when climbing in the front or backside of the metal cylinder. Additionally, for each experimental setup, it was checked that when a given animal was standing on the hindlegs, that the centre point did not move into the climbing-zone. Importantly though, within each experiment, tracking- and arena-settings were kept consistent for each trial, making comparisons between groups not affected by these small inaccuracies. Vertical climbing duration (in top + middle zone), frequency in top, and distance moved in the full cylinder, were retrieved and further processed for analysis.

To ascertain appropriate levels of natural climbing behaviour, exclusion-criteria were implemented. The exclusion-criteria were applied to the "Total climbing" (top + middle zone) duration at baseline, and a given mouse was excluded from the climbing study, if it spent either 1) more than 570 seconds, or 2) less than 30 seconds climbing, out of the full 600 seconds tested (10 min). This constitutes 95% of the time tested, where the mouse was either climbing or non-climbing, respectively. The rationale being, that the assay is intended to quantify voluntary climbing activity, and potential conditions which could modify this activity, and this goal would be undermined if, 1) the animal did not consider the base as a safe place to relax / avoid climbing, or 2) if the animal had no natural desire to climb under normal circumstances. Under such circumstances mice were considered non-compliant and were excluded. When possible, baseline analysis was made before animals were allocated into groups, for exclusion of extreme- and non-climbers.

Depending on the experiment, different camera-types were used (Nikon Coolpix AW300, JVC Everio GZ-MG645, Logitech webcams: "BRIO 105", "C920 Pro", Infrared: Basler, acA1300-60gm), and the Ethovision-tracking was possible for all cameras, despite the animal moving behind the wire mesh. But be aware that for automated scoring, the contrast between the animal's fur-colour and the background wall and base-stand can be key for optimal detection. For one infrared camera, this proved particularly difficult, especially when the animal was on the base stand, despite using a grey stand for black mice. Therefore, the 'distance moved' parameter was deemed compromised for this dataset (migraine model), as the animal was not detected properly while on the base, but when moving around in the cylinder, the tracking worked optimally, making the frequency and duration climbing parameters intact. Later, the issue was resolved by using white platforms for infrared tracking of black mice.

##### 1.2.2. Mechanical allodynia, von Frey (intraplantar CFA study)

Animals were habituated to the testing room for 1 hr prior to each session, then placed in red tinted PVC cups on a plastic grid (Bioseb, BIO-STD2_EVF) for 10-15 minutes before assessment of mechanical sensitivity through the "ascending stimulus" method [6; 26], where animals are tested from the lowest to highest filament eliciting a positive response in ascending order. 3 out of 5 withdrawal reactions (paw withdrawal or paw lick) to the same filament were recorded as a positive response, with all 5 applications performed within approximately 30 seconds, with a minimum 5 second pause between applications of the same filament. Filaments included ranged from 0.07 to 2 g (g = gram force) (0.07, 0.16, 0.4, 0.6, 1.0, 1.4, 2.0) (Bioseb, BIO-VF-M). Both ipsi and contralateral paws were tested as internal control.

##### 1.2.3. Facial allodynia, von Frey and acetone droplet test (FRICT-ION study)

Animals were habituated to light restraint (Plexiglas tubes, with their tails fixed in place) for three days before behavioural testing was performed. On test day, mice were placed in light restraint tubes for at least one hour before testing began. The same female experimenter performed all time points (NKG). Von Frey filaments were used to assess facial mechanical hypersensitivity by probing the face directly behind the whisker pad using the up/down method. Facial cold allodynia was assessed using the acetone droplet test. A 3 mL syringe attached to PE10 tubing with a flared end was used to place a 10 µL droplet of acetone on the animal's face directly posterior to the whisker pad. The time spent reacting (grooming, face wiping, etc) was recorded in seconds. Two trials were performed per animal, and the average of both was reported.

##### 1.2.4. Dynamic Weight Bearing (MIA study)

Dynamic Weight Bearing (DWB) was assessed in the MIA-model, using the Dynamic Weight Bearing 2.0 apparatus (Bioseb, Vitrolles, France), as before [2]. The mice were tested one sex at a time to minimize scent of the opposite sex. Mice were individually placed in an 11 x 11 x 20 cm chamber on a sensor pad detecting pressure, and a camera in the lid detecting posture. Mice were allowed to first habituate to the device for 5 min, before capturing pressure data and live recording for 5 min of free movement within the chamber (sensor threshold: 30, background: 65, immobility: 700ms). The equipment was cleaned with 70% ethanol between each subject.

Manual validations of the acquisitions were next assessed blindly for each acquired trial. The manual validation consisted of verifying the placement of the animal's limbs by comparing live recordings with a scaled map of the activated pressure sensors of the floor sensor pad. Sequences where the experimenter was uncertain about the correct pairing of limbs and pressure readouts from other body parts, such as the tail, were excluded. A minimum of 1 minute of validated testing was used to calculate weight bearing characteristics for each mouse [23; 24]. Of the many values computed by the program following validation, only the DWB values of the hindlimbs were used. DWB values were obtained as the percentage of body weight (%BW) distributed on the ipsilateral leg during immobility.

As one recording did not meet the preset criteria of 1minute of useful, validated recording on day 15/16, the data for that animal was not included.

##### 1.2.5. Arthritis score (CAIA study)

For the CAIA studies, clinical signs of active joint inflammation was assessed daily using the previously published methods by Nandakumar et al. [21] In short, each animal was assessed for visual signs of arthritis - redness and swelling - by clinical paw scoring. Each animal was gently restrained, and inflammation was assessed on all limbs, giving each inflamed toe or knuckle 1 point, while an inflamed wrist or ankle was awarded 5 points (see **Fig S11**). Thereby, each limb could receive a score from 0 to 15 (five toes + five knuckles + one wrist/ankle), while 0 to 60 points for each mouse. Thereby, the higher the score, the more widespread signs of disease activity.

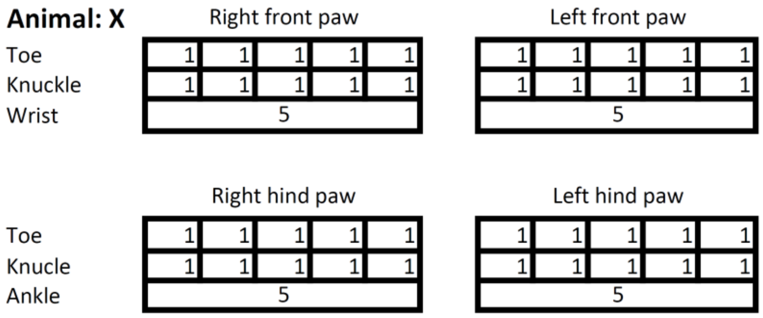

**Fig S11**. Arthritis scoring scheme. Constructed based on description from Nandakumar et al. [21].

#### 1.2.6. Burrowing (CAIA study)

Burrowing behaviour was assessed as an ethologically relevant, species-typical behaviour reflecting spontaneous behavioural function [27; 28]. The test is based on previously described burrowing paradigms with minor modifications [27]. The burrowing test was performed in a standard transparent plastic cage (Macrolone Type 3) without bedding, with the burrowing tunnel including burrowing material placed inside. Burrowing tunnels were self-build as previously[27] and were made from grey polyvinyl chloride (PVC) tubes (diameter: 75mm, length: 210mm) which were closed in one end and raised by metal screws in the other end (see **Fig S12**). For burrowing-substrate, we used the standard saw dust also used for bedding. Each animal had a dedicated burrowing tunnel + cage, but when switching between animals, they were washed with warm water and soap, then disinfected using Virkon S followed by 70% ethanol and allow to air-dry.

Habituation was performed over three consecutive days from day -5 to day -3, prior to CAIA induction on day 0. During the first two days, mice were habituated in pairs, whereas on the third day mice were performed individually in the testing cage. On the first habituation day, mice were exposed to the burrowing tunnel without bedding material for 1 h. On the second and third days, the tunnel contained 100g of bedding material (wood shavings, Brogaarden), and mice were allowed to interact with the apparatus and burrowing material for 2 h. At the end of each habituation period the mice were returned to their home cages.

Experimental sessions were conducted over four days; two baseline assessments on day -2 to day -1 prior to CAIA induction, followed by assessments on day 6 and 13. During each experimental session, mice were tested individually in their assigned cages using the same burrowing tunnel throughout the study. Each tunnel was filled with 100 g of bedding material (wood shavings, Brogaarden), and mice were allowed to interact with the apparatus for 120 min. Following the test period, the bedding material remaining inside the tunnel was weighed, and the amount displaced was used as the measure of burrowing activity.

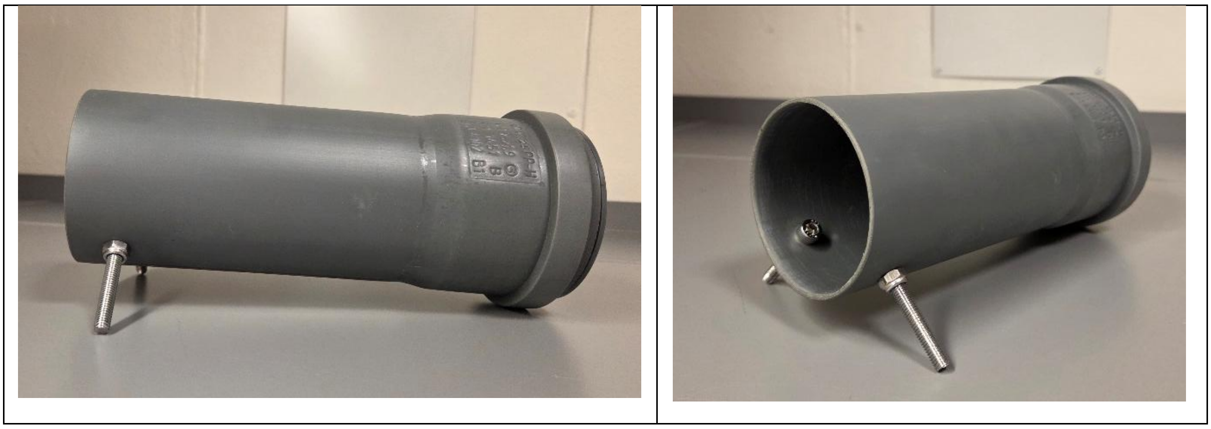

**Fig S12:** Burrowing tunnels.

#### 1.2.7. Limb use (Bone Pain studies)

During each assessment session, mice were first habituated with their cage mates in a standard transparent cage for 125 x 266 x 185 mm (Techniplast, Italy) for ~10 minutes. Afterwards, individual mice were placed in a similar cage and video-recorded (Logitech C920 Pro Webcam) for 3 mins. A masked observer then evaluated their gait use and scored for limb use score. For breast cancer bone pain model, mice were scored as follows: 4 – Normal gait, 3 – Insignificant limping, 2 – significant limping and shift in bodyweight towards contralateral limb, 1- significant limping and partial lack of use of the ipsilateral limb, 0 – total lack of use of the ipsilateral limb [15]. For multiple myeloma mice, mice were scored as follows:4 – normal gait, 3 – insignificant limping, 2- significant limping and shift in bodyweight distribution towards contralateral limb, 1 – significant limping and motor impairment, 0 – paraplegia[7].

#### 1.2.8. Static Weight bearing (Bone pain studies)

Incapacitance Tester Version 5.2 (Linton Instrumentation, UK) was used to assess the static weight bearing on the hind legs. Before baseline assessment, mice were habituated and trained to position themselves on rearing position for a minimum of 3 seconds with individual hind paws placed on top of independent weight-sensing plates. For each assessment session, weight bearing was recorded over period of 3 seconds in triplicates for each mouse. The distribution of weight on the hind paws was determined with the following equation: % weight bearing = weight on ipsilateral limb/weight on ipsilateral limb + weight on contralateral limb (total weight on hind limbs).

#### 1.2.9. Home cage activity and wheel running (FD study)

Mice were single housed in Digital Ventilated Cages (DVCs; Tecniplast) equipped with electromagnetics wheels (GYM500, Tecniplast) where mice locomotion activity and wheel running distance are automatically tracked [16]. The DVC rack is equipped with an electromagnetic sensor plate employing capacitive sensing technology to monitor the locomotion activity of the mouse within the cage. This sensor plate contains 12 electrodes that emit electromagnetic fields. When the mouse passes through different fields, a disturbance is generated, which means the data is recorded and measured. Once the cage is placed on the rack properly, continuous and undisturbed readings are available every hour, which enables sustained monitoring during the 12-hour dark period when the mice are most active, without experimental intervention. Furthermore, the DVC running wheel where two magnets are placed was equipped inside the cages and automatically monitor the rotation, speed, and distance by matching the sensor on the DNC rack. For analysis, data generated during the dark phase were included.

#### 1.2.10. X ray imaging and bone deformity scoring (FD study)

X ray imaging was conducted weekly using Lumina XR IVIS (In Vivo Imaging System) apparatus (Caliper Life Sciences, Teralfene). Mice were sedated and maintained using 2.5% isoflurane (1000 mg/g isoflurane, Attane vet, ScanVet) for 5-10 mins. X ray images were obtained of the entire skeleton.

The x-ray images were analysed to determine FD burden of each individual animal at different time points. The primary focus was to follow the progression and to investigate a possible correlation between disease burden and performance in the climbing assay. An ordinal scale was defined in advance with 0 - 3 points depending on the lesion severity in each area of interest (**Table S2**).

The scoring was performed by a manual, independent evaluation of each segment involved in the scoring system. The areas of interest were the calcaneus, tibia, fibula, femur, radius and ulna. The tibia was divided into distal and proximal parts (**Figure S13**). The right and left legs were scored individually and subsequently added together to achieve a total score for each mouse. The scorings were performed by one scorer, and the average score was calculated and used for analysis. The images were blinded during scoring.

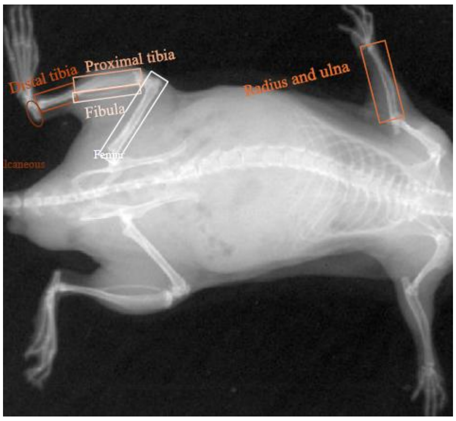

**Figure S13. Areas of interest of scoring on a reference mouse.**

| Bone | Score | Description |
|---|---|---|
| Calcaneus | 0 | Intact |
| | 1 | Mild radiolucent change |
| | 2 | Cortical thinning and/or bowing |
| | 3 | |
| Distal Tibia | 0 | Intact |
| | 1 | Mild radiolucent change |
| | 2 | Endocortical irregularity and cortical thinning |
| | 3 | Marked endocortical loss, expansion, bowing, cortical fracture |
| Proximal Tibia | 0 | Intact |

| | | |
|---|---|---|
| | 1 | Radiolucent/ground-glass lesion with endocortical loss |
| Fibula | 0 | Intact |
| | 1 | Mild radiolucent change |
| Femur | 0 | Intact |
| | 1 | Endocortical irregularity or cortical thinning |
| | 2 | Heterogeneous marrow density, expansion, bowing, cortical fracture |
| Radius/Ulna | 0 | Intact |
| | 1 | Mild radiolucent change |
| | 2 | Bowing, expansion, endocortical irregularity, cortical thinning, or cortical loss |

**Table S2. FD mouse bone scoring template**

1.2.11. Mouse grimace scoring (mouse model of levcromakalim-induced migraine):

For grimace scoring, mice were placed in a white cardboard expresso cup held in a horizontal position at the edge of a table using a custom-made device (**Fig S14**). Mice were tunnel handled from home cage to the cup placed in front of a video camera (LEGRIA HF G70, Canon Inc., Japan). During recordings, the experimenter was silently present in a corner of the room. To ensure sufficient light to separate facial features in the black coated mice, recordings were made with 1900-2000 lux illumination. The light source was in the ceiling, and an additional light source was placed above the set up (Luxo T88/87, 230V, 50Hz).

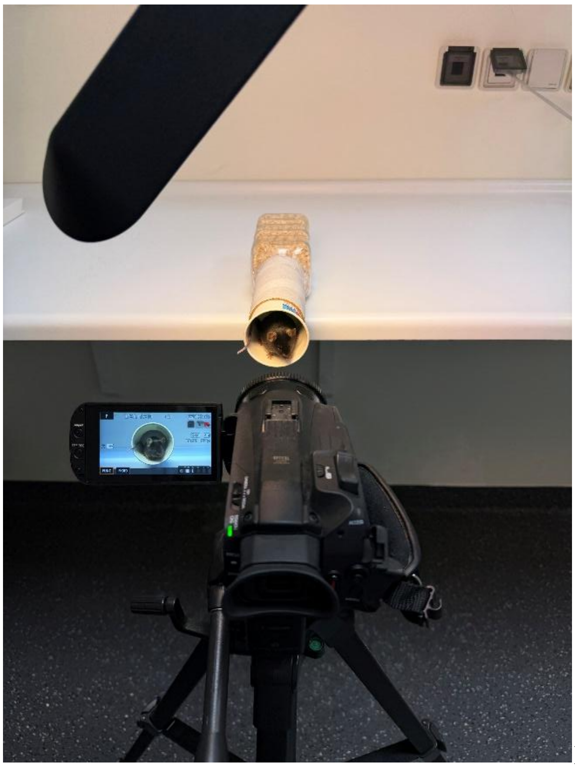

**Fig S14.** Setup for MGS scoring.

Mice were habituated to the espresso cups a single time for 30 minutes a few days before baseline measurement. Furthermore, mice were acclimatized to the set-up for 15 min. immediately before recoding. The actual recording for grimace scoring was 11 minutes. Videos were scored by use of PainFace an open-source machine learning tool developed specifically for mouse grimace scoring of black coated mice (http:/painface.net [19]). Fau model ID: default (20250826-generel), pain-mgs model ID: default (2022115-black), Sample Rate: High (1 frame/second). For each frame, PainFace detects four facial features; orbits, nose, ears and whisker and each is assigned a value from 0 to 2 [18]proving mean total grimace scores between 0 and 8. The first 10 frames of the video were skipped to make sure correct tracking of the mouse and the next 600 frames were analysed (10 min). Only frames with all four facial features detected were included, and measurements of each facial feature were required to have a confidence score of at least 0.8. Recordings with less than 75 valid frames were discarded as recommended by the developers. In our study, these settings resulted in an average of 205 frames scored pr. recording which were averaged to provide the mean total grimace score.

##### 1.2.12. Determination of oestrus cycle (mouse model of levcromakalim-induced migraine):

Female mice were subject to vaginal lavage after behavioural testing on baseline and experimental Days 0 and 4 following procedures slightly adapted from Cora et al. [4]. To prevent contamination of the sample, a paper towel was used to wipe off any noticeable debris/urine on the vulva. A 100 µL pipette tip was filled with 50 µL saline and the tip was inserted a maximum of 2 mm into the vaginal canal. The canal was then flushed 2-3 times with approximately 25 µL of saline. A drop of the collected vaginal fluid was placed on a glass slide. The fluid was smeared in an even thin layer with the pipette end, then dried and subsequently stained by a 1 min dip in 0.1% toluidine blue. Last, the slide was rinsed in distilled water for one minute to wash away excess dye. The dried sample was evaluated on a light microscope (Nikon Eclipse Ni) using the 10x and 20x lenses and categorised into one of the four stages of the mouse oestrous cycle: prooestrus, oestrus dioestrus, metoestrus depending on the presence of cell morphologies and cellularity [4] (**Fig S15**). Two individual assessors performed independent scoring of the 78 samples (JJJ & SLC). For 33 samples, there were discrepancies between the verdicts, and these were then scored by a third independent assessor (AH). The final verdict was the score determined by two of the three assessors. One sample was inconclusive and excluded due to limited number of cells.

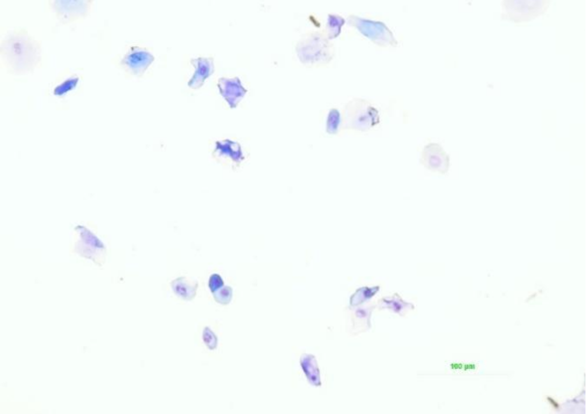

**A. Prooestrus**, dominated by nucleated epithelial cells, seen as round dense cells with a basophilic nucleus. Few keratinized cells can be spotted, often with no leukocytes.

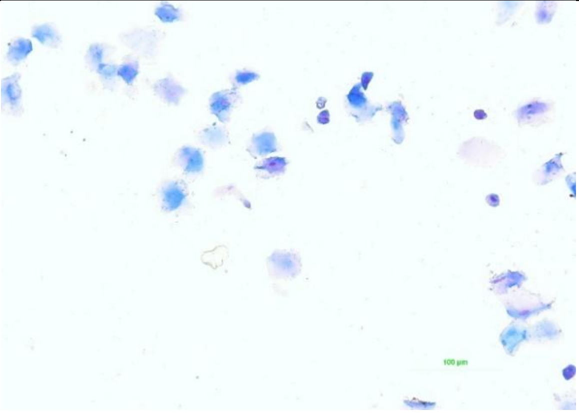

**B. Oestrus** stage dominated by anucleated keratinized epithelial cells, seen as large cells, with ridged edges and absent nuclei. Bacteria are visible in the background.

| | |
|---|---|
| 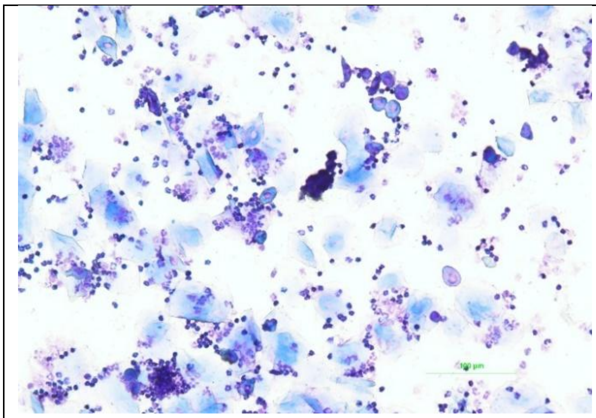 | 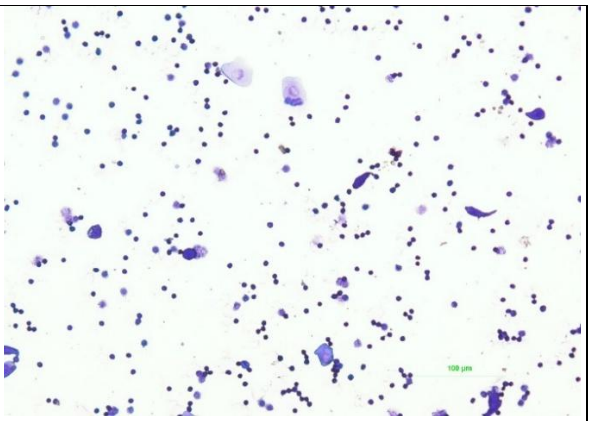 |
| **C. Metoestrus,** with many anucleated keratinized epithelial cells and leukocytes | **D. Dioestrus**, with both leukocytes and nucleated epithelial cells, often observed with fewer anucleated keratinized epithelial cells |

**Fig S15. Oestrus classification.**

### Supplementary data

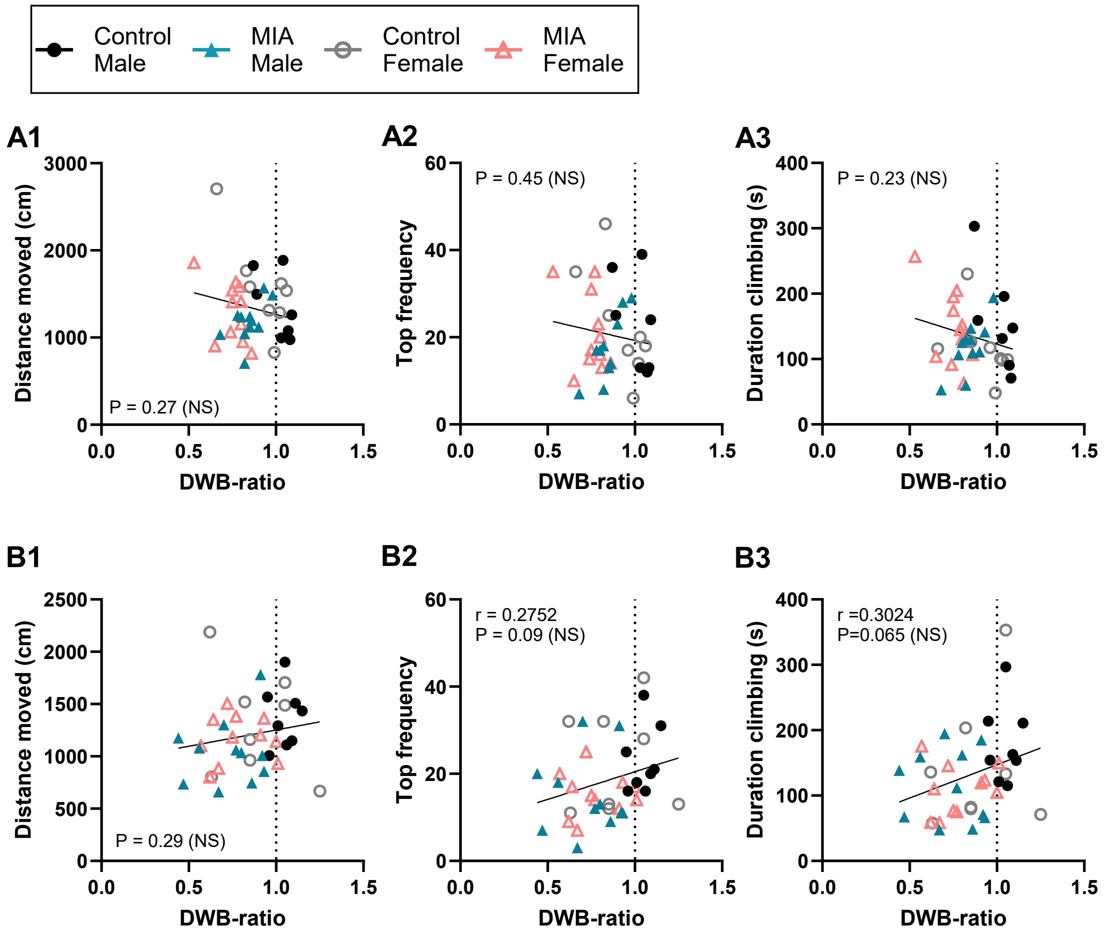

**Supplementary Figure S15**, MIA. **A**: Correlations between vertical climbing and DWB at two weeks. **A1**) Distance moved, **A2**) Top frequency, **A3**) Duration climbing. **B**: Correlations between vertical climbing and DWB at four weeks. **B1**) Distance moved, **B2**) Top frequency, **B3**) Duration climbing. Analysis conducted via Pearson's correlation analysis.

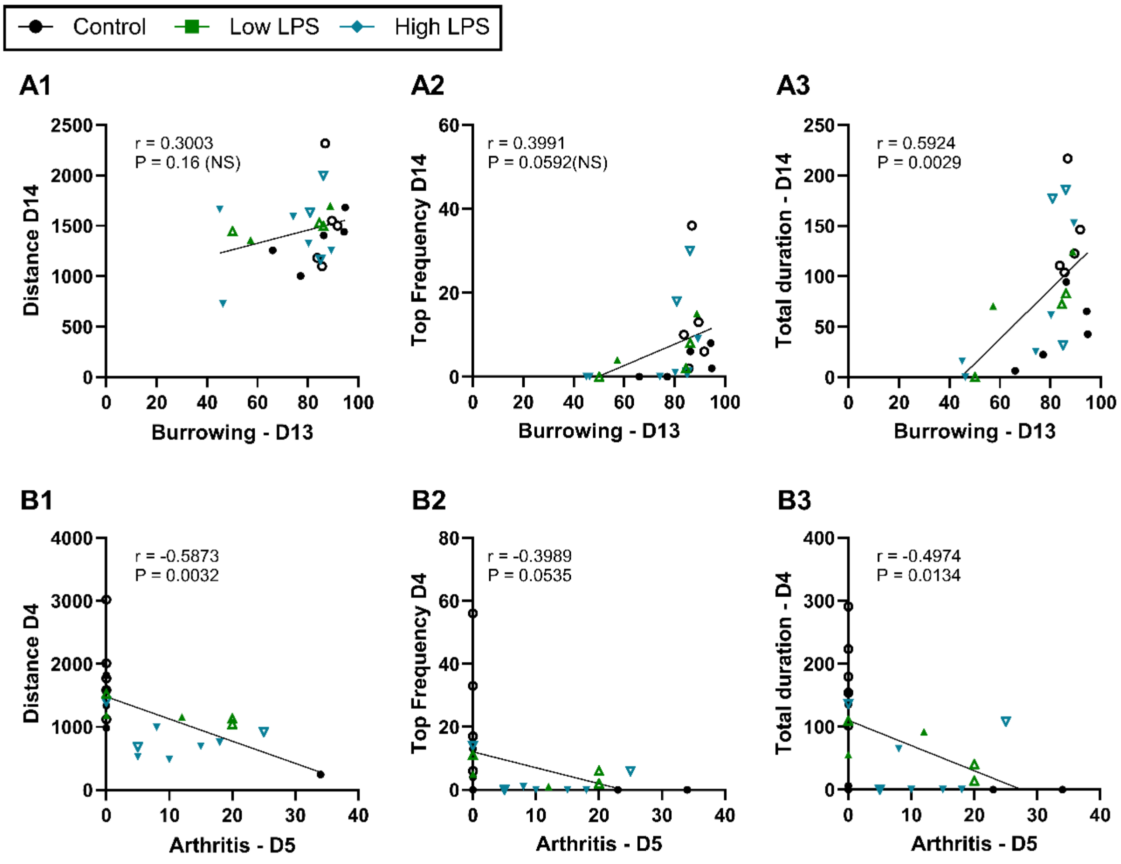

**Supplementary Figure S16.** CAIA supplementary data. **A:** Correlations between burrowing D13 and vertical climbing D14. **A1**) Distance moved, **A2**) Top frequency, **A3**) Duration climbing. **B:** Correlations between Arthritis score D5 and vertical climbing outcomes on D4. **B1**) Distance moved, **B2**) Top frequency, **B3**) Duration climbing. Analysis conducted via Pearson's correlation analysis.

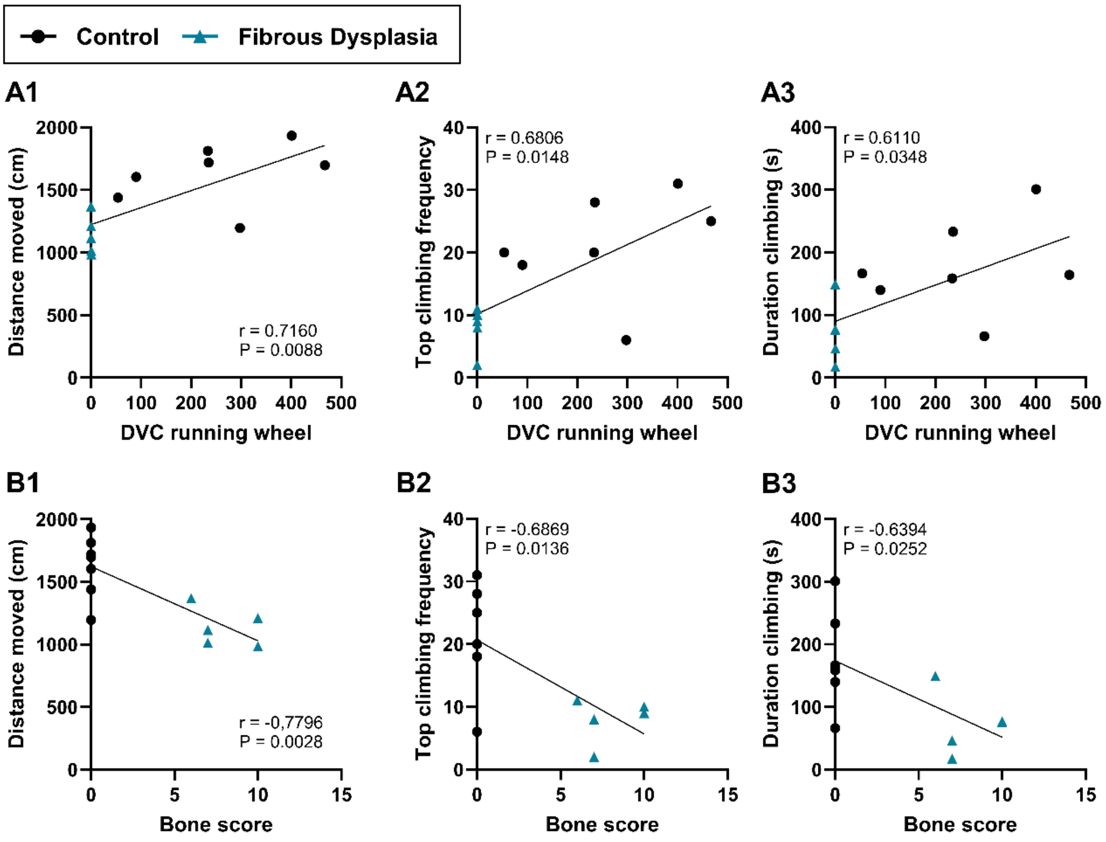

**Supplementary Figure S17.** Fibrous Dysplasia supplementary data. Correlations between climbing and **A**: running wheel: **A1**) Distance moved, **A2**) Top frequency, **A3**) Duration climbing, and **B**) bone score: **B1**) Distance moved, **B2**) Top frequency, **B3**) Duration climbing. Analysis conducted via Pearson's correlation analysis.

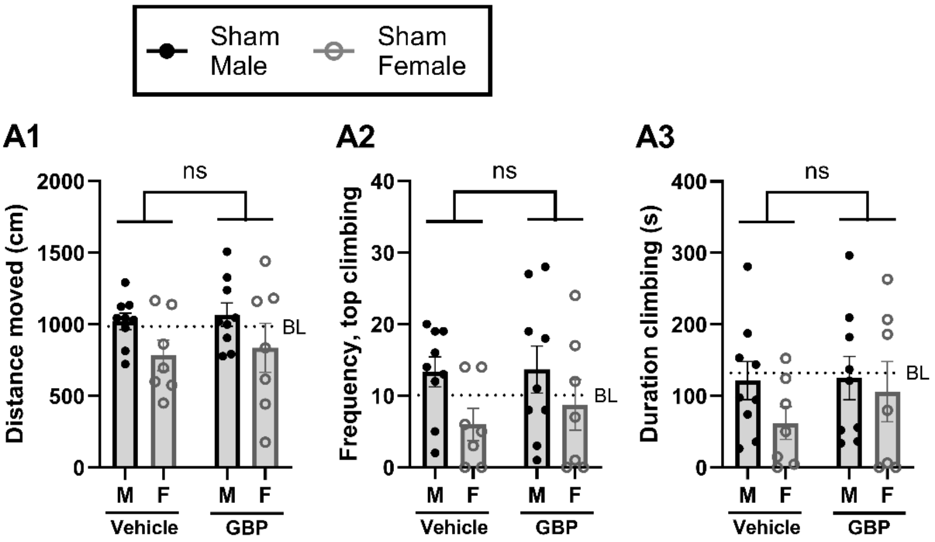

**Supplementary Figure S18.** Sham GBP. (**A**) On Day 27/30 a cross-over study testing i.p. gabapentin (GBP, 30 mg/kg) vs. vehicle was conducted, showing no effects of GBP on climbing behaviour in sham control animals, as measures through (**A1**) distance moved, (**A2**) duration climbing, and (**A3**) frequency in the top. Dotted line suggests pre-injury baseline average. *: P<0.05, **: P<0.01, NS: not significant, as suggested by injury-effect in a 2-way ANOVA (factors: sex & injury). F = female, M = male.

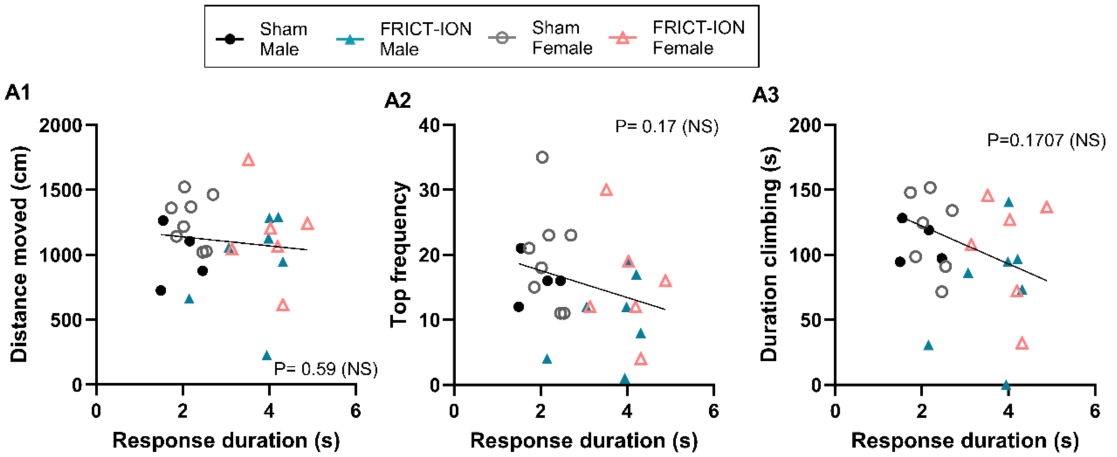

**Supplementary Fig S19**. FRICT-ION model, additional correlations with cold allodynia on D14. None are significant. **A1**) Distance moved, **A2**) Top frequency, **A3**) Duration climbing. Analysis conducted via Pearson's correlation analysis.

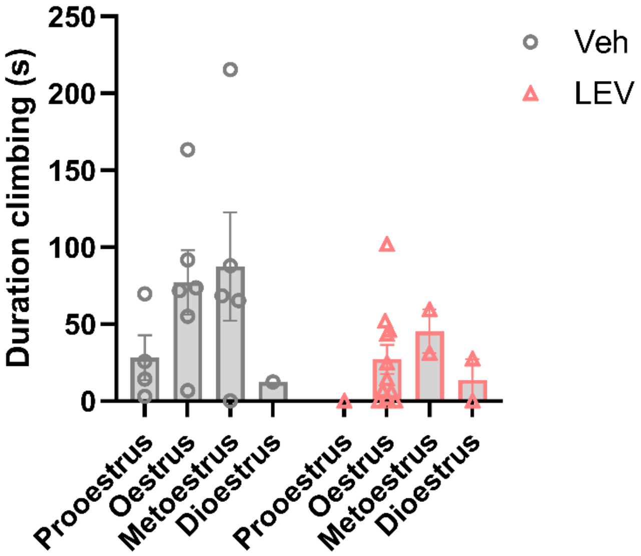

**Supplementary Fig S19.** Oestrus cycle in the LEV-induced migraine model. Day 0 + 4, 1h after injection combined. 2-way ANOVA clarified: $F_{cycle}$ (3, 24) = 1.420, P=0.2614 (NS).

### Supplementary Table S3 – Results of statistical analysis

| Fig | Analysis | F-values | Post test |
|---|---|---|---|
| **Fig 1. Incision model** (Male: Ctrl: n=11, Incision: n=12, Female: Ctrl: n=12, Incision: n=10) Meloxicam study (males only): incision-vehicle: n=9, incision-meloxicam: n=12) Missing video's/data-points (random): a full female cohort at 72h (12animals), 2 females: 48h, 1female 72h. Therefore, mixed-effects models were applied to account for missing values. | | | |
| A1. Distance Timecourse | Mixed-effects model, sex*injury*time, covariate: baseline (SPSS, mixed model analysis, linear) | $F_{sex}$ (1,50.6) = 2.631, P=0.111 (NS)<br>$F_{injury}$ (1,49.6) = 3.726, P=0.059 (NS)<br>$F_{time}$ (4,142.9) = 12.059, P<0.001<br>$F_{Covariate, BL}$ (1,50.7) = 4.920, P=0.031<br>$F_{sex*injury}$ (1,51.4)= 0.195, P=0.66 (NS)<br>$F_{time*injury}$ (4,140.3) = 1.156, P=0.333 (NS) | |
| A2. Distance – 4h | 2-way ANOVA, sex*injury | $F_{sex}$ (1,41) = 2.931, P=0.094 (NS)<br>$F_{injury}$ (1,41) = 6.499, P=0.015<br>$F_{injury*sex}$ (1,41) = 1.295, P=0.26 (NS) | (injury effect presented on fig) |
| B1. Frequency in top, timecourse | Mixed-effects model, sex*injury*time, covariate: baseline (SPSS, mixed model analysis, linear) | $F_{sex}$ (1,49.6) = 1.441, P=0.236 (NS)<br>$F_{injury}$ (1, 48.5) = 6.599, P=0.013<br>$F_{time}$ (4, 141.2) = 10.202, P<0.001<br>$F_{Covariate, BL}$ (1,49.7) = 3.856, P=0.055 (NS)<br>$F_{sex*injury}$ (1,50.8)= 0.046, P=0.83 (NS)<br>$F_{time*injury}$ (4,138.2) = 1.281, P=0.281 (NS) | |
| B2. Frequency in top – 4h | 2-way ANOVA, sex*injury | $F_{sex}$ (1,41) = 1.081, P=0.30 (NS)<br>$F_{injury}$ (1,41) = 11.93, P=0.0013<br>$F_{injury*sex}$ (1,41) = 0.442, P=0.51 (NS) | (injury effect presented on fig) |
| C1. Total duration climbing, timecourse | Mixed-effects model, sex*injury*time, covariate: baseline (SPSS, mixed model analysis, linear) | $F_{sex}$ (1,40.1) = 3.497, P=0.069 (NS)<br>$F_{injury}$ (1,39.7) = 4.847, P=0.034<br>$F_{time}$ (4,38.9) = 5.016, P=0.002<br>$F_{Covariate, BL}$ (1,39.0) = 9.062, P=0.005<br>$F_{sex*injury}$ (1,39.3)= 0.001, P=0.97 (NS)<br>$F_{time*injury}$ (4,36.599) = 1.144, P=0.351 (NS) | |
| C2. Total duration climbing – 4h | 2-way ANOVA, sex*injury | $F_{sex}$ (1,41) = 2.171, P=0.15 (NS)<br>$F_{injury}$ (1,41) = 4.817, P=0.034<br>$F_{injury*sex}$ (1,41) = 0.026, P=0.87 (NS) | (injury effect presented on fig) |
| D1. Distance – 4h, meloxicam | Unpaired T-test, One-tailed | T=0.8813, df = 19, P =0.1946 (NS) | |
| D2. Frequency in top – 4h, meloxicam | Unpaired T-test, one-tailed | T=1.559, df = 19, P =0.0678 (NS) | |
| D3. Total duration climbing – 4h, meloxicam | Unpaired T-test, one-tailed | T=0.9958, df = 19, P =0.1659 (NS) | |
| **Fig 2. CFA intraplantar** (Male: ctrl: n=6(5)*, CFA: n=6, Female: ctrl: n=6, CFA: n=6) (*one animal was excluded from the male control-group for the climbing data-set, due to the climbing exclusion-criteria at baseline.) | | | |
| A. Mechanical allodynia, vF. | RM ANCOVA, sex*injury, covariate: baseline, excl D11 (SPSS, general linear model) | $F_{time}$(2, 36) = 1.445, P=0.248 (NS)<br>$F_{time*injury}$(2, 36) = 0.033, P=0.97 (NS)<br>(Within-subjects effects, sphericity assumed/confirmed)<br><br>$F_{sex}$ (1,19) = 0.347, P=0.563 (NS)<br>$F_{injury}$ (1,19) = 219.1, P<0.001<br>$F_{sex*injury}$ (1,19)= 2.770, P=0.112 (NS)<br><br>$F_{covariate, BL}$ (1,19) = 5.403, P=0.031 | EMMs with Bonferroni:<br>D2: <0.001<br>D4: <0.001<br>D7: <0.001 |

| | | | |
|---|---|---|---|
| B. Distance moved. | RM ANCOVA, sex*injury, covariate: baseline, excl D11 (SPSS, general linear model) | $F_{time}(2, 36) = 4.248$, P=0.022<br>$F_{time*injury}(2, 36) = 2.229$, P=0.122 (NS)<br>(Within-subjects effects, sphericity assumed/confirmed)<br><br>$F_{sex}$ (1,18) = 2.964, P=0.102 (NS)<br>$F_{injury}$ (1,18) = 2.689, P=0.118 (NS)<br>$F_{sex*injury}$ (1,18)= 0.958, P=0.341 (NS)<br><br>$F_{covariate, BL}$ (1,18) = 7.181, P=0.015 | EMMs with Bonferroni:<br>D2: 0.034<br>D4: 0.388 (NS)<br>D7: 0.262 (NS) |
| C. Top climbing frequency | RM ANCOVA, sex*injury, covariate: baseline, excl D11 (SPSS, general linear model) | $F_{time}(2, 36) = 4.423$, P=0.019<br>$F_{time*injury}(2, 36) = 1.461$, P=0.245 (NS)<br>(Within-subjects effects, sphericity assumed/confirmed)<br><br>$F_{sex}$ (1,18) = 0.726, P=0.41 (NS)<br>$F_{injury}$ (1,18) = 7.098, P=0.016<br>$F_{sex*injury}$ (1,18)= 0.566, P=0.46 (NS)<br><br>$F_{covariate, BL}$ (1,18) = 12.846, P=0.002 | EMMs with Bonferroni:<br>D2: 0.006<br>D4: 0.081 (NS)<br>D7: 0.042 |
| D. Duration climbing | RM ANCOVA, sex*injury, covariate: baseline, excl D11 (SPSS, general linear model) | $F_{time}(1.4, 26.1) = 3.876$, P=0.046<br>$F_{time*injury}(1.4, 26.1) = 1.352$, P=0.269 (NS)<br>(Within-subjects effects, greenhouse-geisser)<br><br>$F_{sex}$ (1,18) = 3.134, P=0.113 (NS)<br>$F_{injury}$ (1,18) = 5.274, P=0.034<br>$F_{sex*injury}$ (1,18)= 1.839, P=0.192 (NS)<br><br>$F_{covariate, BL}$ (1,18) = 3.134, P=0.094 (NS) | EMMs with Bonferroni:<br>D2: 0.007<br>D4: 0.122 (NS)<br>D7: 0.207 (NS) |
| E1. Correlations, D2, Distance (%) * vF (%) | Simple linear regression, and Pearson's correlation analysis | r = 0.4589, F(1,21)=5.601, P=0.0276 | |
| E2. Correlations, D2, Top Frequency (%) * vF (%) | Simple linear regression, and Pearson's correlation analysis | r = 0.6145, F(1,21)=12.74, P=0.0018 | |
| E3. Correlations, D2, Duration (%) * vF (%) | Simple linear regression, and Pearson's correlation analysis | r = 0.5278, F(1,21)=8.108, P=0.0096 | |
| F1. Distance moved, D22, NTX, females | Unpaired T-test, one-tailed | T=1.019, df = 10, P =0.1660 (NS) | |
| F2. Top climbing frequency, D22, NTX, females | Unpaired T-test, one-tailed | T=1.813, df = 10, P =0.050 | |
| F3. Duration climbing, D22, NTX, females | Unpaired T-test, one-tailed | T=2.683, df = 10, P =0.0115 | |
| **Fig 3: Osteoarthritis (MIA)**<br>(ctrl-groups: n=8, MIA-groups: n=11*)<br>(*one male-MIA = excluded at baseline, one female-MIA = euthanized due to MIA toxicity) | | | |

| | | | |
|---|---|---|---|
| A1. DWB D8-9 | 2-way ANOVA, sex*injury | $F_{sex}$ (1,34) = 0.086, P=0.77 (NS)<br>$F_{injury}$ (1,34) = 209.8, P<0.0001<br>$F_{injury*sex}$ (1,34) = 0.003, P=0.97 (NS) | (ANOVA injury effect presented on fig) |
| A2. DWB D 15-16 | 2-way ANOVA, sex*injury | $F_{sex}$ (1,33) = 7.167, P=0.0115<br>$F_{injury}$ (1,33) = 26.58, P<0.0001<br>$F_{injury*sex}$ (1,33) = 0.014, P=0.91 (NS) | (ANOVA injury effect presented on fig) |
| A3. DWB D29-30 | 2-way ANOVA, sex*injury | $F_{sex}$ (1,34) = 0.964, P=0.333 (NS)<br>$F_{injury}$ (1,34) = 15.44, P=0.0004<br>$F_{injury*sex}$ (1,34) = 3.685, P=0.063 (NS) | (ANOVA injury effect presented on fig) |
| B. Distance moved. | RM ANCOVA, sex*injury, covariate: baseline (SPSS, general linear model) | $F_{time}$(7, 231) = 2.231, P=0.033<br>$F_{time*injury}$(7, 231) = 1.876, P=0.074 (NS)<br>(Within-subjects effects, sphericity assumed)<br><br>$F_{sex}$ (1,33) = 1.077, P=0.307 (NS)<br>$F_{injury}$ (1,33) = 5.125, P=0.030<br>$F_{sex*injury}$ (1,33)= 1.592, P=0.216 (NS)<br><br>$F_{covariate, BL}$ (1,33) = 33.968, P<0.001 | EMMs with Bonferroni:<br>D3: 0.068 (NS)<br>D7: *<br>D10: 0.080 (NS)<br>D14: NS<br>D20: NS<br>D28: 0.07 (NS)<br>D36: NS<br>D42: NS |
| C. Top climbing frequency | RM ANCOVA, sex*injury, covariate: baseline (SPSS, general linear model) | $F_{time}$(4.9, 164.5) = 2.345, P=0.044<br>$F_{time*injury}$(4.9, 164.5) = 1.924, P=0.093 (NS)<br>(Within-subjects effects, greenhouse-geisser)<br><br>$F_{sex}$ (1,33) = 0.911, P=0.347 (NS)<br>$F_{injury}$ (1,33) = 11.883, P=0.002<br>$F_{sex*injury}$ (1,33)= 0.650, P=0.426 (NS)<br><br>$F_{covariate, BL}$ (1,33) = 19.751, P<0.001 | EMMs with Bonferroni:<br>D3: **<br>D7: **<br>D10: *<br>D14: NS<br>D20: *<br>D28: **<br>D36: NS<br>D42: * |
| D. Duration climbing | RM ANCOVA, sex*injury, covariate: baseline (SPSS, general linear model) | $F_{time}$(7, 231) = 1.169, P=0.322 (NS)<br>$F_{time*injury}$(7, 231) = 2.57, P=0.014<br>(Within-subjects effects, sphericity assumed)<br><br>$F_{sex}$ (1,33) = 0.196, P=0.661 (NS)<br>$F_{injury}$ (1,33) = 9.480, P=0.004<br>$F_{sex*injury}$ (1,33)= 1.311, P=0.260 (NS)<br><br>$F_{covariate, BL}$ (1,33) = 27.193, P<0.001 | EMMs with Bonferroni:<br>D3: **<br>D7: ***<br>D10: **<br>D14: NS<br>D20: NS<br>D28: *<br>D36: NS<br>D42: 0.082 (NS) |
| E1. Correlations, D7, Distance * DWB ratio | Simple linear regression, and Pearson's correlation analysis | r = 0.4461, F(1,36) = 8.946, P=0.0050 | |
| E2. Correlations, D7, Top Frequency * DWB ratio | Simple linear regression, and Pearson's correlation analysis | r = 0.3969, F(1,36) = 6.732, P=0.0136 | |
| E3. Correlations, D7, Duration * DWB ratio | Simple linear regression, and Pearson's correlation analysis | r = 0.4610, F(1,36) = 9.717, P=0.0036 | |
| **Fig 4. Rheumatoid arthritis (CAIA)**<br>(Control: Fem: n=5, Male: n=4. Low: Fem: n=3, Male: n= 2. High: Fem: n=4, Male: n=5)<br>24 mice were excluded due to pre-set criteria for baseline climbing behaviour, and one animal was euthanized on day 6, due to predefined humane endpoints. | | | |
| A. Arthritis score | RM ANCOVA, sex*injury (SPSS, general linear model) | $F_{time}$(2.2, 37.7) = 18.564, P<0.001<br>$F_{time*injury}$(4.4, 37.7) = 5.665, P<0.001<br>(Within-subjects effects, greenhouse-geisser) | EMMs with Bonferroni:<br>D4: NS<br>D5: |

| | | | |
|---|---|---|---|
| | | $F_{sex}$ (1,17) = 0.037, P=0.85 (NS)<br>$F_{injury}$ (2,17) = 8.619, P=0.003<br>$F_{sex*injury}$ (2,17)= 0.151, P=0.861 (NS) | -ctrl vs high: *<br>-ctrl vs low: NS<br>D8:<br>-ctrl vs high: ***<br>-ctrl vs low: **<br>D14<br>-ctrl vs high: *<br>-ctrl vs low: 0.065 (NS) |
| B. Burrowing behaviour | RM ANCOVA, sex*injury, covariate: baseline 1 & 2 (SPSS, general linear model) | $F_{time}$(1, 15) = 0.261, P=0.617 (NS)<br>$F_{time*injury}$(2, 15) = 7.597, P=0.005<br>(Within-subjects effects, sphericity assumed)<br><br>$F_{sex}$ (1,15) = 0.124, P=0.730 (NS)<br>$F_{injury}$ (2,15) = 8.768, P=0.003<br>$F_{sex*injury}$ (2,15)= 0.212, P=0.812 (NS)<br><br>$F_{covariate, BL2}$ (1,15) = 0.391, P=0.541 (NS) | EMM's with Bonferroni:<br>D6:<br>-ctrl vs high: ***<br>-ctrl vs low: 0.052 (NS)<br>D13: NS |
| C. Distance Moved | RM ANCOVA, sex*injury, covariate: baseline (SPSS, general linear model) | $F_{time}$(1.4, 22.5) = 5.104, P=0.024<br>$F_{time*injury}$(2.8, 22.5) = 5.807, P=0.005<br>(Within-subjects effects, greenhouse-geisser)<br><br>$F_{sex}$ (1,16) = 0.3, P=0.591 (NS)<br>$F_{injury}$ (2,16) = 1.291, P=0.302 (NS)<br>$F_{sex*injury}$ (2,16)= 0.051, P=0.95 (NS)<br><br>$F_{covariate, BL}$ (1,16) = 6.920, P=0.018 | EMM's with Bonferroni:<br>D4:<br>-ctrl vs high: *<br>-ctrl vs low: *<br>D8: NS<br>D14: NS |
| D. Top climbing frequency | RM ANCOVA, sex*injury, covariate: baseline (SPSS, general linear model) | $F_{time}$(2, 32) = 1.475, P=0.244 (NS)<br>$F_{time*injury}$(4, 32) = 2.112, P=0.102 (NS)<br>(Within-subjects effects, sphericity assumed/confirmed)<br><br>$F_{sex}$ (1,16) = 1.438, P=0.248 (NS)<br>$F_{injury}$ (2,16) = 0.947, P=0.409 (NS)<br>$F_{sex*injury}$ (2,16)= 0.619, P=0.551 (NS)<br><br>$F_{covariate, BL}$ (1,16) = 1.51, P=0.237 (NS) | EMM's with Bonferroni:<br>D4: NS<br>D8: NS<br>D14: NS |
| E. climbing duration | RM ANCOVA, sex*injury, covariate: baseline (SPSS, general linear model) | $F_{time}$(2, 32) = 0.142, P=0.868 (NS)<br>$F_{time*injury}$(4, 32) = 2.573, P=0.057 (NS)<br>(Within-subjects effects, sphericity assumed/confirmed)<br><br>$F_{sex}$ (1,16) = 4.004, P=0.063 (NS)<br>$F_{injury}$ (2,16) = 2.983, P=0.079 (NS)<br>$F_{sex*injury}$ (2,16)= 2.504, P=0.113 (NS)<br><br>$F_{covariate, BL}$ (1,16) = 0.27, P=0.61 (NS) | EMM's with Bonferroni:<br>D4:<br>-ctrl vs high: **<br>-ctrl vs low: 0.098<br>D8: NS<br>D14: NS |
| F1. distance – burrowing correlation | Simple linear regression, and Pearson's correlation analysis | r = 0.7170, F(1,21)=22.22, P=0.0001 | |
| F2. Top frequency – burrowing correlation | Simple linear regression, and Pearson's correlation analysis | r = 0.5300, F(1,21)= 8.204, P=0.0093 | |
| F3. Total duration – burrowing correlation | Simple linear regression, and Pearson's correlation analysis | r = 0.6665, F(1,21)= 16.78, P=0.0005 | |
| **Fig 5. Cancer Induced Bone Pain** | | | |

| | | | |
|---|---|---|---|
| **Part 1: Breast CIBP**<br>(Only females, Sham: n=9 Cancer: n=7*)<br>(*2 mice were excluded due to tendon misplacement)) | | | |
| A. Breast cancer. Limb use | Only descriptive presentation. | | |
| B. Breast cancer. WB% | RM ANCOVA, injury, covariate: baseline (SPSS, general linear model) | $F_{time}(2.4, 31.3) = 0.531$, P=0.626 (NS)<br>$F_{time*injury}(2.4, 31.3) = 19.62$, P<0.001<br>(Within-subjects effects, greenhouse geisser)<br><br>$F_{injury}$ (1,13) = 32.103, P<0.001<br>$F_{covariate, BL}$ (1,13) = 0.114, P=0.74 (NS) | EMM's with Bonferroni:<br>D6: NS<br>D7: NS<br>D8:**<br>D9:**<br>D10:**<br>D11:**<br>D12:***<br>D13:***<br>D14:*** |
| C. Breast cancer. Distance moved | RM ANCOVA, injury, covariate: baseline (SPSS, general linear model) | $F_{time}(4,52) = 1.833$, P=0.137 (NS)<br>$F_{time*injury}(4,52) = 5.488$, P<0.001<br>(Within-subjects effects, sphericity assumed)<br><br>$F_{injury}$ (1,13) = 4.401, P=0.056 (NS)<br>$F_{covariate, BL}$ (1,13) = 2.715, P=0.123 (NS) | EMM's with Bonferroni:<br>D6: NS<br>D8: NS<br>D10: 0.064 (NS)<br>D12:*<br>D14: ** |
| D. Breast cancer - Top climbing frequency | RM ANCOVA, injury, covariate: baseline (SPSS, general linear model) | $F_{time}(4, 52) = 0.618$, P=0.65 (NS)<br>$F_{time*injury}(4,52) = 5.112$, P=0.002<br>(Within-subjects effects, sphericity assumed)<br><br>$F_{injury}$ (1,13) = 2.566, P=0.133 (NS)<br>$F_{covariate, BL}$ (1,13) = 3.080, P=0.103 (NS) | EMM's with Bonferroni:<br>D6: NS<br>D8: NS<br>D10: NS<br>D12: *<br>D14: * |
| E. Breast cancer – duration climbing | RM ANCOVA, injury, covariate: baseline (SPSS, general linear model) | $F_{time}(4,52) = 1.966$, P=0.113 (NS)<br>$F_{time*injury}(4,52) = 4.033$, P=0.006<br>(Within-subjects effects, sphericity assumed)<br><br>$F_{injury}$ (1,13) = 2.630, P=0.129 (NS)<br>$F_{covariate, BL}$ (1,13) = 0.708, P=0.42 (NS) | EMM's with Bonferroni:<br>D6: NS<br>D8: NS<br>D10: NS<br>D12: 0.064 (NS)<br>D14: * |
| F1. Correlation D14, Distance * WB | Simple linear regression, and Pearson's correlation analysis | r = 0.7178, F(1,14) = 14.8, P=0.0017 | |
| F2. Correlation D14, Top frequency * WB | Simple linear regression, and Pearson's correlation analysis | r = 0.6119, F(1,14) = 8.378, P=0.0118 | |
| F3. Correlation D14. Duration * WB | Simple linear regression, and Pearson's correlation analysis | r = 0.5834, F(1,14) = 7.224, P=0.0177 | |
| **Fig 5, part 2: Multiple Myeloma CIBP**<br>(Males only. Sham: n=7*, Cancer: n=9)<br>(* one animal was excluded due to baseline exclusion criteria for climbing, and one for tendon misplacement / error during surgery) | | | |
| G. Multiple Myeloma. Limb use | Only descriptive presentation. | | |
| H. Multiple Myeloma. | RM ANCOVA, injury, covariate: baseline | $F_{time}(3.39, 44.12) = 0.938$, P=0.44 (NS)<br>$F_{time*injury}(3.39, 44.12) = 3.477$, P=0.02 | EMM's with Bonferroni: |

| | | | |
|---|---|---|---|
| Distance moved | (SPSS, general linear model) | (Within-subjects effects, greenhouse geisser)<br><br>$F_{injury}$ (1,13) = 1.017, P=0.332 (NS)<br>$F_{covariate, BL}$ (1,13) = 6.297, P=0.026<br><br>D29-35:<br>$F_{injury}$ (1,13) = 4.80, P=0.047 | D8-25: NS<br>D29: 0.074 (NS)<br>D31: *<br>D33: 0.10 (NS)<br>D35: 0.10 (NS) |
| I. Multiple Myeloma - Top climbing frequency | RM ANCOVA, injury, covariate: baseline (SPSS, general linear model) | $F_{time}$(4.48, 58.2) = 0.521, P=0.74 (NS)<br>$F_{time*injury}$(4.48, 58.2) = 1.238, P=0.30 (NS)<br>(Within-subjects effects, greenhouse geisser)<br><br>$F_{injury}$ (1,13) = 0.614, P=0.447 (NS)<br>$F_{covariate, BL}$ (1,13) = 1.310, P=0.273 (NS)<br><br>D29-35:<br>$F_{injury}$ (1,13) = 1.423, P=0.254 (NS) | EMM's with Bonferroni:<br>D8-29: NS<br>D31: 0.087 (NS)<br>D33: NS<br>D35: NS |
| J. Multiple Myeloma – duration climbing | RM ANCOVA, injury, covariate: baseline (SPSS, general linear model) | $F_{time}$(10, 130) = 0.751, P=0.68 (NS)<br>$F_{time*injury}$(10, 130) = 0.969, P=0.47 (NS)<br>(Within-subjects effects, sphericity assumed/confirmed)<br><br>$F_{injury}$ (1,13) = 0.283, P=0.604 (NS)<br>$F_{covariate, BL}$ (1,13) = 0.018, P=0.895 (NS)<br><br>D29-35:<br>$F_{injury}$ (1,13) = 0.311, P=0.587 (NS) | EMM's with Bonferroni:<br>All NS |
| K1. Correlation, last measure. Distance * limb use | Simple linear regression, and Pearson's correlation analysis | r = 0.7026, F(1,14) = 13.65, P=0.0024 | |
| K2. Correlation, last measure, Top frequency * limb use | Simple linear regression, and Pearson's correlation analysis | r = 0.6023, F(1,14) = 7.970, P=0.0135 | |
| K3. Correlation, last measure. Duration * limb use | Simple linear regression, and Pearson's correlation analysis | r = 0.4916, F(1,14) = 4.462, P=0.0531 (NS) | |
| **Fig 6. Fibrous Dysplasia**<br>(males only, control: n=7, FD: n=5) | | | |
| A1.Home cage activity | RM ANCOVA, injury, covariate: baseline 2 (SPSS, general linear model) | $F_{time}$(23, 207) = 2.101, P=0.003<br>$F_{time*injury}$(23, 207) = 3.816, P=0.011<br>(Within-subjects effects, sphericity assumed/confirmed)<br><br>$F_{injury}$ (1,9) = 14.623, P=0.004<br>$F_{covariate, BL}$ (1,9) = 16.911, P=0.003 | EMM's with Bonferroni:<br>D8: **<br>D9: **<br>D10:**<br>D11:***<br>D12:**<br>D13:**<br>D14:***<br>D15:*<br>D16:0.51<br>D17: NS<br>D18: NS<br>D19: NS<br>D20: NS<br>D21: NS<br>D22: NS |

| | | | |
|---|---|---|---|
| | | | D23: NS |
| A2. AUC of activity | Unpaired t-test | T=4.755, df = 10, P =0.0008 | |
| B1. Running wheel distance | RM ANCOVA, injury, covariate: baseline 2 (SPSS, general linear model) | $F_{time}(23, 207) = 0.881$, P=0.624 (NS)<br>$F_{time*injury}(23, 207) = 6.102$, P<0.001<br>(Within-subjects effects, sphericity assumed/confirmed)<br><br>$F_{injury}$ (1,9) = 48.174, P<0.001<br>$F_{covariate, BL}$ (1,9) = 56.275, P<0.001 | EMM's with Bonferroni:<br>D6:*<br>D7: **<br>D8: ***<br>D9: ***<br>D10:***<br>D11:***<br>D12:***<br>D13:***<br>D14:***<br>D15:***<br>D16:***<br>D17: ***<br>D18: ***<br>D19: ***<br>D20: **<br>D21: **<br>D22: **<br>D23:0.06 |
| B2. AUC of running wheel | Unpaired t-test, with Welch's correction (unequal variance) | T=4.009, df = 6.276, P =0.0064 | |
| C. x-ray - Bone score | Only descriptive presentation. | | |
| D. Distance moved | RM ANCOVA, injury, covariate: baseline (SPSS, general linear model) | $F_{time}(1, 9) = 2.988$, P=0.118 (NS)<br>$F_{time*injury}(1, 9)$ = 18,532, P=0.002<br>(Within-subjects effects, greenhouse geisser)<br><br>$F_{injury}$ (1,9) = 22.652, P<0.001<br>$F_{covariate, BL}$ (1,9) = 14.439, P=0.004 | EMM's with Bonferroni:<br>D6: NS<br>D13: P<0.001 |
| E. Frequency in top | RM ANCOVA, injury, covariate: baseline (SPSS, general linear model) | $F_{time}(1, 9) = 0.67$, P=0.802 (NS)<br>$F_{time*injury}(1, 9) = 5.188$, P=0.049<br>(Within-subjects effects, greenhouse geisser)<br><br>$F_{injury}$ (1,9) = 7.178, P=0.025<br>$F_{covariate, BL}$ (1,9) = 1.890, P=0.202 (NS) | EMM's with Bonferroni:<br>D6: NS<br>D13: P=0.009 |
| F. Duration climbing | RM ANCOVA, injury, covariate: baseline (SPSS, general linear model) | $F_{time}(1, 9) = 0.501$, P=0.497 (NS)<br>$F_{time*injury}(1, 9) = 2.572$, P=0.143 (NS)<br>(Within-subjects effects, greenhouse geisser)<br><br>$F_{injury}$ (1,9) = 5.367, P=0.046<br>$F_{covariate, BL}$ (1,9) = 1.821, P=0.21 (NS) | EMM's with Bonferroni:<br>D6: NS<br>D13: P=0.033 |
| G1. Correlation, DVC locomotion – distance moved | Simple linear regression, and Pearson's correlation analysis | r = 0.6235, F(1,10) = 6.361, P=0.0303 | |
| G2. Correlation, DVC locomotion – top frequency | Simple linear regression, and Pearson's correlation analysis | r = 0.7404, F(1,10) = 12.13, P=0.0059 | |
| G1. Correlation, DVC | Simple linear regression, and | r = 0.5826, F(1,10) = 4.636, P=0.0568 (NS) | |

| | | | |
|---|---|---|---|
| locomotion – duration climbed | Pearson's correlation analysis | | |
| **Fig 7. Neuropathic pain**<br>A-D: males: sham: n=9, SNI: n=10, females: sham: n=7, SNI: n=9*<br>*one female was excluded from the "veh vs GBP" test due to clinical problems unrelated to the experiment<br>E: males only, control: n=10, SNI: n=12. | | | |
| A. Distance moved. | RM ANCOVA, sex*injury, covariate: baseline (SPSS, general linear model) | $F_{time}(4, 120) = 0.990$, P=0.416 (NS)<br>$F_{time*injury}(4, 120) = 0.890$, P=0.472 (NS)<br>(Within-subjects effects, sphericity assumed)<br><br>$F_{sex}$ (1,30) = 5.306, P=0.028<br>$F_{injury}$ (1,30) = 5.324, P=0.028<br>$F_{sex*injury}$ (1,30)= 0.016, P=0.902 (NS)<br><br>$F_{covariate, BL}$ (1,30) = 31.630, P<0.001 | EMMs with Bonferroni:<br>D4: *<br>D9: NS<br>D16: NS<br>D27/30-veh: *<br>D27/30-GBP: NS |
| B. Top climbing frequency | RM ANCOVA, sex*injury, covariate: baseline (SPSS, general linear model) | $F_{time}(3.2, 95.6) = 2.527$, P=0.059 (NS)<br>$F_{time*injury}(3.2, 95.6) = 1.081$, P=0.369 (NS)<br>(Within-subjects effects, greenhouse-geisser)<br><br>$F_{sex}$ (1,30) = 11.412, P=0.002<br>$F_{injury}$ (1,30) = 17.014, P<0.001<br>$F_{sex*injury}$ (1,30)= 2.521, P=0.123 (NS)<br><br>$F_{covariate, BL}$ (1,30) = 69.334, P<0.001 | EMMs with Bonferroni:<br>D4: **<br>D9: *<br>D16: **<br>D27/30-veh: **<br>D27/30-GBP: * |
| C. Duration climbing | RM ANCOVA, sex*injury, covariate: baseline (SPSS, general linear model) | $F_{time}(2.6, 78.2) = 0.275$, P=0.89 (NS)<br>$F_{time*injury}(2.6, 78.2) = 1.941$, P=0.138 (NS)<br>(Within-subjects effects, greenhouse-geisser)<br><br>$F_{sex}$ (1,30) = 0.167, P=0.206 (NS)<br>$F_{injury}$ (1,30) = 14.331, P<0.001<br>$F_{sex*injury}$ (1,30)= 0.469, P=0.499 (NS)<br><br>$F_{covariate, BL}$ (1,30) = 25.559, P<0.001 | EMMs with Bonferroni:<br>D4: *<br>D9: **<br>D16: **<br>D27/30-veh: ***<br>D27/30-GBP: * |
| D1. Distance – SNI only – GBP vs veh | 2-way ANOVA, sex*GBP | $F_{sex}$ (1,16) = 0.2678, P=0.6119 (NS)<br>$F_{GBP}$ (1,16) = 11.58, P=0.0036<br>$F_{GBP*sex}$ (1,16) = P=0.9965 (NS)<br><br>$F_{subject}$ (16,16) = 2.612, P=0.0317 | (injury effect presented on fig) |
| D2. Top climbing frequency – SNI only – GBP vs veh | 2-way RM ANOVA, sex*GBP | $F_{sex}$ (1,16) = 0.1734, P=0.6827 (NS)<br>$F_{GBP}$ (1,16) = 4.151, P=0.0585 (NS)<br>$F_{GBP*sex}$ (1,16) = 0.3296, P=0.5739 (NS)<br><br>$F_{subject}$ (16,16) = 4.333, P=0.0028 | (injury effect presented on fig) |
| D3. Total duration. SNI only – GBP vs veh | 2-way RM ANOVA, sex*GBP | $F_{sex}$ (1,16) = 0.2219, P=0.4692 (NS)<br>$F_{GBP}$ (1,16) = 5.743, P=0.0291<br>$F_{GBP*sex}$ (1,16) = 0.2219, P=0.64 (NS)<br><br>$F_{subject}$ (16,16) = 3.345, P=0.0104 | (injury effect presented on fig) |
| E1. Distance – naïve controls | RM ANCOVA, injury, covariate: baseline (SPSS, general linear model) | $F_{time}(2, 38) = 0.174$, P=0.841 (NS)<br>$F_{time*injury}(2, 38) = 0.339$, P=0.863 (NS)<br>(Within-subjects effects, sphericity assumed/confirmed)<br><br>$F_{injury}$ (1,19) = 6.054, P=0.024<br>$F_{covariate, BL}$ (1,19) = 4.046, P=0.059 (NS) | EMM's with Bonferroni:<br>D3: *<br>D7: *<br>D12: NS |
| E2. Top climbing frequency – naïve controls | RM ANCOVA, injury, covariate: baseline (SPSS, general linear model) | $F_{time}(2, 38) = 0.833$, P=0.443 (NS)<br>$F_{time*injury}(2, 38) = 0.166$, P=0.848 (NS)<br>(Within-subjects effects, sphericity assumed/confirmed) | EMM's with Bonferroni:<br>D3: ***<br>D7: ** |

| | | | |
|---|---|---|---|
| | | $F_{injury}$ (1,19) = 17.443, P<0.001<br>$F_{covariate, BL}$ (1,19) = 1.811, P=0.194 (NS) | D12: ** |
| E3. Total duration. – naïve controls | RM ANCOVA, injury, covariate: baseline (SPSS, general linear model) | $F_{time}$(2, 38) = 1.734, P=0.19 (NS)<br>$F_{time*injury}$(2, 38) = 1.119, P=0.337 (NS)<br>(Within-subjects effects, sphericity assumed/confirmed)<br><br>$F_{injury}$ (1,19) = 24.396, P<0.001<br>$F_{covariate, BL}$ (1,19) = 5.911, P=0.025 | EMM's with Bonferroni:<br>D3: ***<br>D7: ***<br>D12: ** |
| **Fig 8. Migraine**<br>(male: veh: n=8, LEV: n=9, female: veh: n=8, LEV: n=8) | | | |
| A. Grimace | 2-way ANOVA, sex*injury | $F_{sex}$ (1,36) = 3.574, P=0.0668 (NS)<br>$F_{injury}$ (1,36) = 11.97, P=0.0014<br>$F_{injury*sex}$ (1,36) = 0.7556, P=0.39 (NS) | (injury effect presented on fig) |
| B Top frequency | RM ANCOVA, injury, covariate: baseline (SPSS, general linear model) | $F_{time}$(3, 84) = 0.42, P=0.74 (NS)<br>$F_{time*injury}$(3, 84) = 0.958, P=0.41 (NS)<br>(Within-subjects effects, sphericity assumed)<br><br>$F_{injury}$ (1,28) = 2.694, P=0.112 (NS)<br>$F_{sex}$ (1,28) = 0.302, P=0.59 (NS)<br>$F_{covariate, BL}$ (1,28) = 3.078, P=0.09 (NS) | EMM's with Bonferroni:<br>Across sex:<br>D1-1h: P=0.028<br>D1-2h: NS<br>D5-1h: P=0.072 (NS)<br>D5-2h: NS |
| C Duration climbing | RM ANCOVA, injury, covariate: baseline (SPSS, general linear model) | $F_{time}$(3, 84) = 1.363, P=0.26 (NS)<br>$F_{time*injury}$(3, 84) = 3.812, P=0.013<br>(Within-subjects effects, sphericity assumed)<br><br>$F_{injury}$ (1,28) = 8.461, P=0.007<br>$F_{sex}$ (1,28) = 0.202, P=0.66 (NS)<br>$F_{covariate, BL}$ (1,28) = 5.281, P=0.029 | EMMs with Bonferroni:<br>Across sex:<br>D0-1h: P<0.001<br>D0-2h: NS<br>D4-1h: P=0.017<br>D4-2h: NS |
| D1 Correlation, grimace delta – top frequency | Simple linear regression, and Pearson's correlation analysis | r = -0.4605, F(1,30) = 8.072, P=0.0080 | |
| D2 Correlation, grimace delta – Duration climbing | Simple linear regression, and Pearson's correlation analysis | r = -0.4779, F(1,30) = 8.879, P=0.0057 | |
| **Fig 9. FRICT-ION** | | | |
| A. vF | RM ANCOVA, injury, covariate: baseline (SPSS, general linear model) | $F_{time}$(1.3, 26.2) = 0.805, P=0.041 (NS)<br>$F_{time*injury}$(1.3, 26.2) = 0.264, P=0.32 (NS)<br>(Within-subjects effects, sphericity violated, greenhouse-geisser corrected)<br><br>$F_{injury}$ (1,20) = 16.333, P<0.001<br>$F_{sex}$ (1,20) = 1.0, P = 0.33 (NS)<br>$F_{sex*injury}$ (1,20) = 0.21, P=0.652 (NS)<br>$F_{covariate, BL}$ (1,20) = 0.013, P=0.91 (NS) | |
| B. ADT | RM ANCOVA, injury, covariate: baseline (SPSS, general linear model) | $F_{time}$(2, 40) = 0.119, P = 0.89 (NS)<br>$F_{time*injury}$(2, 40) = 1.343, P = 0.27 (NS)<br>(Within-subjects effects, sphericity assumed)<br><br>$F_{injury}$ (1,20) = 124.048, P < 0.001<br>$F_{sex}$ (1,20) = 3.195, P=0.089 (NS)<br>$F_{sex*injury}$ (1,20) = 0.012, P=0.915(NS)<br>$F_{covariate, BL}$ (1,20) = 2.368, P=0.14 (NS) | |

| | | | |
|---|---|---|---|
| C. Distance moved | RM ANCOVA, injury, covariate: baseline (SPSS, general linear model) | $F_{time}(3, 60) = 2.704$, P=0.053 (NS)<br>$F_{time*injury}(3, 60) = 0.357$, P=0.785 (NS)<br>(Within-subjects effects, sphericity assumed)<br><br>$F_{injury}$ (1,20) = 0.565, P=0.461 (NS)<br>$F_{sex}$ (1,20) = 0.199, P=0.66 (NS)<br>$F_{sex*injury}$ (1,20) = 2.661, P=0.118<br>$F_{covariate, BL}$ (1,20) = 13.225, P=0.002 | |
| D. Top Frequency | RM ANCOVA, injury, covariate: baseline (SPSS, general linear model) | $F_{time}(3, 60) = 1.729$, P=0.17 (NS)<br>$F_{time*injury}(3, 60) = 0.471$, P=0.70 (NS)<br>(Within-subjects effects, sphericity assumed)<br><br>$F_{injury}$ (1,20) = 5.027, P=0.036<br>$F_{sex}$ (1,20) = 0.18, P=0.68 (NS)<br>$F_{sex*injury}$ (1,20) = 178, P=0.69<br>$F_{covariate, BL}$ (1,20) = 11.701, P=0.003 | |
| E. Duration climbing | RM ANCOVA, injury, covariate: baseline (SPSS, general linear model) | $F_{time}(3, 60) = 0.299$, P=0.83 (NS)<br>$F_{time*injury}(3, 60) = 0.269$, P=0.85 (NS)<br>(Within-subjects effects, sphericity assumed)<br><br>$F_{injury}$ (1,20) = 4.662, P=0.043<br>$F_{sex}$ (1,20) = 0.262, P=0.61 (NS)<br>$F_{sex*injury}$ (1,20) = 0.001, P=0.98 (NS)<br>$F_{covariate, BL}$ (1,20) = 6.568, P=0.019 | |
| F1. Correlations, D14, Distance (%) * vF (%) | Simple linear regression, and Pearson's correlation analysis | r = 0.1700, F(1,23) = 0.6846, P=0.4165 (NS) | |
| F2. Correlations, D14, Top Frequency * vF (g) | Simple linear regression, and Pearson's correlation analysis | r = 0.3203, F(1,23) = 2.629, P=0.1186 (NS) | |
| F3. Correlations, D14, Duration (s) * vF (g) | Simple linear regression, and Pearson's correlation analysis | r = 0.4027, F(1,23)= 4.452, P=0.0459 | |
| **Fig 10. Comparisons** | | | |
| B. Baseline comparisons – climbing duration. | 2way ANOVA, sex*experiment | $F_{experiment}$ (10,277) = 15.73, P<0.0001<br>$F_{sex}$ (1,277) = 0.8243, P=0.36 (NS) | Tukey post hoc.<br>CFA vs MIA: *<br>CFA vs LEV: *<br>CFA vs MM: ***<br>SNI1 vs SNI2: ***<br>SNI1 vs MIA: ***<br>SNI1 vs MM: ***<br>SNI1 vs BCIBP: ***<br>SNI2 vs ION: *<br>SNI2 vs Incis: ***<br>SNI2 vs LEV: ***<br>SNI2 vs MM: *<br>SNI2 vs CAIA: **<br>ION vs MIA: **<br>ION vs MM: ***<br>ION vs: BCIBP: *<br>Incis vs MIA: ***<br>Incis vs MM: ***<br>Incis vs BCIBP: ***<br>MIA vs FD: *<br>MIA vs MM: *<br>MIA vs CAIA: ***<br>LEV vs MM: ***<br>LEV vs BCIBP: ***<br>FD vs MM: ***<br>MM vs CAIA: ***<br>B-CIBP vs CAIA: ** |

| | | |
|---|---|---|
| C: baselines, oestrus cycle. | One way ANOVA, cycle stage | F (3, 25) = 0.4208, P = 0.7396 |
| D: | Descriptive | |

NS = Non significant. *: P<0.05, **: P<0.01, ***:P<0.001.

### Table S4: Overview of model-specific results.

| | ***Baseline, covariate*** | | |
|---|---|---|---|
| ***Model*** | **Distance** | **Top frequency** | **Duration** |
| *Incision* | P = 0.031* | P = 0.055 (NS) | P = 0.005* |
| *CFA* | P = 0.015* | P = 0.002* | P = 0.094 (NS) |
| *MIA* | P < 0.001* | P < 0.001* | P < 0.001* |
| *CAIA* | P = 0.018* | NS | NS |
| *Breast CIBP* | NS | NS | NS |
| *MM CIBP* | P=0.026* | NS | NS |
| *FD* | P = 0.004* | NS | NS |
| *SNI-1* | P<0.001* | P<0.001* | P<0.001* |
| *SNI-2* | P = 0.059 (NS) | NS | P=0.025* |
| *LEV* | N/A | P=0.09(NS) | P = 0.029* |
| *FRICT-ION* | P = 0.002* | P = 0.003* | P = 0.019* |

**Table 1:** Summarized data from statistical analysis, presented in full in **supplementary Table S3**. All data was analysed using RM ANCOVA (or mixed effects analysis), factors: sex*injury*time, covariate = baseline. For datasets without significant injury-effects, but significant time*injury interactions, the particular days of significant differences between control and injury group are mentioned, as estimated by Estimated Marginal Means with Bonferroni correction. For results below P=0.1, the P-value was specified, while for P>0.1, they were simply stated: NS = non-significant. *: P<0.05. Models: Incision: plantar incisions model, CFA: intraplantar Complete Freund's Adjuvant, MIA: Monoiodoacetate Induced Arthritis, CAIA: Collagen Antibody Induced Arthritis, Breast CIBP: Breast Cancer-Induced Bone Pain, MM CIBP: Multiple Myeloma Cancer-Induced Bone Pain, FD: Fibrous Dysplasia, SNI: Spared Nerve Injury (SNI-1: male-female experiment, SNI-2: male-control experiment), LEV: Levcromacalim-induced migraine, FRICT-ION: Foramen Rotundum Inflammatory Constriction Trigeminal Infraorbital Nerve injury.

**Table S5. Author details, conflicts of interest and funding sources.**

| Name | Email | Conflict of interest | Funding | Orcid |
|---|---|---|---|---|
| Morten E. Malmberg | | Nothing to declare | NA (UCPH funded) | 0009-0000-9849-080X |
| Tobias Bro | | Nothing to declare | NA | 0009-0009-9404-8243 |
| Edward Ciputra | | Nothing to declare | Funded by the European Union. BonePainIII Grant No. 101120524 | |
| Pernille Colding-Jørgensen | | Nothing to declare | CALAR | 0000-0002-0163-1790 |
| Jannie Jutta Jeppesen | | Nothing to declare | NA | |
| Carolyn Marie Goddard | | Nothing to declare | Lundbeck Foundation grant R389-2021-1596, Neuroscience Academy Denmark | 0009-0000-5356-1336 |
| Ida Buur Kanneworff | | Employee of Zyneyro, a company developing nonopioid drugs for chronic pain; the present study is independent of and unrelated to Zyneyro's research and development activities. | Innovation Fund Denmark (Industrial PhD 3194-00050B) | 0009-0002-5697-1271 |
| Heather Allen | | Nothing to declare. | NA | |
| Kathrine Louise Jensen | | Employee of Zyneyro, a company developing nonopioid drugs for chronic pain; the present study is independent of and unrelated to Zyneyro's research and development activities. | NA | 0000-0001-8985-9126 |
| Helene Dragelund Garcia | | Nothing to declare | NA | 0000-0003-1303-1140 |
| Yingyu Tang | | Nothing to declare | Funded by the European Union. BonePainIII Grant No. 101120524 and Fonden til Lægevidenskabens Fremme | 0009-0007-3490-6694 |

| | | | | |
|---|---|---|---|---|
| Kholud Walid El Masri | | Nothing to declare | CALAR | 0009-0005-6876-1687 |
| Andrea Soeberg | | Nothing to declare | NA | |
| Astrid Thostrup Davidsen | | Nothing to declare | NA | |
| Naomi K. Grabus | | Nothing to declare. | NA | |
| Anja Holm | | Nothing to declare. | NA (TRACE funded) | 0000-0001-8796-1872 |
| Sarah Louise Christensen | | Nothing to declare. | BRIDGE - Translational Excellence Programme<br>Novo Nordisk Foundation (NNF23SA0087869). | 0000-0002-7985-4783 |
| Karina Obelitz-Ryom | | Nothing to declare. | Independent Research Fund Denmark (3162-00044B) | 0000-0002-2771-4842 |
| Chelsea Hopkins | | Nothing to declare. | Carlsberg Foundation, grant CF23-1620<br>Imaging data were collected at the Center for Advanced Bioimaging (CAB) Denmark, University of Copenhagen, which is operated with funding from the Novo Nordisk Foundation (NNF23OC0082200). | 0000-0003-1320-7644 |
| Anne-Marie Heegaard | | Nothing to declare. | Funded by the European Union. BonePainIII Grant No. 101120524 | 0000-0003-2754-0800 |
| Kenneth Lindegaard Madsen | | Cofounder of Zyneyro, a company developing nonopioid drugs for chronic pain; the present study is independent of and unrelated to Zyneyro's research and development activities. | Funded by Lundbeckfonden Ascending Investigator (LBF R467-2024-544) | 0000-0001-9274-6691 |
| Andreas Toft Sørensen | | Cofounder of Zyneyro, a company developing nonopioid drugs for chronic pain; the present study is independent of and unrelated | NA | 0000-0001-5887-9247 |

| | | | | |
|---|---|---|---|---|
| | | to Zyneyro's research and development activities. | | |
| Rajesh Khanna | | Cofounder of Regulonix LLC, a company developing nonopioids drugs for chronic pain: the present study is independent and unrelated to Regulonix's research and development activities. | Startup funds from the University of Florida. | 0000-0002-9066-2969 |
| Klas S.P. Abelson | | Nothing to declare | NA | 0000-0002-4986-7882 |
| Otto Kalliokoski | | Nothing to declare | NA | 0000-0001-6531-3487 |
| Sara Hestehave | | Nothing to declare | NA | 0000-0002-1837-521X |

**Abbreviations:** CALAR = Center for Applied Laboratory Animal Research, DK. NA = Not Applicable.

**Table S6. Authorship Contribution – specified.**

| **Name** | **Conceptualization** | **Designed experiments** | **Conducting** | **Analysis** | **Drafting manuscript** | **Review and editing** | **Supervising** |
|---|---|---|---|---|---|---|---|
| M. E. Malmberg | Incision-studies + climbing device | Incision studies | Incision studies | | Incision-methods | Yes | |
| T. Bro | Incision-studies + climbing device | Incision studies | Incision studies | Incision studies | | yes | |
| E. Ciputra | CIBP studies | CIBP studies | CIBP studies | CIBP studies | CIBP methods | Yes | |
| P. Colding-Jørgensen | | CAIA study | CAIA study | | | Yes | |
| J.J. Jeppesen | | LEV studies | LEV studies | LEV studies | LEV methods | Yes | |
| C.M. Goddard | | CFA study | CFA study | | | Yes | |
| I.B. Kanneworff | | SNI studies | SNI studies | | | Yes | |
| H.N. Allen | FRICT-ION | FRICT-ION | FRICT-ION | FRICT-ION | FRICT-ION methods | Yes | |

| | | | | | | | |
|---|---|---|---|---|---|---|---|
| N.K. Grabus | | FRICT-ION | FRICT-ION | FRICT-ION | FRICT-ION methods | Yes | |
| K.L. Jensen | | SNI studies | SNI studies | | SNI methods | Yes | |
| H.D. Garcia | | MIA study | MIA studies | | | Yes | |
| Y. Tang | | FD studies | FD studies | FD studies | FD methods | Yes | |
| K.W. E. Masri | | | CAIA study | CAIA study | | Yes | |
| A.Soeberg | | | FD studies | FD studies | | Yes | |
| A.T. Davidsen | | | CIBP studies | CIBP studies | | Yes | |
| A. Holm | | | | LEV studies | | Yes | |
| S.L. Christensen | LEV studies | | | | | Yes | LEV studies |
| K. Obelitz-Ryom | LEV studies | LEV studies | | LEV studies | LEV methods | Yes | LEV studies |
| C. Hopkins | CIBP + FD studies | CIBP + FD studies | | | | Yes | |
| AM. Heegaard | CIBP + FD studies | CIBP + FD studies | | | | yes | CIBP + FD studies |
| K.L. Madsen | CFA + SNI study | | | | | Yes | SNI + CFA studies |
| A.T. Sørensen | CFA study | | | | | Yes | SNI + CFA studies |
| R. Khanna | | | | | | Yes | FRICT-ION |
| K.S.P. Abelson | Incision studies | | | | | YES | Incision + CAIA studies |
| O. Kalliokoski | | | | | | Yes | Incision studies and general statistical analysis |

| | | | | | | | |
|---|---|---|---|---|---|---|---|
| S. Hestehave | All model specific elements, project coordination and development of climbing device | Incision, MIA, CAIA, SNI, CFA, | MIA studies | Incision, SNI, CFA, MIA, CAIA, FRICT-ION | Merged methods, drafted figures and performed statistical analysis. Drafted remaining manuscript | Yes | Supervised and coordinated the entire project incl. individual experiments. |

Abbreviations: FD = Fibrous Dysplasia. CIBP = Cancer Induced Bone Pain models. Incision = Postoperative Plantar incision model. SNI = Spared Nerve Injury, CFA = intraplantar injection of Complete Freund's Adjuvant, MIA = Monoiodoacetate Induced Arthritis, CAIA = Collagen Antibody Induced Arthritis, LEV = Levcromakalim. FRICT-ION = Foramen Rotundum Inflammatory Constriction Trigeminal Infraorbital Nerve injury.

[10] Hestehave S, Allen HN, Gomez K, Duran P, Calderon-Rivera A, Loya-López S, Rodríguez-Palma EJ, Khanna R. Small molecule targeting NaV1.7 via inhibition of

CRMP2-Ubc9 interaction reduces pain-related outcomes in a rodent osteoarthritic model. PAIN 2025;166(1):99-111.

[22] Parkes I, Schorscher-Petcu A, Gan Q, Browne LE. Precision cutaneous stimulation in freely moving mice: eLife Sciences Publications, Ltd, 2026.

[23] Rudnik-Jansen I, Schrijver K, Woike N, Tellegen A, Versteeg S, Emans P, Mihov G, Thies J, Eijkelkamp N, Tryfonidou M, Creemers L. Intra-articular injection of triamcinolone

acetonide releasing biomaterial microspheres inhibits pain and inflammation in an acute arthritis model. Drug delivery 2019;26(1):226-236.
[24] Rudnik-Jansen I, Woike N, de Jong S, Versteeg S, Kik M, Emans P, Mihov G, Thies J, Eijkelkamp N, Tryfonidou M, Creemers L. Applicability of a Modified Rat Model of Acute Arthritis for Long-Term Testing of Drug Delivery Systems. Pharmaceutics 2019;11(2).
[25] Santos EJ, Giddings AN, Kandil FA, Negus SS. Climbing behavior by mice as an endpoint for preclinical assessment of drug effects in the absence and presence of pain. Frontiers in Pain Research 2023;4.
[26] Scholz J, Broom DC, Youn DH, Mills CD, Kohno T, Suter MR, Moore KA, Decosterd I, Coggeshall RE, Woolf CJ. Blocking caspase activity prevents transsynaptic neuronal apoptosis and the loss of inhibition in lamina II of the dorsal horn after peripheral nerve injury. J Neurosci 2005;25(32):7317-7323.
[27] Sliepen SHJ, Diaz-Delcastillo M, Korioth J, Olsen RB, Appel CK, Christoph T, Heegaard AM, Rutten K. Cancer-induced Bone Pain Impairs Burrowing Behaviour in Mouse and Rat. In Vivo 2019;33(4):1125-1132.
[28] Wodarski R, Delaney A, Ultenius C, Morland R, Andrews N, Baastrup C, Bryden LA, Caspani O, Christoph T, Gardiner NJ, Huang W, Kennedy JD, Koyama S, Li D, Ligocki M, Lindsten A, Machin I, Pekcec A, Robens A, Rotariu SM, VoB S, Segerdahl M, Stenfors C, Svensson CI, Treede R-D, Uto K, Yamamoto K, Rutten K, Rice ASC. Cross-centre replication of suppressed burrowing behaviour as an ethologically relevant pain outcome measure in the rat: a prospective multicentre study. Pain 2016;157(10):2350-2365.
[29] Zhao X, Deng P, Iglesias-Bartolome R, Amornphimoltham P, Steffen DJ, Jin Y, Molinolo AA, de Castro LF, Ovejero D, Yuan Q, Chen Q, Han X, Bai D, Taylor SS, Yang Y, Collins MT, Gutkind JS. Expression of an active Gα(s) mutant in skeletal stem cells is sufficient and necessary for fibrous dysplasia initiation and maintenance. Proc Natl Acad Sci U S A 2018;115(3):E428-e437.
